# Base-editing a single missense mutation in A20 enhances CAR-T cell efficacy

**DOI:** 10.1101/2025.10.05.680562

**Authors:** Adam Blaisdell, Stefanie Bachl, Luis R. Sandoval, Carter Ching, Christopher J. Bowman, Nupura Kale, Manu Prabandham, Morgan Diolaiti, Claire Havig, Rommel Advincula, Nika Lenci, Zhongmei Li, Emily Yamashita, Charlotte H. Wang, Shimin Zhang, Qi Liu, Philip Achacoso, Dorothea Stibor, Inger Øynebråten, Jin Seo, Alan Ashworth, Alexander Marson, Chun Jimmie Ye, Barbara A. Malynn, Justin Eyquem, Julia Carnevale, Averil Ma

**Author notes:** These authors contributed equally (Adam Blaisdell, Stefanie Bachl, Luis R. Sandoval).

## Abstract

T cell exhaustion limits the efficacy of cancer immunotherapies. Here, we performed genome-wide loss-of-function screening in repetitively stimulated human T cells and identified the mulitfunctional ubiquitin-modifying protein A20/TNFAIP3 as a major negative regulator of exhausted T cell persistence. Protein large language modeling, deep base-editing mutagenesis, and studies in immunocompetent mice with domain-specific inactivating mutations revealed A20’s non-enzymatic M1 ubiquitin-binding zinc finger 7 (A20^ZF7^) motif as critical to suppression of anti-tumor immunity. A20^ZF7^-deficient CD8^+^ tumor-infiltrating lymphocytes (TILs) resisted terminal exhaustion and circumvented an unappreciated mechanism restraining perforin degranulation in terminally exhausted cells. Human chimeric antigen receptor (CAR)-T cells engineered via base-editing to inactivate A20^ZF7^ via a single missense mutation also resisted exhaustion, secreted more perforin and robustly suppressed cancer *in vivo*. These studies pinpoint A20^ZF7^ as a novel T cell checkpoint and reveal precision base-editing of missense mutations as an effective approach to enhance CAR-T cell therapy.

## Introduction

Tumors induce an exhausted state in tumor-infiltrating lymphocytes (TILs) and CAR-T cells alike, characterized by increased expression of inhibitory receptors (IRs), diminished tumoricidal capacity, and reduced cytokine production^1^. Nevertheless, exhausted CD8 T cells (Tex) continue to express perforin and granzymes^2–4^, suggesting that cytokines and cytotoxic granule proteins are regulated differently in Tex, and hinting at posttranscriptional restraint of cytotoxic potential. However, the ability of Tex to degranulate *in situ* within the tumor microenvironment (TME) has yet to be evaluated.

Identification of targets that can unlock the cytotoxic potential of Tex could markedly enhance cellular therapies. Repetitive *in vitro* T cell receptor (TCR) stimulation can mimic the chronic antigenic stimulation that drives T cell exhaustion, and when combined with CRISPR/Cas9 knockout screens can unveil key regulators of this process. To date, genome-wide screens in human T cells have utilized assays with a short course of one or two tumor cell co-cultures for repetitive antigenic stimulation^5,6^. However, a longer course of repeated tumor cell co-cultures could more closely model TIL experience in the TME, and uncover new therapeutic targets relevant to later stages of Tex differentiation.

In addition to Cas9-directed gene ablation, base-editors (BEs) have emerged as an efficient method to interrogate how individual amino acid changes affect T cell behaviors^7,8^. While BEs can ablate genes via nonsense mutations, their unique power lies in the precise tuning of protein function via hyper- or hypomorphic missense mutations. For multifunctional proteins, this approach could specifically tune one function while preserving others. In the context of engineering next-generation adoptive cellular therapies (ACTs) such as TCR-T or CAR-T cells, BE-directed missense mutations could prove more efficacious and potentially safer. While large-scale BE screens have singled out missense mutations that improve CAR-T cell function *in vitro*^7,8^, this approach has to our knowledge yet to boost efficacy *in vivo*.

A20/TNFAIP3 is a multifunctional ubiquitin (Ub)-modifying protein linked to autoimmune diseases^9^. A20 contains enzymatic and non-enzymatic domains with distinct functions that cooperate to regulate NF-κB and other Ub-dependent pathways^9^. Complete A20 deficiency in mice leads to systemic inflammation and perinatal lethality^10^, while knock-in (KI) mice with A20 domain-specific hypomorphic mutations survive long into adulthood and exhibit variable degrees of spontaneous or inducible inflammation^11–13^. Complete or hypomorphic A20 deficiency within T cells can augment primary TCR responses but can also lead to cell death and impaired memory response to infection^14–18^. However, it remains unknown whether A20 or its hypomorphs impact T cell exhaustion, or whether A20 represents a suitable target for ACTs. Prior studies have implicated A20 as a suppressor of CD8 TIL persistence in the TME^19^, as well as CD8 TIL cytotoxicity and effector cytokine (IFNγ and TNF) production^20–22^, though the relevant anti-tumor effector mechanism was never directly tested. Notably, the requirement for specific effector cytokines versus cytotoxic granule proteins to suppress tumors can vary widely across models^23–30^, and thus identifying targets that can enhance CD8 TIL cytotoxicity in multiple ways could lead to broader therapeutic applications.

Here, we use a genome-scale CRISPR knockout screen with a long duration of repetitive antigenic stimulation, a protein large language model for predicting missense variant effects, a BE tiling screen of the A20 gene, and a panel of A20 KI mice to unveil A20’s non-enzymatic zinc finger 7 (A20^ZF7^) domain as a potent instructor of T cell exhaustion. Through the lens of A20 deficiency, we discover restrained perforin degranulation as an underrecognized post-transcriptional regulatory feature of Tex biology that is sharply reversed by abrogating the A20^ZF7^ motif. Importantly, this enhanced degranulation, rather than increased effector cytokine production, provides the critical anti-tumor effector mechanism in A20^ZF7^-deficient Tex. Finally, we show that a BE-guided missense mutation to inactivate A20^ZF7^ substantially improves the efficacy of human CAR-T cells both *in vitro* and *in vivo*.

## Results

### A genome-wide CRISPR perturbation screen identifies A20/*TNFAIP3* as a key regulator of T cell dysfunction upon chronic antigen stimulation

The ability to repeatedly eradicate tumor cells during sustained tumor antigen exposure is an indispensable characteristic of therapeutic T cells. To reveal genes that have the potential to enhance T cell persistence and deter dysfunction, we employed an *in vitro* system that models chronic exposure to tumor antigen^31,32^. We engineered human T cells to express the NY-ESO-1 specific TCR (1G4) and repeatedly co-cultured them with fresh NY-ESO-1^+^ A375 melanoma cells (**Supp. Fig. 1A**). After multiple rounds of tumor exposure, we assessed the ability of TCR-T cells to eradicate tumor cells *in vitro*. TCR-T cells that were continuously exposed to tumor antigen displayed a substantial decline in their capacity to control tumor cell growth after each successive round of tumor stimulation, while TCR-T cells matched for time in culture, without prior exposure to antigen, maintained robust tumor-killing capability (**Supp. Fig. 1B**). Additionally, cell surface inhibitory receptors (IRs) whose expression are associated with T cell exhaustion (LAG-3, TIM-3, PD-1, and CD39) increased as TCR-T cells were repeatedly exposed to tumor cells, with the difference between tumor-exposed and unexposed TCR-T cells growing progressively over successive rounds of stimulation (**Supp. Fig. 1C**). Finally, the capacity of TCR-T cells to secrete effector cytokines (IFNγ, TNF) and cytotoxic granule proteins (perforin and granzyme B [GzmB]) decreased after several rounds of exposure to tumor cells (**Supp. Fig. 1C**), providing further evidence of dysfunction from chronic antigen stimulation. Together these observations provide evidence that the chronic antigen stimulation we used models key aspects of TIL dysfunction.

To nominate gene modifications that improve TIL persistence and deter dysfunction in the TME, we performed a genome-wide CRISPR/Cas9 knockout screen using this repetitive tumor stimulation system^33^. Primary human T cells from two human donors were transduced with the 1G4 TCR along with a genome-wide single-guide RNA (sgRNA) library^34^, and Cas9 protein was introduced by electroporation^31,33^. We then exposed pooled Cas9-edited TCR-T cells to nine successive rounds of tumor cell stimulation in the presence of IL-2 (**Fig. 1A**), which greatly exceeds the number of repeated tumor exposures in prior genome-wide screens in human T cells^5,6^. We maintained an effector to target (E:T) ratio of 1:1 for each stimulation until the bulk TCR-T cells could no longer effectively eliminate the tumor cells, indicating a reduction in their functional capacity. Pooled edited TCR-T cells were sequenced either before initial antigenic exposure (Time-Zero; T0) or after 4 rounds (Time-Intermediate; TI) or 9 rounds (Time-Final; TF) of antigenic stimulation. A parallel set of edited TCR-T cells was cultured for the duration of the screen with IL-2 but not exposed to tumor cells (TF_ctrl_). We identified numerous sgRNAs enriched in TF when compared with T0 (**Fig. 1B**; Supp. Tables 1-2), including sgRNAs targeting genes with established roles in T cell fitness and exhaustion (*RASA2*, *CISH*, *DNMT3A*, *MAPK14*, and the Cullin-5 complex members *CUL5* and *RNF7*)^31,35–38^, as well as sgRNAs targeting subunits of Mediator (*MED15*, *MED16*, and *MED24*) and SWI/SNF (*SMARCD1*) — protein complexes highlighted by prior chronic stimulation screens^6,39^. Intriguingly, we also discovered in our chronic stimulation screen strong enrichment for A20/*TNFAIP3* sgRNAs (rank 4; **Fig. 1B**), a target that was not enriched in these prior screens. A20 sgRNAs were also highly correlated between T cell donors (**Supp. Fig. 1D**). When we compared TF_ctrl_ with T0, A20 was ranked 4718th (**Supp. Fig. 1E**; Supp. Table 3), suggesting that the high rank in TF compared with T0 was specific to repetitive stimulation. Comparison of TI with T0 showed A20 was ranked 70th (**Supp. Fig. 1E**; Supp. Tables 4-5), indicating a relative increase in the A20 persistence advantage with increasing chronic stimulation pressure.

**Fig. 1.**
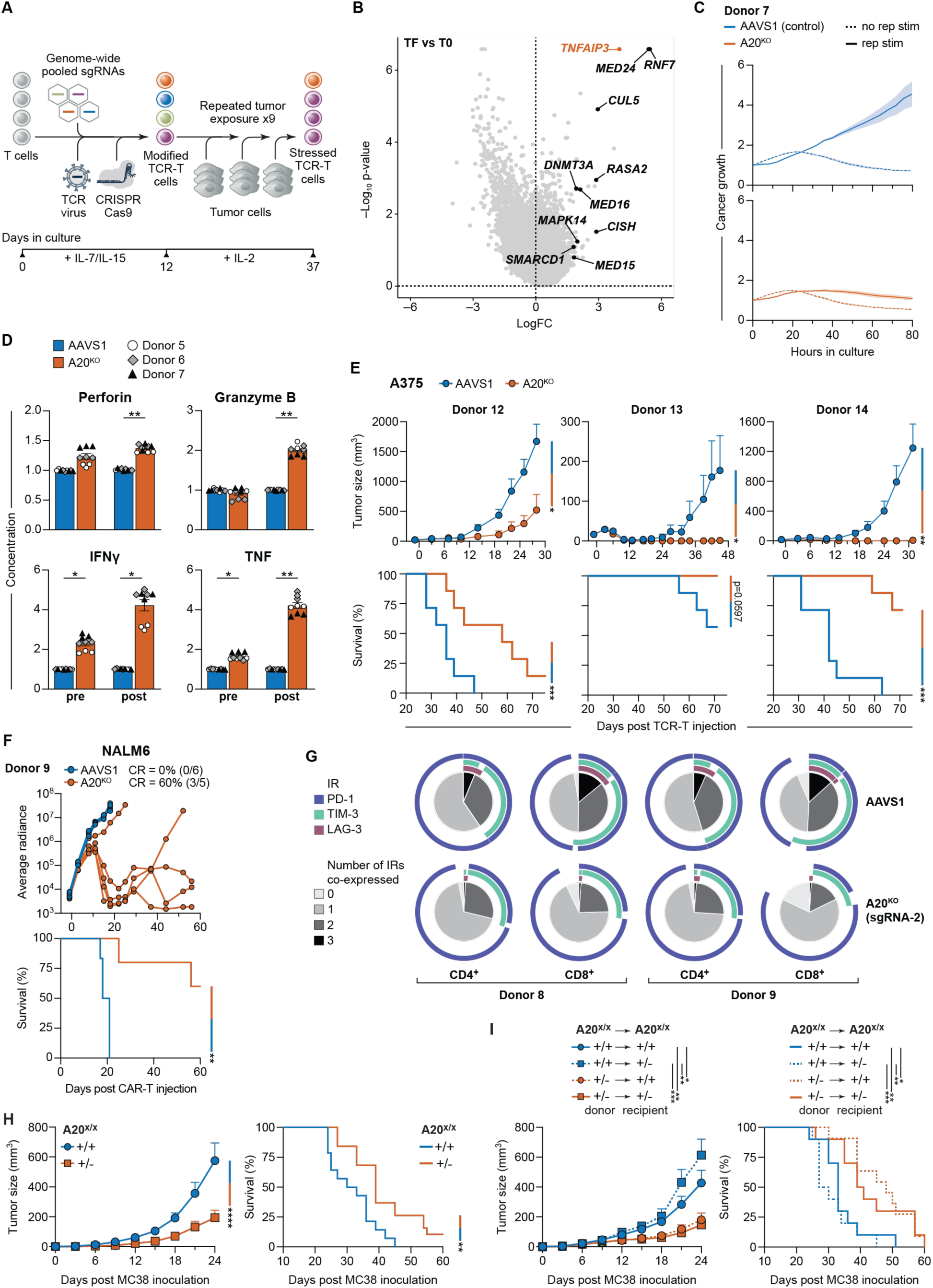
A genome-wide CRISPR perturbation screen identifies A20/*TNFAIP3* as a key regulator of T cell dysfunction upon chronic antigen stimulation. **(A)** Graphic depiction of genome-wide CRISPR screen. **(B)** Volcano plot portraying gene level scores for log fold change (FC; x axis) and p-value (y axis) generated by MAgECK analysis for TF versus T0. Screen includes two human T cell donors. Highlighted genes include A20/*TNFAIP3* (red) and genes with log FC ≥ 1.8 and known roles in T cell fitness (black). **(C)** Normalized growth curves of NY-ESO-1^+^ A375 melanoma target cells. 1G4 TCR-T cells without prior repetitive stimulation (no rep stim) or after 5 prior rounds of repetitive stimulation (rep stim) were co-cultured with A375 cells at an effector:target (E:T) ratio of 1:2. Curves depict mean ± SEM of n = 3 technical replicates from one of two T cell donors. **(D)** Normalized supernatant concentrations (pg/ml) of secreted effector granule proteins (top row) or cytokines (bottom row), as determined by LEGENDplex. TCR-T cells were analyzed before (pre) or after (post) 3 rounds of repetitive stimulation. Graphs depict mean ± SEM of n = 3 donors with n = 3 technical replicates per donor. Each technical replicate was normalized to mean of AAVS1 group for corresponding donor and condition. Significance was assessed using paired t test. **(E)** (Above) Averaged A375 growth curves in mice treated with 1G4 TCR-T cells. n = 7 mice per group using one T cell donor. (Below) Corresponding survival analysis. **(F)** (Above) Individual NALM6 growth curves in mice treated with CD19 CAR-T cells. n = 5 mice per group using one T cell donor. Radiance units = photons/s/cm^2^/steradian. (Below) Corresponding survival analysis. CR = complete response. **(G)** SPICE analyses of IR co-expression on CD4^+^ and CD8^+^ CAR-T cells isolated from bone marrow of NALM6-bearing mice. n = 3 mice per group per donor of n = 2 T cell donors. **(H)** Averaged MC38 growth (top) and survival (bottom) curves in A20 haploinsufficient (A20^+/−^) versus WT littermate (A20^+/+^) mice. n = 14-19 mice per group combined from three separate experiments. **(I)** Averaged MC38 growth (top) and survival (curves) in bone marrow chimeric mice of the indicated donor and recipient genotypes. n = 10-11 mice per group combined from two separate experiments. **(E,F,H,I)** Growth curves depict mean + SEM; significance was assessed using two-way ANOVA. Survival curve significance was assessed using a log-rank test. *p<0.05; **p<0.01; ***p<0.001; ****p<0.0001.

We further investigated whether A20 ablation could enhance human T cell function in the setting of chronic antigen exposure. We edited 1G4 TCR-T cells either with an A20-targeted sgRNA not included in the initial screen (sgRNA-1) or with a control sgRNA targeting the AAVS1 safe harbor locus (Supp. Table 6). After acute TCR stimulation, A20-targeted (A20^KO^) cells demonstrated a trend toward increased proliferative capacity (**Supp. Fig. 1F**). Furthermore, after repeated stimulation, A20^KO^ cells displayed improved tumor killing (**Fig. 1C; Supp. Fig. 1G**) and increased secretion of effector cytokines and cytotoxic granule proteins (**Fig. 1D**) when compared with AAVS1 control cells. These findings suggest that A20^KO^ TCR-T cells resist functional decline induced by chronic antigenic exposure.

### A20 ablation enhances ACT anti-tumor activity *in vivo*

To investigate whether A20 ablation could improve human ACTs *in vivo*, we challenged immunodeficient NOD-*scid* IL2Rγ^null^ (NSG) mice with A375 melanoma and treated with 1G4 TCR-T cells. Remarkably, mice that received A20^KO^ TCR-T cells suppressed A375 growth significantly better than recipients of AAVS1 TCR-T cells (**Fig. 1E**). We then generated A20^KO^ or AAVS1 CD19 *TRAC*-CAR-T cells, where a CD19 CAR with a CD28 costimulatory endodomain is integrated into the *TRAC* locus, previously shown to outperform conventionally generated CAR-T cells^40^. We tested their ability to control CD19^+^ luciferase-expressing NALM6 B cell leukemia growth in NSG mice. Serial monitoring via bioluminescence imaging (BLI) revealed that mice treated with A20^KO^ CAR-T cells controlled NALM6 growth significantly better than recipients of AAVS1 CAR-T cells, and achieved a higher complete response (CR) rate (**Fig. 1F**). We validated these findings using a different A20-targeted sgRNA (sgRNA-2; Supp. Table 6) and using two T cell donors (**Supp. Fig. 1H-1I**). Upon flow cytometric profiling of bone marrow in treated mice, we discovered that only 18-29% of A20^KO^ CAR-T cells co-expressed two or more IRs, compared with 40-51% of control AAVS1 CAR-T cells (**Fig. 1G**). These findings suggest that A20^KO^ CAR-T cells persist in more functional states than AAVS1 CAR-T cells *in vivo*. We further observed a trend toward increased numbers of A20^KO^ CAR-T cells within the bone marrow TME (**Supp. Fig. 1J**), but no clear alteration in memory phenotypes (**Supp. Fig. 1K**). Overall, A20 ablation in human TCR-T or CAR-T cells potently enhances their ability to suppress tumor growth *in vivo*.

To test whether modulating A20 expression levels can limit anti-tumor immunity in immunocompetent settings, we compared the growth of MC38 – a widely used colon adenocarcinoma derived from C57BL/6 inbred mice and considered to be a “hot” tumor that responds well to immunotherapy – in heterozygous A20^+/−^ mice and littermate control wild-type (WT; A20^+/+^) mice. Of note, A20^+/−^ mice appear grossly normal without spontaneous inflammation, but have reduced A20 expression and increased susceptibility to experimentally-induced inflammation^41,42^; homozygous A20^−/−^ mice die perinatally and are hence not suitable for testing anti-tumor responses^10^. A20^+/−^ mice consistently suppressed MC38 growth better than A20^+/+^ littermates, with improved overall survival (**Fig. 1H**). We next tested the anti-tumor efficacy of A20^+/−^ haploinsufficiency in immune cells by generating bone marrow chimeric mice reconstituted with A20^+/−^ versus A20^+/+^ hematopoietic cells, and challenging these mice with MC38. Chimeras bearing A20^+/−^ hematopoietic cells suppressed MC38 tumor growth better than chimeras containing A20^+/+^ hematopoietic cells (**Fig. 1I**). Together, these findings show that reduction of A20 expression in hematopoietic cells enhances anti-tumor immunity. When combined with our human ACT results, they suggest that modulating A20 in T cells drives this superior anti-tumor immunity.

**Supp. Fig. 1.**
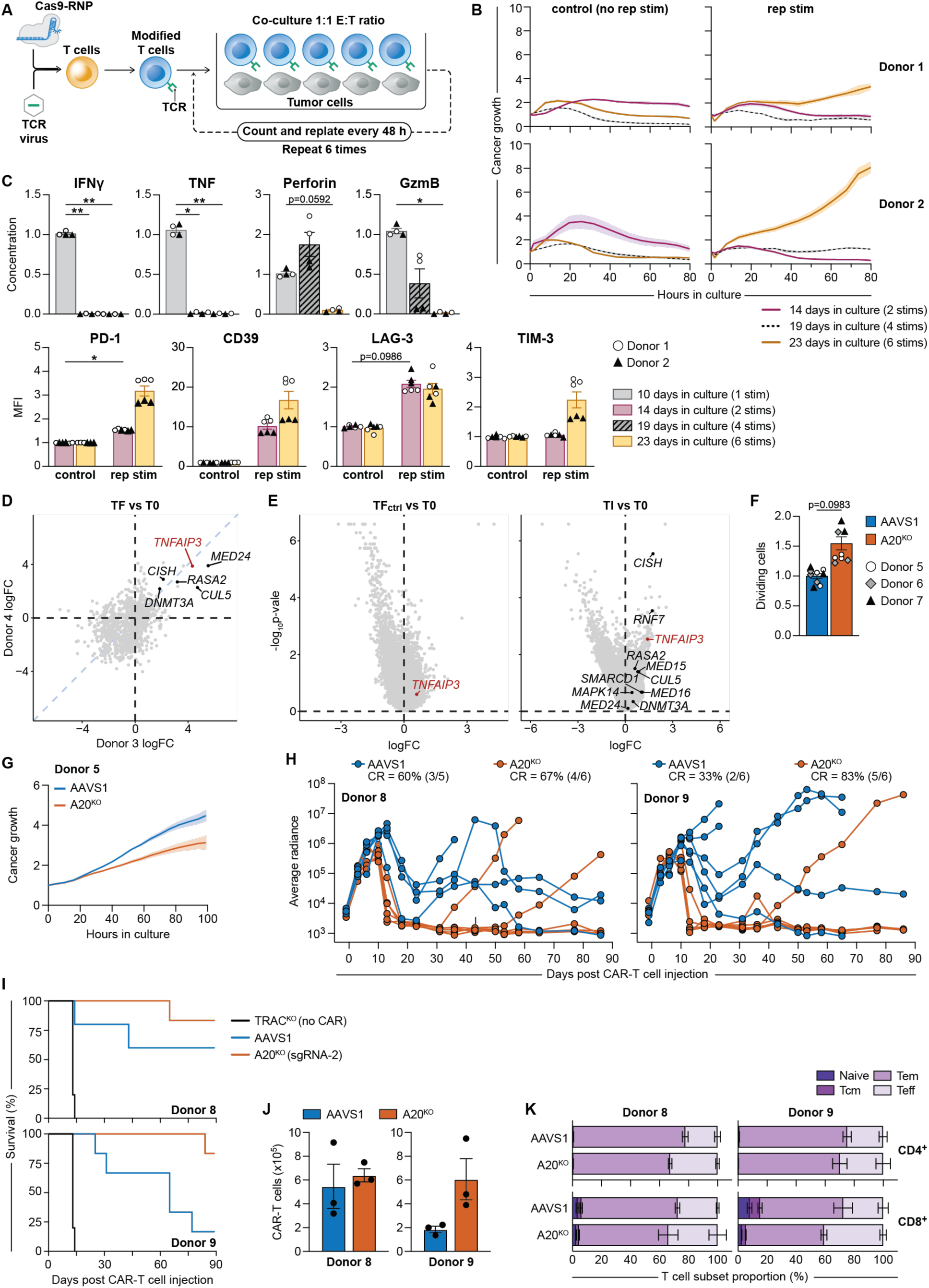
A20 is a key regulator of human T cell dysfunction upon chronic antigen stimulation. **(A)** Schematic illustration of the human TCR-T cell repetitive stimulation assay. **(B)** Normalized cancer growth curves of NY-ESO-1^+^ A375 melanoma target cells co-cultured with 1G4 TCR-T cells. Right panels depict co-cultures with TCR-T cells that had already undergone either 2, 4 or 6 rounds of repetitive tumor cell stimulation (rep stim; 14, 19 and 23 days in culture, respectively). Left panels depict co-cultures with control TCR-T cells that were matched for time in culture but had not undergone repetitive tumor cell stimulation (no rep stim). Effector:target (E:T) ratio = 1:1. Growth curves depict mean ± SEM of n = 3 technical replicates from n = 2 T cell donors. **(C)** (Top row) Normalized supernatant concentrations (pg/ml) of secreted effector molecules, as determined by LEGENDplex. Analysis shows TCR-T cells that had undergone either 1, 4, or 6 rounds of repetitive stimulation (10, 19 or 23 days in culture, respectively). (Bottom row) Normalized IR expression on TCR-T cells. Analysis shows TCR-T cells after 2 or 6 rounds of repetitive tumor antigenic stimulation (rep stim) compared with TCR-T cells matched for time in culture but not subjected to repetitive stimulation (control). MFI = median fluorescence intensity. Each technical replicate was normalized to the mean value of the control (no rep stim; bottom row) or 10 days in culture group (top row) for the corresponding donor. Bar graphs depict mean ± SEM of n = 2 donors, with n = 2-3 technical replicates per group per donor. Significance was assessed using paired t test. *p<0.05; **p<0.01. **(D)** Donor to donor correlation of TF versus T0. Highlighted genes include A20/*TNFAIP3* (red) and genes with known roles in T cell fitness (black). **(E)** Volcano plot comparing gene level scores from TFctrl versus T0 groups (left; n = 2 T cell donors) or from TI versus T0 groups (right; n = 1 T cell donor). Highlighted genes include A20/*TNFAIP3* (red) and genes with known roles in T cell fitness (black). **(F)** Normalized dividing TCR-T cells (CellTrace Violet^lo^), as determined by proliferation assay after 72 h of stimulation. Each technical replicate was normalized to mean value of AAVS1 for corresponding donor. Bar graphs depict mean ± SEM of n = 3 donors with n = 3 technical replicates per group per donor. Significance was assessed by paired t test. **(G)** Normalized cancer growth curves of NY-ESO-1^+^ A375 melanoma target cells. TCR-T cells after 5 rounds of repetitive stimulation were co-cultured with A375 cells at E:T ratio of 1:4. Growth curves depict mean ± SEM of n = 3 technical replicates from n = 1 T cell donor. **(H)** Individual NALM6 growth curves of mice treated with CAR-T cells. Left panel depicts different donor than that used in Figure 1F. Right panel depicts same donor as used in Figure 1F, but different sgRNA used to target A20^KO^. n = 6 mice per group. CR = complete response. **(I)** Survival analysis for (H). Significance was assessed using a log-rank test. **(J)** Absolute numbers of total (CD4^+^ and CD8^+^ combined) CAR-T cells isolated from bone marrow of NALM6-bearing mice 14 days after CAR-T cell injection (mean ± SEM, n = 2 T cell donors, n = 3 mice per group per donor). Significance was assessed using unpaired t test with Welch’s correction. **(K)** Differentiation status of CD4^+^ or CD8^+^ CAR-T cells from (e). Naive = CD45RA^+^CD62L^+^; central memory (Tcm) = CD45RA^−^CD62L^+^; effector memory (Tem) = CD45RA^−^ CD62L^−^; effector (Teff) = CD45RA^+^CD62L^−^ (mean ± SEM, n = 2 T cell donors, n = 3 mice per group per donor). **(D,E)** Statistics were determined by MAgECK analysis.

### Inactivation of the zinc finger 7 domain of A20 invigorates anti-tumor immunity

Complete loss of A20 from cells can perturb both cellular activation and cell death responses in distinct cell types and differentiation states^10,16,41,43,44^. These disparate outcomes reflect the ability of A20 to regulate various ubiquitinated signaling complexes^11,16,45^. The A20 protein contains several structurally defined biochemical motifs that perform distinct biochemical functions (**Fig. 2A**).^46–52^ Selective mutation of individual A20 motifs may thus differentially impact T cell functions such as activation, differentiation, cytokine secretion, and/or survival. Accordingly, we sought to investigate the domains and associated biochemical functions of A20 that regulate acute T cell responses. We started with ESM1b, a protein large language model^53,54^, and utilized a modified workflow capable of predicting the phenotypic consequences of missense variants throughout the genome^53,54^. Specific interrogation of the A20 open reading frame highlighted several hotspot regions within A20 where missense variants were predicted to have a high likelihood of functional impact (**Fig. 2A**). To complement this approach with a defined functional outcome, we reanalyzed data from our large-scale BE screen in human T cells^7^. This screen utilized a massive sgRNA library tiled across the coding sequences of known regulators of T cell activity, including A20, to probe for missense mutations capable of enhancing T cell effector functions. Briefly, T cells were activated, transduced with BEs and sgRNAs, expanded in cytokines, and then restimulated. T cells with the highest level of activation (e.g., TNF production) upon restimulation were then flow-sorted and sequenced to uncover enriched sgRNAs and their associated missense mutations. Analysis of this BE mutagenesis screen revealed sgRNAs that linked A20 missense mutations with enhanced TNF production by activated T cells (**Fig. 2A**). Alignment of our ESM1b and BE screen analyses nominated three A20 domains as potentially critical for T cell functionality: the ovarian tumor (A20^OTU^) domain, the seventh zinc finger motif (A20^ZF7^), and to a lesser extent the fourth zinc finger motif (A20^ZF4^) (**Fig. 2A**). The A20^OTU^ and A20^ZF4^ domains harbor a deubiquitinase (DUB) and an E3 ubiquitin (Ub) ligase, respectively, and A20^ZF4^ can also bind to K63 Ub chains^9^. The A20^ZF7^ motif binds to M1 linked Ub chains, but has no known enzymatic activity^51,52,55,56^.

**Fig. 2.**
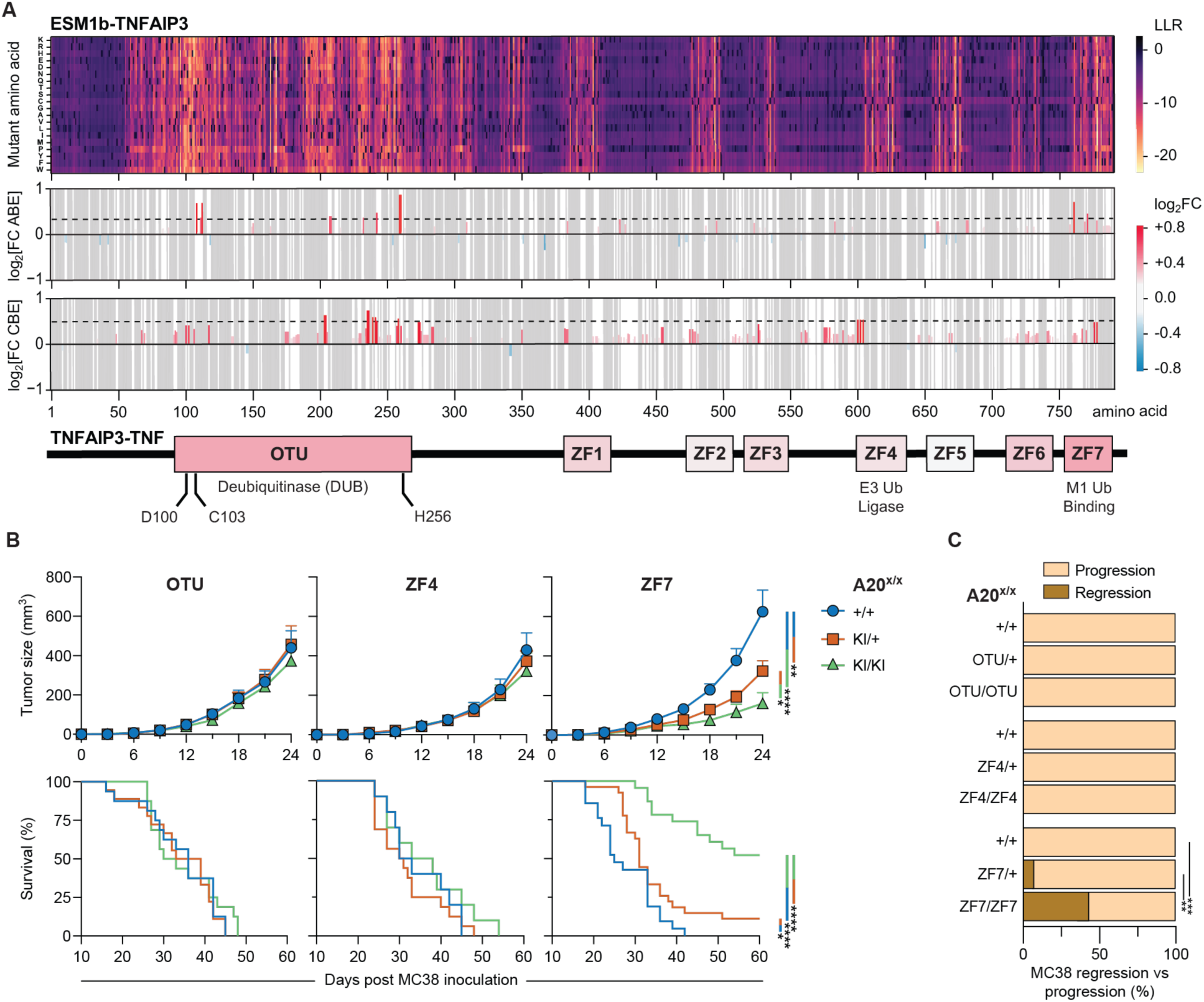
Inactivation of A20’s zinc finger 7 domain invigorates anti-tumor immunity. **(A)** (Top) Heatmap depicting ESM1b log likelihood ratio (LLR) score for all possible amino acid missense mutations (left) along the length of the A20 protein, with more negative score indicating greater likelihood of pathogenicity. (Middle) Average effects log2FC on human T cell TNF production by missense mutation base-edits across the open reading frame of A20/*TNFAIP3*, using either adenine (ABE; above) or cytosine (CBE; below) base-editors. Red and blue indicate increased versus decreased TNF production over unedited cells, respectively. Dashed line indicates average log2FC of all knockout guides within the ABE or CBE screen targeting the first 250 amino acids. Gray indicates amino acid residues not targeted by screen. (Bottom) Annotated domains of A20 protein (UniProt), with box colors indicating average effect of base-edit screen guides targeting those domains. Annotation includes DUB catalytic triad of OTU domain (D100, C103, H256). Ub = ubiquitin. **(B)** (Above) Averaged MC38 growth curves in A20^OTU^ (left), A20^ZF4^ (middle), and A20^ZF7^ (right) KI mice. Growth curves depict mean + SEM of n = 16-18 (A20^OTU^), 10-16 (A20^ZF4^) or 21-27 (A20^ZF7^) mice per genotype combined from two to four independent experiments. Significance was assessed using two-way ANOVA or mixed-effects analysis. (Below) Corresponding survival analyses. Significance was assessed using a log-rank test. **(C)** Proportion of mice from (B) that exhibited MC38 clearance versus those that did not (progression). Significance was assessed using Fisher’s exact test. *p<0.05; **p<0.01; ***p<0.001; ****p<0.0001.

To understand how these A20 motifs regulate anti-tumor immunity *in vivo*, we leveraged our unique series of A20 knock-in (KI) mice carrying germline hypomorphic missense mutations that precisely abrogate the functions of either the A20^OTU^, A20^ZF4^, or A20^ZF7^ motifs^11,12^, leaving the remainder of the A20 protein intact. Of note, while A20^−/−^ mice die shortly after birth, A20^OTU/OTU^, A20^ZF4/ZF4^, and A20^ZF7/ZF7^ mice live over 8 months, thereby allowing us to analyze their responses to implanted tumors. Using the MC38 model, we found that tumor growth in A20^OTU^ or A20^ZF4^ KI mice matched that of wild-type (WT; A20^+/+^) littermates (**Fig. 2B**), arguing against a dominant role for these domains in regulating anti-tumor immunity. In striking contrast, both heterozygous A20^ZF7/+^ and homozygous A20^ZF7/ZF7^ mice exhibited significantly slower tumor growth and prolonged survival when compared with A20^+/+^ littermates (**Fig. 2B**). Notably, tumor growth did not diverge between A20^+/+^ and A20^ZF7^ KI mice until days 9-12 following tumor inoculation (**Fig. 2B**). This timing hinted at an important role for the A20^ZF7^ domain in regulating tumor-specific T cell responses, which typically require 7-10 days for naïve antigen-specific lymphocytes to become activated, proliferate, and differentiate into effector cells. Moreover, after initial growth, MC38 tumor clearance was observed in 7% and 43% of A20^ZF7/+^ and A20^ZF7/ZF7^ mice, respectively. By contrast, no tumor clearance was observed in A20^+/+^, A20^OTU^ KI, or A20^ZF4^ KI mice (**Fig. 2C**). Thus, A20^ZF7^ inactivation produces a robust anti-tumor immune response capable of sustained tumor rejection.

The vigor of anti-tumor immune responses varies between tumor types and between host immune backgrounds. We therefore tested responses of A20^+/+^, A20^ZF7/+^ and A20^ZF7/ZF7^ littermates to challenges with two additional heterotopic tumor models. We first used B16F10 melanoma, a C57BL/6 “cold” tumor that responds poorly to immunotherapy. As with MC38, both A20^ZF7/+^ and A20^ZF7/ZF7^ mice suppressed B16F10 tumor growth better than A20^+/+^ littermates, accompanied by prolonged survival (**Supp. Fig. 2A-2B**). Next, we used CT26 colon carcinoma, a “hot” tumor derived from BALB/c mice. BALB/c mice exhibit more robust type 2 immune responses characterized by IL-4, IL-5, and IL-13 expression, and less robust type 1 responses typified by IFNγ and TNF expression^57^. We backcrossed A20^ZF7^ mice to BALB/c mice for seven generations and, upon challenge with CT26, again observed that both A20^ZF7/+^ and A20^ZF7/ZF7^ mice suppressed tumor growth better than A20^+/+^ littermates (**Supp. Fig. 2A-2B**). Hence, the A20^ZF7^ domain restrains anti-tumor immunity against multiple tumor types and across divergent immune backgrounds.

Augmented primary anti-tumor immune responses can lead to enhanced memory responses. On the other hand, some genetic enhancements of acute CAR-T cell responses can compromise the durability of these responses, resulting in tumor relapse^58^. Also of note, A20 deficiency reduces memory T cell responses to bacterial infection^18^. We therefore investigated whether A20^ZF7/ZF7^ mice that cleared their primary tumor challenge also developed a robust anti-tumor memory response. We rechallenged these mice with four times the number of MC38 cells used in the primary challenge, and compared tumor growth with age-matched A20^ZF7/ZF7^ mice receiving their primary tumor challenge (i.e. naive to tumor; **Supp. Fig. 2C**). As no A20^+/+^ mice cleared their primary challenge, this genotype was not available for comparison in these experiments. Strikingly, A20^ZF7/ZF7^ mice that cleared their primary tumor challenge nearly completely resisted their secondary challenge, with significantly less tumor growth and prolonged survival when compared to age-matched A20^ZF7/ZF7^ mice receiving their primary tumor challenge (**Supp. Fig. 2D**). Therefore, abrogation of the A20^ZF7^ motif also confers a strong memory anti-tumor immune response.

**Supp. Fig. 2.**
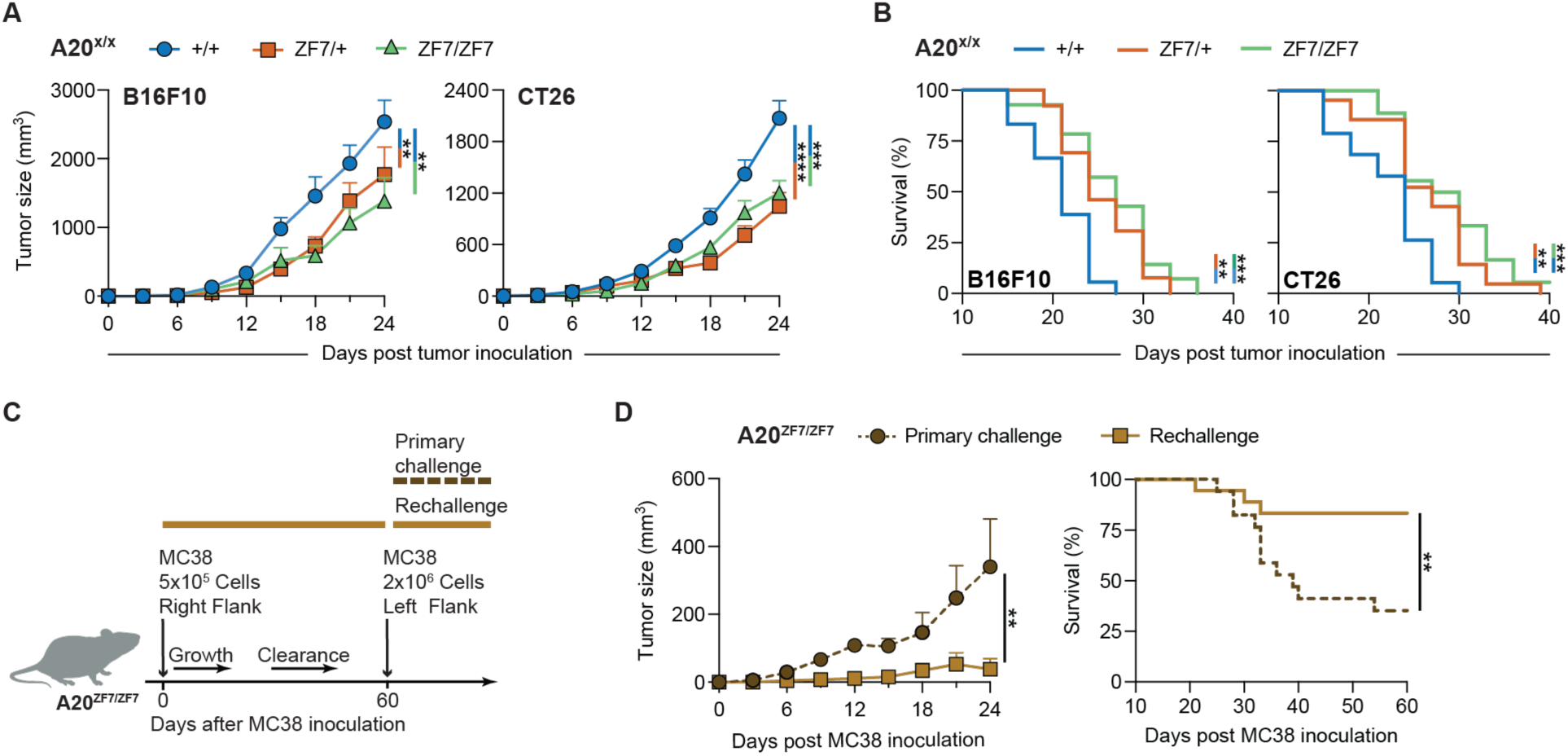
A20^ZF7^ inactivation augments anti-tumor immunity across tumor models and promotes anti-tumor memory. **(A)** Averaged B16F10 (left) and CT26 (right) growth curves in A20^ZF7^ KI mice. n = 13-18 (B16F10) or 18-21 (CT26) mice per genotype combined from three independent experiments. **(B)** Survival analyses corresponding to (A). **(C)** Overview of workflow for MC38 rechallenge experiments of A20^ZF7/ZF7^ mice that cleared their first challenge. **(D)** (Left) Averaged MC38 growth curves from A20^ZF7/ZF7^ mice that cleared their initial challenge and were rechallenged with MC38, compared with age-matched A20^ZF7/ZF7^ mice that were naive to tumor (primary challenge). n = 9-17 mice per group combined from three independent experiments. (Right) Corresponding survival analysis. **(A,B,D)** Growth curves depict mean + SEM; significance was assessed using two-way ANOVA. Survival significance was assessed using a log-rank test. **p<0.01; ***p<0.001; ****p<0.0001.

### CD8^+^ T cell-intrinsic A20^ZF7^ inactivation invigorates anti-tumor immunity

Mutation of A20^ZF7^ perturbs homeostasis of both innate and adaptive immune cells^12^. We thus investigated which immune cells are critical for the superior anti-tumor immunity of A20^ZF7/+^ and A20^ZF7/ZF7^ mice. We began by interbreeding A20^ZF7^ mice with *Rag1*^−/−^ mice to remove B and T lymphocytes. RAG1 deficiency abolished the advantages in MC38 tumor suppression and survival of both A20^ZF7/+^ and A20^ZF7/ZF7^ mice (**Supp. Fig. 3A**), while also eliminating tumor clearance (**Supp. Fig. 3B**). The critical RAG1-dependent lymphocytes did not appear to be B cells or γδ T cells, as A20^ZF7/+^ and A20^ZF7/ZF7^ mice with genetic ablation of either B cells (*Ighm*^−/−^ a.k.a. μMT) or γδ T cells (*Tcrd*^−/−^) retained tumor suppression advantages over A20^+/+^μMT and A20^+/+^*Tcrd*^−/−^ controls, respectively (**Supp. Fig. 3A-3B**). We therefore turned our attention to CD4^+^ and CD8^+^ T cells. When compared to isotype control-treated mice, antibody-mediated depletion of CD4^+^ T cells greatly enhanced MC38 suppression and clearance in A20^+/+^ and A20^ZF7/+^ mice (**Fig. 3A**, **3C**), consistent with prior reports^59,60^ and likely attributable to depletion of regulatory CD4^+^ T cells (Tregs). Meanwhile, CD4^+^ T cell depletion from A20^ZF7/ZF7^ mice had minimal impact on MC38 suppression or clearance (**Fig. 3A**, **3C**), arguing against a major role for CD4^+^ T cells in orchestrating superior anti-tumor immunity in these mice. Conversely, antibody-mediated depletion of CD8^+^ T cells from A20^ZF7/+^ and A20^ZF7/ZF7^ mice completely abolished their advantages in tumor suppression and clearance (**Fig. 3A**, **3C**), as did genetic ablation of CD8^+^ T cells (*Cd8a*^−/−^; **Fig. 3B-3C**). Of note, antibody-mediated CD8^+^ T cell depletion had minimal impact on MC38 growth in A20^+/+^ mice (**Fig. 3A**), suggesting their CD8^+^ TILs are not significantly contributing to tumor suppression, and are thus likely highly dysfunctional and potentially immunosuppressive^61^. Together these results highlight the centrality of CD8^+^ T cells to superior anti-tumor immunity in A20^ZF7^ KI mice.

**Fig. 3.**
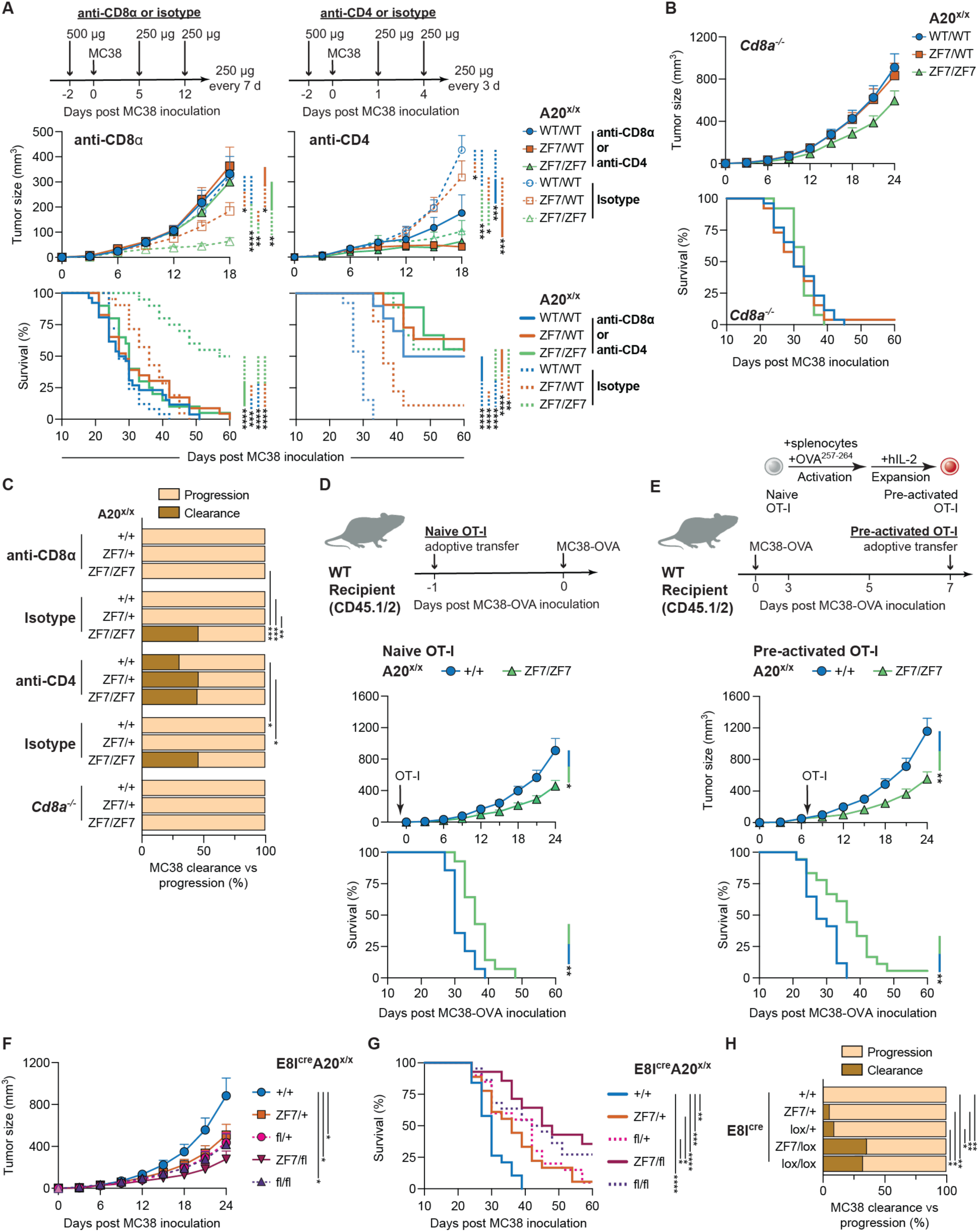
CD8^+^ T cell-intrinsic A20^ZF7^ inactivation invigorates anti-tumor immunity. **(A)** Averaged MC38 growth (top) and survival (bottom) curves from A20^ZF7^ KI mice with indicated antibody-mediated CD4^+^ T cell (anti-CD4) or CD8^+^ T cell (anti-CD8α) depletion. Isotype = isotype control antibody for anti-CD4 or anti-CD8α. n = 9-13 (anti-CD4) or 20-26 (anti-CD8α) mice per group combined from two (anti-CD4) or four (anti-CD8α) independent experiments. **(B)** Averaged MC38 growth (top) and survival (bottom) curves from A20^ZF7^ KI mice with genetic CD8^+^ T cell ablation (*Cd8a^−/−^*). n = 13-26 mice per genotype combined from three separate experiments. **(C)** Proportion of mice from (A) and (B) that exhibited MC38 clearance versus those that did not (progression). **(D)** Averaged MC38-OVA growth (above) and survival (below) curves from mice that received adoptive transfer of naive OT-I cells. n = 14 mice per group combined from three separate experiments. **(E)** Averaged MC38-OVA growth (above) and survival (below) curves from mice that received adoptive transfer of pre-activated OT-I cells. n = 17-18 mice per group combined from three separate experiments. **(F)** Averaged MC38 growth curves from E8I^cre^ mice interbred with mice carrying floxed A20 (A20^fl^) and/or A20^ZF7^ KI alleles. n = 19-25 mice per genotype combined from four separate experiments. **(G)** Survival curves from (F). **(H)** Proportion of mice from (F) that exhibited MC38 clearance versus progression. **(A,B,D,E,F,G)** Growth curves depict mean + SEM; significance was assessed using two-way ANOVA. Survival curve significance was assessed using log-rank test. **(C,H)** Significance was assessed using Fisher’s exact test. *p<0.05; **p<0.01; ***p<0.001; ****p<0.0001.

We next investigated whether A20^ZF7^ inactivation specifically within CD8^+^ TILs could improve anti-tumor immunity. We interbred A20^ZF7^ KI mice with OT-I transgenic mice expressing a TCR specific for the ovalbumin peptide OVA^257–264^. We then adoptively transferred either naive A20^ZF7/ZF7^ OT-I or control A20^+/+^ OT-I cells (CD45.2) into congenic (CD45.1) WT hosts (**Fig. 3D**).

After challenging these host mice with OVA-expressing MC38 cells (MC38-OVA), we observed that recipients of naïve A20^ZF7/ZF7^ OT-I cells suppressed tumor growth and prolonged survival better than recipients of A20^+/+^ OT-I cells (**Fig. 3D**). Similarly enhanced anti-tumor immunity was observed when naïve A20^ZF7/ZF7^ OT-I cells were tested against B16-OVA melanoma (**Supp. Fig. 3C**). We next employed an adoptive transfer model using pre-activated OT-I cells (**Fig. 3E**). This model focuses specifically on OT-I effector phase functions, bypassing any potential differences that arise between A20^+/+^ and A20^ZF7/ZF7^ OT-I cells at the lymph node priming stage. Furthermore, this model more closely reflects ACT production. As with naive cells, adoptive transfer of pre-activated A20^ZF7/ZF7^ OT-I cells outperformed A20^+/+^ OT-I cells (**Fig. 3E**).

**Supp. Fig. 3.**
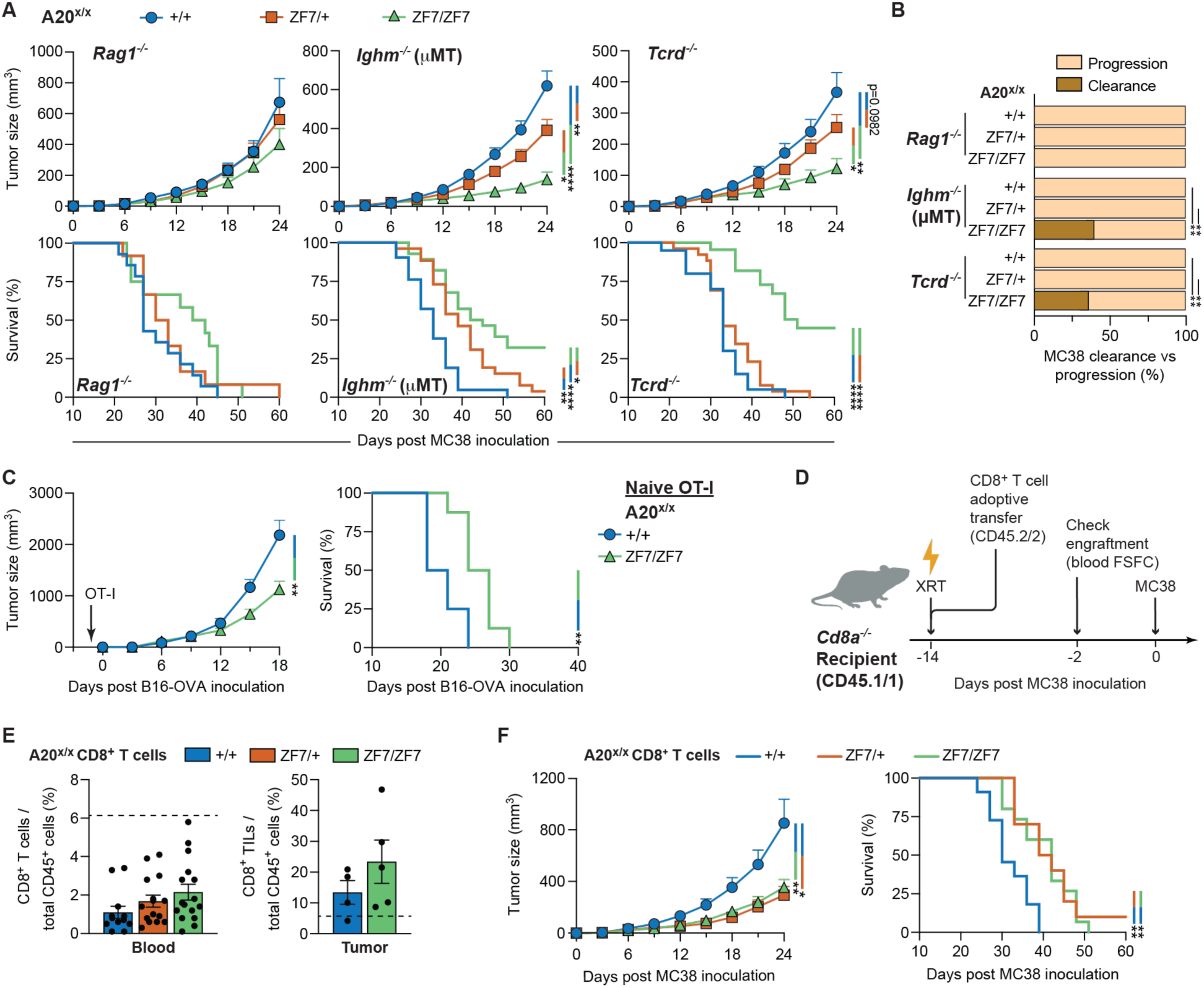
A20^ZF7^ inactivation specifically within CD8^+^ T cells augments anti-tumor immunity. **(A)** (Above) Averaged MC38 growth curves from mice with indicated genetic ablation of lymphocyte subsets. (Below) Corresponding survival analyses. n = 12-14 (*Rag1^−/−^*), 20-34 (μMT) or 20-26 (*Tcrd^−/−^*) mice per genotype combined from three to five separate experiments. **(B)** Proportion of mice from (A) that exhibited MC38 clearance versus those that did not (progression). Significance was assessed using Fisher’s exact test. **(C)** Averaged B16-OVA growth (left) and survival (right) curves from mice that received adoptive transfer of naïve OT-I cells. n = 8 mice per group combined from two separate experiments. **(D)** Overview of workflow for reconstitution of *Cd8a^−/−^* mice with CD8^+^ T cells. XRT = sublethal irradiation. FSFC = full spectrum flow cytometry. **(E)** Bar graphs depicting CD8^+^ T cell reconstitution of blood (left) and MC38 tumors (right) in *Cd8a^−/−^* mice. Blood dashed line represents mean blood CD8^+^ T cell proportion from 5 age-matched non-irradiated non-tumor-bearing WT (*Cd8a^+/+^*) mice. Tumor dashed line represents mean CD8^+^ TIL density from day 12-15 tumors in WT (A20^+/+^) mice (see Figure 4A). Bar graphs depict mean ± SEM of 4-16 mice per group from 1-2 independent experiments. Significance was assessed using one-way ANOVA (blood) or unpaired t test with Welch’s correction (tumor). **(F)** (Left) Averaged MC38 growth curves of *Cd8a^−/−^* mice reconstituted with CD8^+^ T cells. n = 10-15 mice per group combined from two separate experiments. (Right) Corresponding survival analysis. **(A,C,D,G)** Growth curves depict mean + SEM; significance assessed using two-way ANOVA. Survival significance was assessed using log-rank test. *p<0.05; **p<0.01; ****p<0.0001.

To test the ability of A20^ZF7^ KI CD8^+^ T cells to kill MC38 tumors via the recognition of endogenous tumor antigens, we took two separate approaches. First, we utilized *Cd8a*^cre^ (a.k.a. E8I^cre^) mice crossed with A20^ZF7^ KI mice and/or floxed A20 mice (A20^fl^) to generate mice with either CD8^+^ T cell-restricted A20 deficiency (E8I^cre^A20^fl/fl^ mice) or A20^ZF7^ hemizygosity (E8I^cre^A20^ZF7/fl^ mice). E8I^cre^A20^fl/fl^ mice outperformed E8I^cre^A20^+/+^ littermates in tumor growth suppression (**Fig. 3F-3G**) and achieved a clearance rate comparable to A20^ZF7/ZF7^ mice (27%; **Fig. 3H**), confirming that A20^KO^ CD8^+^ T cells boost anti-tumor immunity^20–22^. Remarkably, E8I^cre^A20^ZF7/fl^ mice also demonstrated a clearance rate equivalent to E8I^cre^A20^fl/fl^ mice (36%; **Fig. 3H**), and prolonged mouse survival to a greater extent than E8I^cre^A20^ZF7/+^ or E8I^cre^A20^fl/+^ mice (**Fig. 3G**), indicating that A20^ZF7^ hemizygosity within CD8^+^ T cells promotes anti-tumor immunity. Second, we reconstituted *Cd8a*^−/−^ mice (CD45.1) with congenic (CD45.2) polyclonal CD8^+^ T cells (**Supp. Fig. 3D**). Using sublethal irradiation prior to adoptive transfer, we achieved durable engraftment of the transferred CD8^+^ T cells (**Supp. Fig. 3E**). Importantly, *Cd8a*^−/−^ mice reconstituted with either A20^ZF7/+^ or A20^ZF7/ZF7^ CD8^+^ T cells suppressed MC38 growth significantly better than *Cd8a*^−/−^ mice reconstituted with control A20^+/+^ CD8^+^ T cells (**Supp. Fig. 3F**). Taken together, these data show that abrogating the A20^ZF7^ motif specifically within CD8^+^ T cells enhances anti-tumor immunity.

### A20^ZF7^ inactivation alleviates terminal exhaustion of CD8^+^ T cells

To understand how A20^ZF7^ regulates CD8^+^ TILs, we analyzed CD8^+^ TILs from tumors in A20^+/+^, A20^ZF7/+^ and A20^ZF7/ZF7^ mice by full spectrum flow cytometry (FSFC) 12-15 days after tumor inoculation. This time point was chosen to allow enough time for mice to mount an adaptive immune response, while still maintaining similar tumor sizes across genotypes. We initially observed no significant difference in total CD8^+^ TIL density within A20^+/+^ versus A20^ZF7^ KI mice, a finding that was consistent across tumor models (**Fig. 4A**). We next investigated CD8^+^ TIL subsets, as differences in CD8^+^ T cell differentiation can impact anti-tumor immunity. CD8^+^ TILs can be broadly defined as Tex (PD-1^+^) or PD-1^−^ cells, and Tex can be further divided into progenitor exhausted (Tpex) and terminally exhausted (Ttex) subsets (**Fig. 4B**)^4,62–64^. Tpex have stem-like features but possess minimal cytotoxic potential, and are defined by expression of TCF-1 (**Fig. 4B**). Tpex ultimately differentiate into Ttex^2,4^, characterized by loss of TCF-1 expression, upregulation of the IR TIM-3 (**Fig. 4B**), and increased expression of TOX. Ttex increase their expression of cytotoxic molecules but remain hypofunctional, as TOX orchestrates upregulation of various additional IRs and epigenetic silencing of T cell effector genes^65–67^. In MC38 and CT26 tumors, we did not find any significant differences in the distribution of these CD8^+^ TIL subsets across genotypes, while in B16F10 tumors we observed a greater proportion of Ttex in A20^ZF7/ZF7^ mice (**Fig. 4C**). Overall, quantitative differences in CD8^+^ TIL subsets cannot fully explain superior anti-tumor immunity in A20^ZF7^ KI mice.

**Fig. 4.**
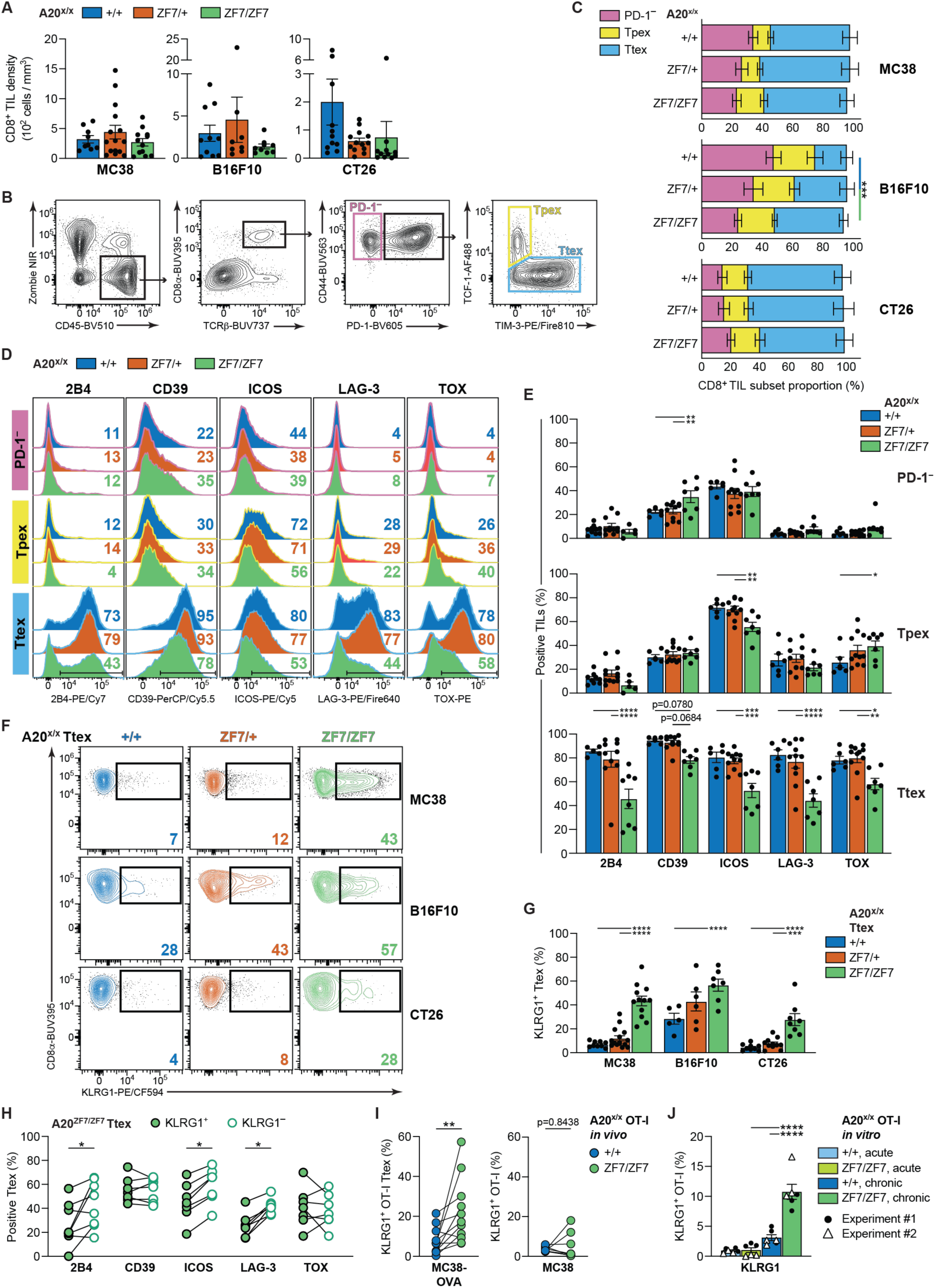
A20^ZF7^ inactivation alleviates terminal exhaustion of CD8^+^ T cells. **(A)** Total intratumoral CD8^+^ TIL density. n = 9-15 (MC38), 8-10 (B16F10), or 11-13 (CT26) samples per genotype combined from two to three separate experiments. **(B)** Gating strategy to identify CD8^+^ TIL subsets. Contour plots represent 10 concatenated samples combined from A20^+/+^, A20^ZF7/+^ and A20^ZF7/ZF7^ MC38 tumors from one experiment. **(C)** CD8^+^ TIL subset proportions from (A). **(D)** IR or TOX expression by MC38 CD8^+^ TIL subsets. Representative FSFC histograms depict 3-5 concatenated samples per genotype from one experiment. Inset = proportion (%) of cells within ranged gate. **(E)** Bar graphs showing proportion of CD8^+^ TIL subsets — PD-1^−^ (top), Tpex (middle), or Ttex (bottom) — from (C) that express IRs or TOX. **(F)** Ttex KLRG1 expression. Representative contour plots from 3-5 concatenated samples per genotype from one experiment. Inset = proportion (%) of KLRG1-expressing cells. **(G)** Proportion of Ttex from (C) that express KLRG1. **(H)** Paired analysis of IR or TOX expression by KLRG1^+^ versus KLRG1^−^ A20^ZF7/ZF7^ Ttex within MC38 tumors from (G). **(I)** Paired analysis of A20^+/+^ versus A20^ZF7/ZF7^ OT-I Ttex expression of KLRG1 following adoptive co-transfer of pre-activated OT-I cells. **(J)** KLRG1 expression by OT-I cells following *in vitro* repetitive (chronic) stimulation or acute stimulation alone, combining technical triplicates from two separate experiments. Significance was assessed using paired t test. **(A-G)** FSFC analyses of day 12-15 tumors. Significance was assessed using one-way ANOVA (A,G) or two-way ANOVA with Tukey’s multiple comparisons test (C,E). **(H,I)** Each symbol/line pair represents cells from the same tumor. n = 6-10 samples combined from two to three independent experiments. Significance was assessed using Wilcoxon matched-pairs signed rank test. *p<0.05; **p<0.01; ***p<0.001; ****p<0.0001.

We next interrogated memory phenotypes of CD8^+^ TIL subsets. Intriguingly, in later-stage (day 24) MC38 tumors, we discovered a significant increase in A20^ZF7/ZF7^ CD8^+^ TILs bearing a tissue resident memory (CD69^+^CD103; Trm) phenotype (**Supp. Fig. 4A**), which could explain in part their robust anti-tumor memory response (**Supp. Fig. 2D**). Conversely, in day 12-15 MC38 tumors, we did not observe any differences among CD8^+^ TILs bearing a Trm or central memory (CD62L^+^; Tcm) phenotype (**Supp. Fig. 4B**). Thus, at this early time point, memory CD8^+^ T cell differentiation is unlikely to account for the superior anti-tumor immunity of A20^ZF7/ZF7^ mice.

We therefore turned our attention to exhaustion by profiling the expression of IRs (2B4, CD39, ICOS and LAG-3) and TOX. With few exceptions, the fraction of PD-1^−^ TILs and Tpex that expressed these markers within MC38 tumors was similar across genotypes (**Fig. 4D-4E**). In striking contrast, a significantly lower percentage of A20^ZF7/ZF7^ Ttex expressed IRs or TOX when compared with A20^+/+^ and A20^ZF7/+^ Ttex (**Fig. 4D-4E**). Similar findings were observed in A20^ZF7/ZF7^ Ttex from B16F10 and CT26 tumors (**Supp. Fig. 4C**). Intriguingly, we also observed a marked increase in KLRG1^+^ A20^ZF7/ZF7^ Ttex across tumor models (**Fig. 4F-4G**), which notably had lower IR expression than KLRG1^−^ A20^ZF7/ZF7^ Ttex (**Fig. 4H**). KLRG1 identifies short-term effector CD8^+^ T cells, and is not commonly seen expressed on Ttex^68,69^. These phenotypic differences were not present in A20^OTU/OTU^ or A20^ZF4/ZF4^ Ttex (**Supp. Fig. 4D**). Thus, A20^ZF7^ inactivation alters CD8^+^ Ttex such that they appear less exhausted.

To test whether A20^ZF7^ inactivation impacts CD8^+^ TIL exhaustion in a truly cell autonomous manner, we adoptively co-transferred congenic pre-activated A20^+/+^ OT-I (CD45.1/1) and A20^ZF7/ZF7^ OT-I (CD45.2/2) cells into WT (CD45.1/2) recipient mice bearing MC38-OVA tumors (**Supp. Fig. 4E**), then analyzed tumors 7 days later by FSFC. Both A20^+/+^ and A20^ZF7/ZF7^ OT-I TILs were present at similar densities and differentiated into Ttex (PD-1^hi^TIM-3^+^) in similar proportions (**Supp. Fig. 4F-4G**). However, A20^ZF7/ZF7^ OT-I Ttex displayed increased levels of KLRG1 (**Fig. 4I**) and decreased levels of IRs and TOX (**Supp. Fig. 4H**) when compared with A20^+/+^ OT-I Ttex within the same TME, confirming the CD8^+^ T cell-intrinsic role of A20^ZF7^ inactivation in relieving the exhaustion phenotype.

We next wanted to gain insight into whether these phenotypic differences in A20^ZF7/ZF7^ Ttex truly required exhaustive (i.e. repetitive or chronic) TCR engagement, or were simply a byproduct of the initial (i.e. acute) TCR stimulus. We first repeated our adoptive co-transfer of pre-activated OT-I cells but replaced MC38-OVA tumor cells with MC38 tumors, thereby removing the *in vivo* antigenic stimulus and ensuring these OT-I cells were only acutely stimulated *in vitro*. Examination of OT-I TILs within MC38 tumors 7 days after adoptive co-transfer revealed densities comparable to that observed in MC38-OVA tumors (**Supp. Fig. 4G**); however, almost all A20^+/+^ and A20^ZF7/ZF7^ OT-I TILs within MC38 tumors remained PD-1^−^, with few differentiating into Ttex (**Supp. Fig. 4F-4G**). As there were not enough OT-I Ttex of either genotype to analyze within MC38 tumors, we instead profiled IRs and TOX on total A20^+/+^ and A20^ZF7/ZF7^ OT-I TILs. This analysis revealed minimal expression of KLRG1 on either A20^+/+^ or A20^ZF7/ZF7^ OT-I TILs (**Fig. 4I**). Furthermore, LAG-3 and TOX expression remained equivalently low in A20^+/+^ and A20^ZF7/ZF7^ OT-I TILs (**Supp. Fig. 4H**). Meanwhile, fewer A20^ZF7/ZF7^ OT-I TILs in MC38 tumors expressed CD39 and ICOS than A20^+/+^ OT-I TILs (**Supp. Fig. 4H**), similar to our findings in MC38-OVA OT-I Ttex. Thus, several core A20^ZF7/ZF7^ Ttex phenotypic differences (KLRG1, LAG-3, TOX) required chronic TCR engagement *in vivo*, while others were likely encoded following acute TCR stimulation (CD39, ICOS). To further support these findings, we submitted A20^+/+^ and A20^ZF7/ZF7^ OT-I cells to an *in vitro* repetitive/chronic stimulation assay benchmarked as a surrogate for exhaustion (**Supp. Fig. 4I**)^70^. For comparison, a parallel set of OT-I cells matched for time in culture received only acute stimulation (**Supp. Fig. 4I**). As with our *in vivo* analysis, this assay revealed increased KLRG1 expression (**Fig. 4J**), and decreased expression of several IRs (LAG-3, PD-1, TIM-3; **Supp. Fig. 4J**), in chronically but not acutely stimulated A20^ZF7/ZF7^ OT-I cells when compared to A20^+/+^ OT-I cells. We also again observed lower expression of CD39 and ICOS on both acutely and chronically stimulated A20^ZF7/ZF7^ OT-I cells when compared to A20^+/+^ OT-I cells (**Supp. Fig. 4J**). Together these results indicate that many of the key phenotypic differences observed in A20^ZF7/ZF7^ Ttex versus WT Ttex require repetitive TCR engagement, while others require only acute stimulation.

**Supp. Fig. 4.**
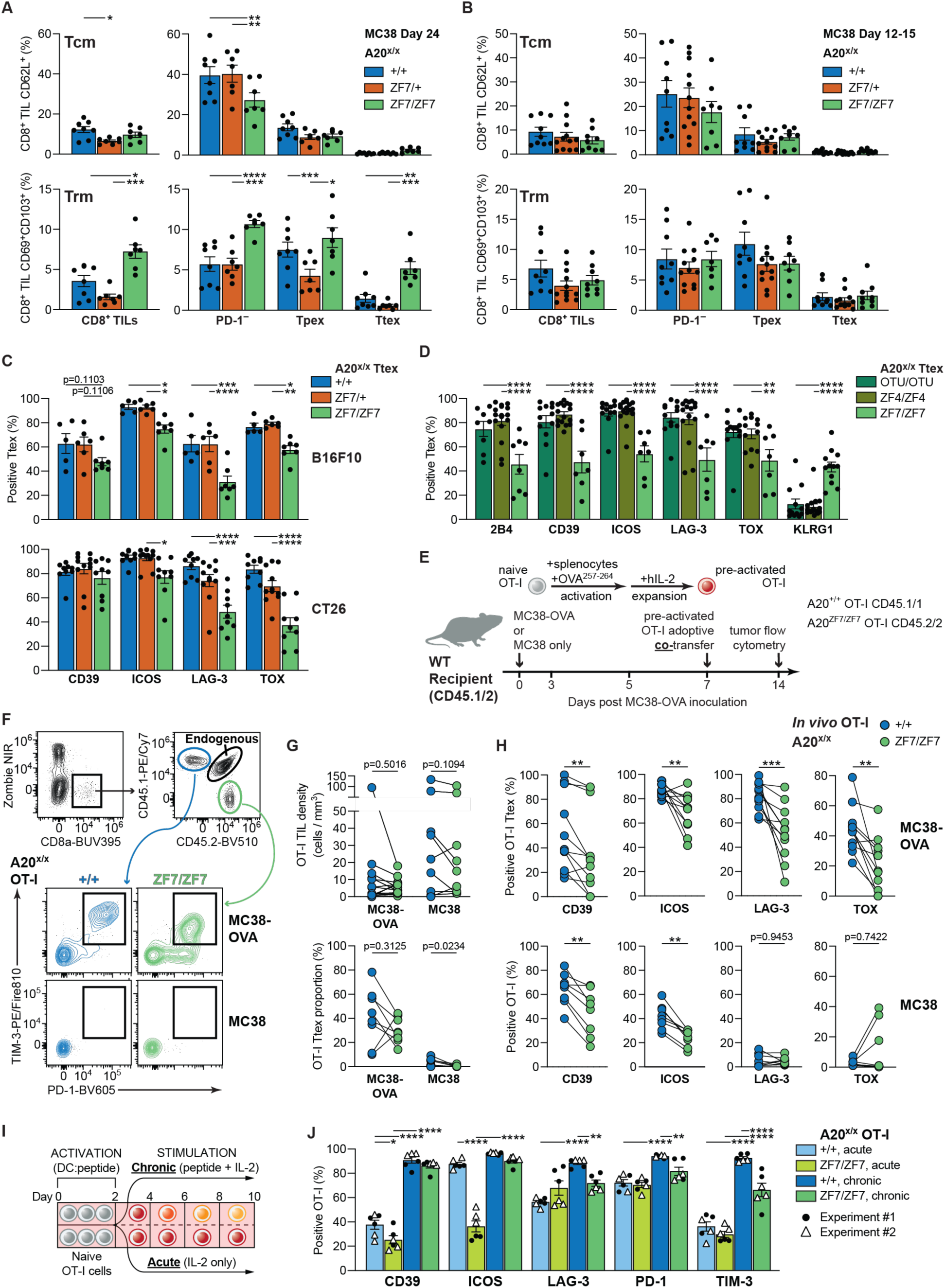
A20^ZF7^ inactivation alleviates terminal exhaustion following repetitive TCR stimulation. **(A)** Proportion of day 24 MC38 total CD8^+^ TILs (left) or TIL subsets (right) with phenotype suggestive of central memory (Tcm; CD62L^+^; top) or tissue resident memory (Trm; CD69^+^CD103^+^; bottom) differentiation. n = 7-8 mice per genotype combined from two separate experiments. **(B)** As in (A), but from day 12-15 MC38 tumors. n = 8-12 tumors per genotype combined from two separate experiments. **(C)** IR or TOX expression by Ttex from B16F10 (top) or CT26 (bottom) tumors. n = 5-7 (B16F10) or 9-11 (CT26) mice per genotype combined from two or three separate experiments. **(D)** IR, TOX or KLRG1 expression by Ttex from MC38 tumors. n = 7-15 mice per genotype combined from two or three separate experiments. A20^ZF7/ZF7^ data is repeated from Fig. 4E and 4G for comparison. **(E)** Overview of workflow for adoptive OT-I co-transfer experiments. **(F)** Gating strategy to identify A20^+/+^ (CD45.1/1) and A20^ZF7/ZF7^ OT-I (CD45.2/2) Ttex in adoptive co-transfer experiments. Contour plots represent FSFC on 9 concatenated MC38-OVA and MC38 tumors from one experiment. Endogenous = endogenous MC38-OVA or MC38 CD8^+^ TILs (CD45.1/2). **(G)** Paired analyses of OT-I TIL density (top) and proportion that differentiated into Ttex (bottom) within MC38 or MC38-OVA tumors. n = 7 samples combined from two experiments. **(H)** Paired analysis of IR and TOX expression by OT-I Ttex (MC38-OVA; top) or total OT-I TILs (MC38; bottom) following adoptive co-transfer. **(I)** Overview of workflow for *in vitro* acute versus chronic/repetitive stimulation assay. **(J)** IR expression by OT-I cells following *in vitro* acute or chronic stimulation. Bar graphs depict mean ± SEM of technical triplicates combined from two separate experiments. Significance was assessed using two-way ANOVA with Bonferroni’s multiple comparisons test. **(A-H)** FSFC analyses from day 12-15 tumors, unless otherwise stated. All bar graphs show mean ± SEM. Significance was assessed using two-way ANOVA with Tukey’s multiple comparisons test. **(G,H)** Each symbol/line pair represents cells retrieved from the same tumor. n = 9-11 samples combined from two separate experiments. Significance was assessed using Wilcoxon matched-pairs signed rank test. *p<0.05; **p<0.01; ***p<0.001; ****p<0.0001.

### Perforin, but not IFNγ or TNF, drives enhanced anti-tumor immunity in A20^ZF7/ZF7^ mice

Reduced exhaustion in A20^ZF7/ZF7^ Ttex may align with enhanced tumoricidal effector function, explaining the superior anti-tumor response of A20^ZF7/ZF7^ mice. To define the relevant anti-tumor effector mechanisms unleashed in A20^ZF7^ KI Ttex, we focused first on co-production of IFNγ and TNF, a commonly cited indicator of TIL dysfunction^4,71,72^. We observed a significant increase of IFNγ^+^ TNF^+^ Ttex in both A20^ZF7/+^ and A20^ZF7/ZF7^ mice upon *ex vivo* stimulation (**Fig. 5A-5B**), consistent with prior reports in A20^KO^ CD8^+^ TILs^20–22^. To test whether increased IFNγ or TNF production by A20^ZF7^ KI Ttex enhances anti-tumor immunity, we interbred A20^ZF7^ KI mice with *Ifng^−/−^* or *Tnf^−/−^* mice. Surprisingly, neither IFNγ or TNF deficiency alone were able to reverse the superior MC38 growth control and prolonged survival of A20^ZF7/+^ and A20^ZF7/ZF7^ mice when compared to A20^+/+^ littermates (**Fig. 5C**). In addition, A20^ZF7/ZF7^*Ifng*^−/−^ and A20^ZF7/ZF7^*Tnf*^−/−^ mice both showed tumor clearance rates similar to A20^ZF7/ZF7^ mice (**Fig. 5D**). To account for potential redundancies or compensatory actions between the anti-tumor activities of IFNγ and TNF, we interbred A20^ZF7^ mice with *Ifng*^−/−^*Tnf*^−/−^ mice that lack both cytokines. Interestingly, A20^ZF7/+^*Ifng*^−/−^ *Tnf*^−/−^ mice were no longer able to control tumor growth or prolong survival better than their A20^+/+^*Ifng*^−/−^*Tnf*^−/−^ mice counterparts (**Fig. 5C**), indicating that these cytokines together drive the superior tumor suppression observed in heterozygous A20^ZF7/+^ mice. In striking contrast, homozygous A20^ZF7/ZF7^*Ifng*^−/−^*Tnf*^−/−^ mice retained significant advantages in controlling tumor growth and prolonging survival when compared to A20^+/+^*Ifng*^−/−^*Tnf*^−/−^ mice, while maintaining a 45% clearance rate (**Fig. 5C-5D**). These findings suggested the presence of an additional IFNγ/TNF-independent tumor growth suppression mechanism unlocked only in homozygous but not heterozygous A20^ZF7^ KI mice, i.e., only below a certain threshold of A20 activity.

**Fig. 5.**
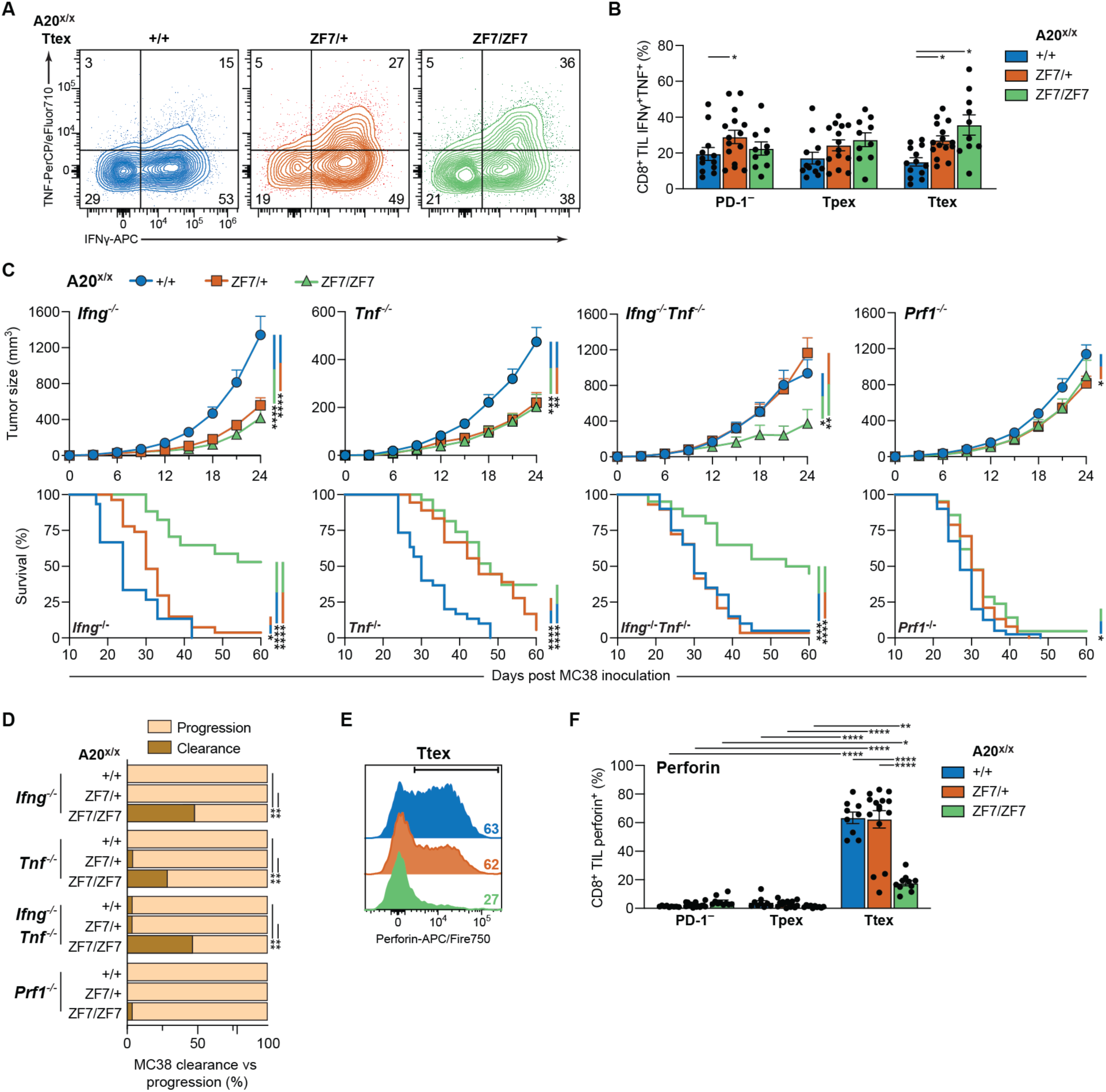
Perforin drives superior anti-tumor immunity in A20^ZF7/ZF7^ mice. **(A)** IFNγ and TNF expression by Ttex following *ex vivo* stimulation with PMA and ionomycin. Representative contour plots depict 2-4 concatenated samples per genotype from one experiment. Inset = proportion (%) of Ttex in that quadrant. **(B)** Proportion of IFNγ^+^TNF^+^ cells by CD8^+^ TIL subset. n = 10-15 mice per genotype combined from three separate experiments. **(C)** Averaged MC38 growth (top) and survival (bottom) curves from A20^ZF7^ KI mice with indicated genetic ablation of IFNγ (*Ifng^−/−^*) and/or TNF (*Tnf^−/−^*) versus perforin (*Prf1^−/−^*). Growth curves depict mean + SEM of 15-27 (*Ifng^−/−^*), 18-30 (*Tnf^−/−^*), 20-29 (*Ifng^−/−^Tnf^−/−^*), or 21-40 (*Prf1^−/−^*) mice combined from three or more separate experiments. **(D)** Proportion of mice from (C) that exhibited MC38 clearance versus those that did not (progression). Significance was assessed using Fisher’s exact test. **(E)** Ttex perforin expression in MC38 tumors. Representative FSFC histograms depicting 3-5 concatenated samples per genotype from one experiment. **(F)** Proportion of perforin-expressing cells within each CD8^+^ TIL subset. n = 9-15 mice per genotype combined from three separate experiments. **(A,B,E,F)** FSFC analyses of day 12-15 MC38 tumors. Bar graphs depict mean ± SEM. Significance was assessed using two-way ANOVA with Tukey’s multiple comparisons test. *p<0.05; **p<0.01; ***p<0.001; ****p<0.0001.

CD8^+^ TILs can also kill target cells via exocytosis of cytotoxic granules (i.e. degranulation) that contain perforin and granzymes. Perforin disrupts target cell membranes to facilitate granzyme entry, and granzymes in turn trigger cell death pathways^73^. Notably, perforin and granzymes appear to be required for tumor suppression in some tumor models, but not others^23–30^. We thus evaluated the perforin-granzyme signaling axis for its role in A20^ZF7^ anti-tumor immunity by crossing A20^ZF7^ KI mice with perforin knockout (*Prf1*^−/−^) mice. Perforin deficiency, unlike IFNγ and/or TNF deficiency, abolished the superior tumor growth control and clearance rate of A20^ZF7/ZF7^ mice (**Fig. 5C-5D**). Importantly, perforin deficiency did not alter CD8^+^ TIL density, CD8^+^ TIL subset distribution, or markers of T cell exhaustion in A20^+/+^, A20^ZF7/+^ or A20^ZF7/ZF7^ mice (**Supp. Fig. 5A-5C**). Therefore, perforin mediates enhanced anti-tumor immunity in A20^ZF7/ZF7^ mice without perturbing CD8^+^ TIL numbers or differentiation.

### A20^ZF7^ inactivation unleashes perforin degranulation by exhausted CD8^+^ T cells

The superior perforin-dependent anti-tumor cytotoxicity of A20^ZF7/ZF7^ mice could result from increased perforin expression by CD8^+^ TILs. We therefore assessed CD8^+^ TIL perforin expression by FSFC which, unlike most cytokines, can be detected without *ex vivo* stimulation. Among CD8^+^ TILs, regardless of genotype, perforin expression was largely confined to Ttex (**Fig. 5E-5F**). These findings align with prior reports of elevated perforin and granzyme levels in Ttex^2–4,62,74,75^. Surprisingly, while A20^+/+^ and A20^ZF7/+^ Ttex exhibited similarly high proportions of perforin-expressing cells, we observed an unexpected *reduction* in the numbers of perforin-expressing A20^ZF7/ZF7^ Ttex when compared to A20^+/+^ Ttex (**Fig. 5E-5F**). Perforin expression was also reduced in A20^ZF7/ZF7^ Ttex from B16F10 and CT26 tumors (**Supp. Fig. 5D**), but was not reduced in A20^OTU/OTU^ or A20^ZF4/ZF4^ Ttex from MC38 tumors (**Supp. Fig. 5E**). These findings highlight a paradox wherein biallelic A20^ZF7^ inactivation unleashes perforin-dependent anti-tumor cytotoxicity, yet engenders fewer perforin-expressing Ttex.

Diminished perforin expression by A20^ZF7/ZF7^ Ttex could result from reduced production or from enhanced degranulation. To distinguish between these possibilities, we first measured side scatter area (SSC-A) as an indicator of cellular complexity which, in the case of CD8^+^ T cells, likely reflects granule content. We observed that SSC-A was greatly increased in Ttex when compared to PD-1^−^ and Tpex subsets across genotypes (**Supp. Fig. 6A**), suggesting that the perforin-expressing Ttex subset also harbored the most granules. Accordingly, we uncovered a strong positive correlation between Ttex perforin expression and SSC-A (**Supp. Fig. 6B**). As with perforin, SSC-A was significantly reduced in A20^ZF7/ZF7^ Ttex when compared to A20^+/+^ and A20^ZF7/+^ Ttex (**Supp. Fig. 6A**), suggesting reduced cytotoxic granule content. Next, we measured the capacity of Ttex to manufacture perforin protein. We stimulated dissociated MC38 tumors *ex vivo* with phorbol 12-myristate 13-acetate (PMA) and ionomycin in the presence of brefeldin to block intracellular vesicle transport and, by extension, inhibit degranulation. Using this approach, we discovered that a similarly high fraction of A20^ZF7/ZF7^ Ttex expressed perforin as compared to A20^+/+^ and A20^ZF7/+^ Ttex (**Fig. 6A**). Thus, A20^ZF7/ZF7^ Ttex are capable of equivalent perforin production, at least in these pharmacologically stimulated cells.

**Fig. 6.**
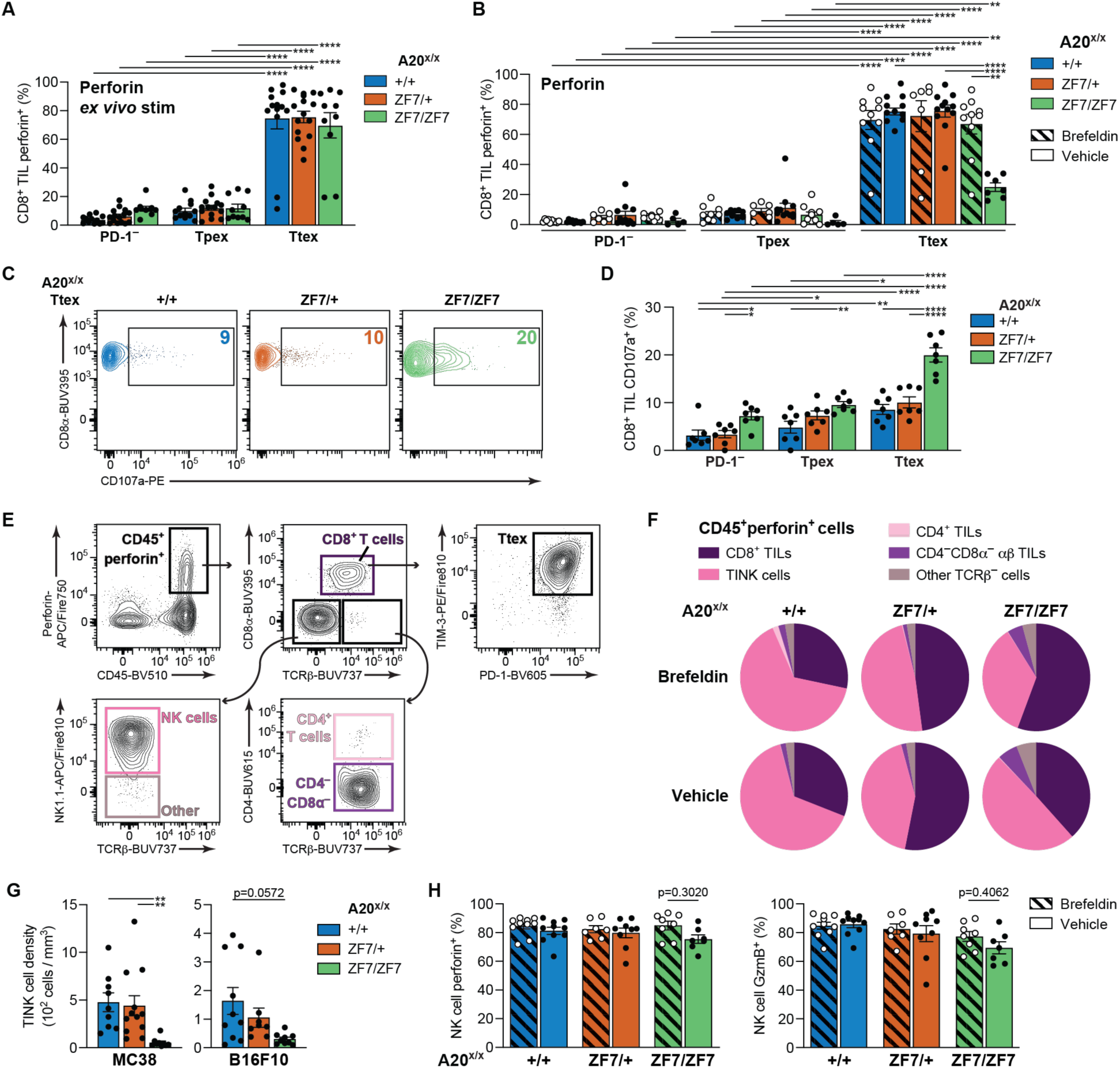
A20^ZF7^ inactivation unleashes Ttex perforin degranulation. **(A)** Perforin expression by CD8^+^ TIL subsets after 6 h *ex vivo* stimulation with PMA, ionomycin and brefeldin. n = 10-15 mice per genotype combined from three separate experiments. **(B)** Proportion of perforin-expressing CD8^+^ TILs in mice treated *in vivo* for 6 h with either brefeldin or vehicle control (5% DMSO). n = 5-12 mice per group combined from two separate experiments. **(C)** CD107a surface expression by Ttex following *ex vivo* stimulation for 1 h with PMA, ionomycin, brefeldin and monensin. Contour plots depict one representative sample per genotype from one experiment. Inset = proportion (%) of CD107a^+^ Ttex. **(D)** CD8^+^ TIL subset surface expression of CD107a, related to (C). n = 7 mice per genotype combined from two separate experiments. **(E)** Gating strategy to identify perforin-expressing cells within tumors, related to (B). Contour plots depict eight concatenated A20^+/+^, A20^ZF7/+^, and A20^ZF7/ZF7^ brefeldin-treated mice from one experiment. **(F)** Proportion of perforin-expressing cells by subset, related to (E). Pie charts depict mean for each cell subset in brefeldin-versus vehicle-treated mice from (B). **(G)** Total intratumoral TINK cell density in MC38 (left) or B16F10 (right) tumors. n = 9-13 (MC38) or 8-10 (B16F10) mice per genotype combined from two (B16F10) or three (MC38) separate experiments. **(H)** Proportion of perforin-expressing (left) or GzmB-expressing (right) TINK cells in mice treated *in vivo* for 6 h with either brefeldin or vehicle control (5% DMSO). n = 7-10 mice per group codmbined from two separate experiments. **(A-H)** FSFC analyses of day 12-15 MC38 tumors. Bar graphs depict mean ± SEM. Significance was assessed using one-way ANOVA (G) or two-way ANOVA with Tukey’s multiple comparisons test (A,B,D,H). *p<0.05; **p<0.01; ***p<0.001; ****p<0.0001.

**Supp. Fig. 5.**
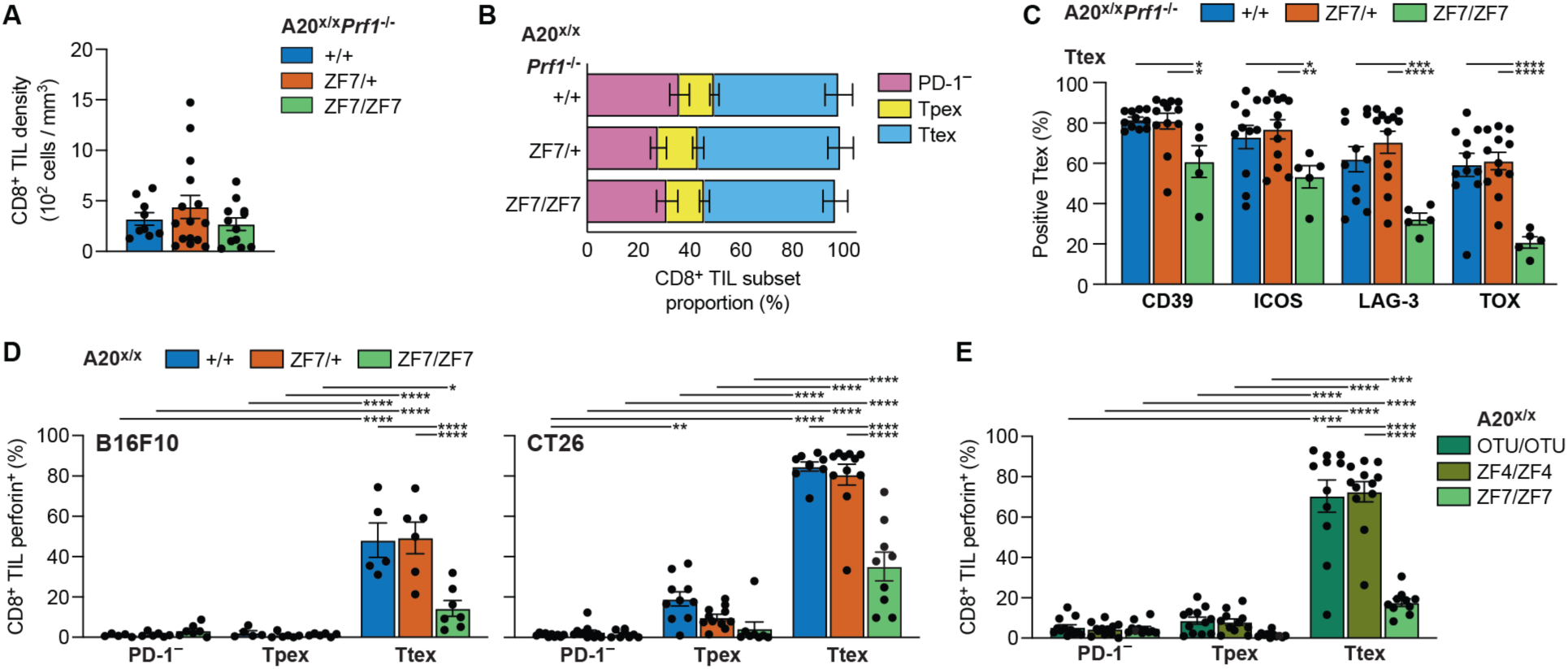
A20^ZF7/ZF7^ Ttex paradoxically exhibit reduced perforin. **(A)** Total MC38 CD8^+^ TIL density in *Prf1^−/−^* mice. n = 9-15 mice per genotype combined from three separate experiments. **(B)** CD8^+^ TIL subset proportions from (A). **(C)** Ttex IR and TOX expression from (B). **(D)** Perforin expression by CD8^+^ TIL subsets within B16F10 (left) or CT26 (right) tumors. **(E)** Perforin expression by CD8^+^ TIL subsets in MC38 tumors. A20^ZF7/ZF7^ data is repeated from Fig. 5F for comparison. **(A-E)** FSFC analyses of day 12-15 tumors. Bar graphs depict mean + SEM. Significance was assessed using one-way ANOVA (A) or two-way ANOVA with Tukey’s multiple comparison’s test (B-E). *p<0.05; **p<0.01; ***p<0.001; ****p<0.0001.

**Supp. Fig. 6.**
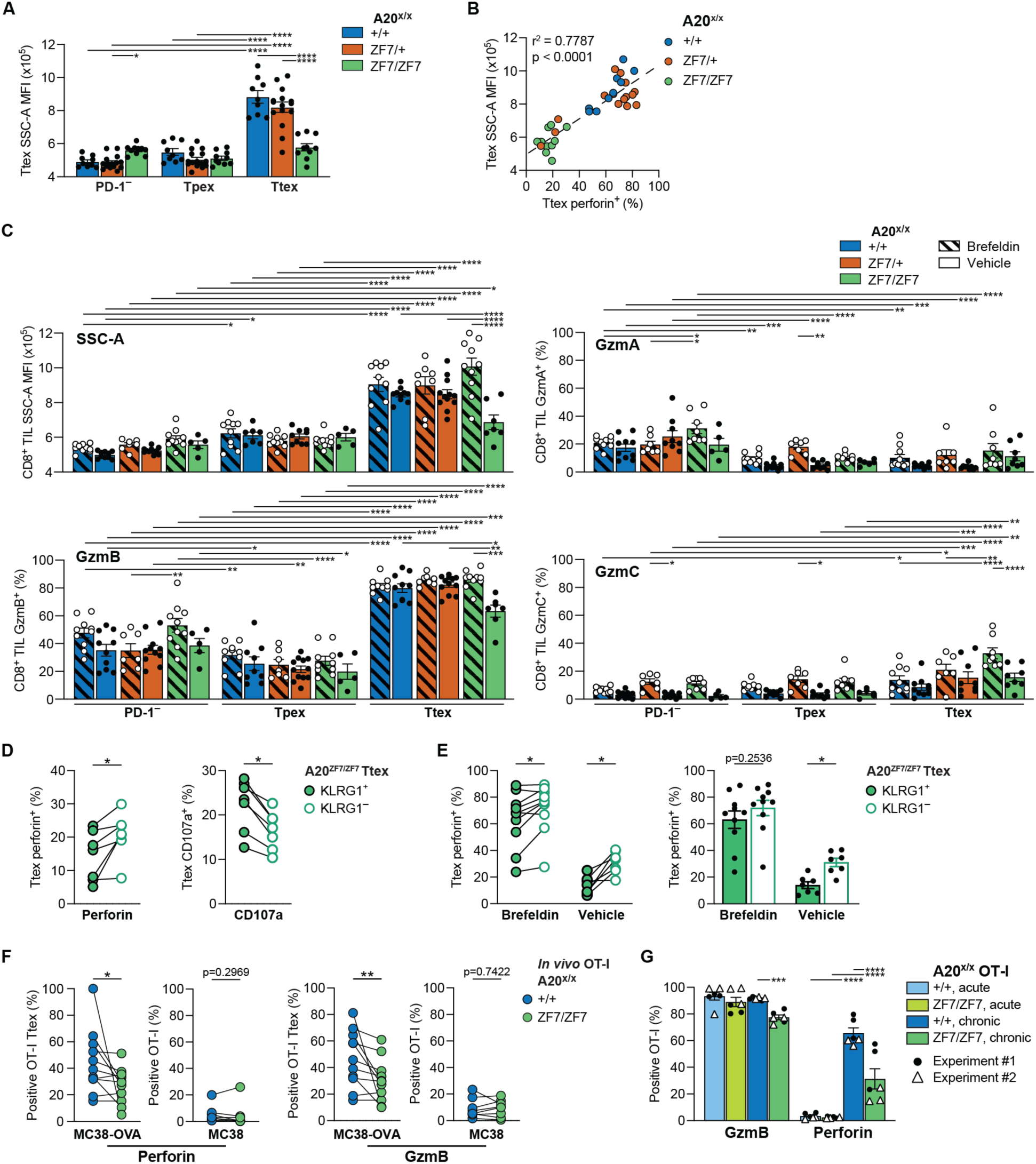
A20^ZF7^ suppression of Ttex degranulation depends upon repetitive TCR stimulation. **(A)** CD8^+^ TIL subset SSC-A mean values. n = 9-15 samples per genotype combined from three separate experiments. **(B)** Scatter plot correlating SSC-A mean from (A) with Ttex perforin expression (see Fig. 5F). Statistics were calculated using simple linear regression (dashed line). **(C)** SSC-A mean (top left) or proportion of GzmA- (top right) versus GzmB- (bottom left) versus GzmC- (bottom right) expressing CD8^+^ TIL subsets in mice treated *in vivo* for 6 h with either brefeldin or vehicle control (5% DMSO). n = 5-12 mice per group combined from two separate experiments. **(D)** Paired analysis of perforin (left; related to Fig. 5F) or CD107a (right; related to Fig. 6D) expression by KLRG1^+^ versus KLRG1^−^ A20^ZF7/ZF7^ Ttex. **(E)** (Left) Paired analysis of perforin expression by KLRG1^+^ versus KLRG1^−^ A20^ZF7/ZF7^ Ttex in mice treated for 6 h *in vivo* with either brefeldin or vehicle control, related to Fig. 6B. (Right) Same data presented in unpaired format. **(F)** Paired analysis of perforin (left) or GzmB (right) expression by OT-I Ttex 7 days following adoptive co-transfer into MC38-versus MC38-OVA-bearing WT mice. **(G)** GzmB and perforin expression by OT-I cells following *in vitro* acute versus chronic/repetitive stimulation. Bar graphs depict mean ± SEM of technical triplicates combined from two separate experiments. Significance was assessed using two-way ANOVA with Bonferroni’s multiple comparisons test. **(A-F)** FSFC analyses of day 12-15 MC38 tumors. Bar graphs depict mean ± SEM, with significance assessed using Bonferroni’s (E) or Tukey’s multiple comparisons test (A,C). Symbol/line pairs represent cells matched from same tumor, with significance assessed using Wilcoxon matched-pairs signed rank test. *p<0.05; **p<0.01; ***p<0.001; ****p<0.0001.

To quantify Ttex perforin content in their native TME without additional stimulation, we treated MC38-bearing mice for 6 hours *in vivo* with brefeldin or vehicle control^76^, then analyzed tumors by FSFC. This experiment revealed several important facets of perforin degranulation by A20^+/+^ and A20^ZF7^ KI CD8^+^ TILs. Most strikingly, brefeldin treatment led to a marked increase in the fraction of perforin^+^ A20^ZF7/ZF7^ Ttex when compared to vehicle treatment, rising to levels that matched those of A20^+/+^ and A20^ZF7/+^ Ttex (**Fig. 6B**), indicating that many A20^ZF7/ZF7^ Ttex are likely degranulating perforin in the absence of brefeldin. Accordingly, surface levels of the degranulation marker CD107a were significantly higher on A20^ZF7/ZF7^ Ttex than on A20^+/+^ or A20^ZF7/+^ Ttex following brief *ex vivo* stimulation (**Fig. 6C-6D**). Meanwhile, the fraction of perforin^+^ Ttex remained similar in brefeldin-versus vehicle-treated A20^+/+^ and A20^ZF7/+^ mice (**Fig. 6B**), suggesting minimal degranulation by these Ttex. Parallel findings of increased SSC-A, GzmB and GzmC were observed in brefeldin-treated A20^ZF7/ZF7^ Ttex, but not in A20^+/+^ or A20^ZF7/+^ Ttex (**Supp. Fig. 6C**). Of note, minimal GzmA expression was observed in Ttex of any genotype (**Supp. Fig. 6C**). Importantly, while PD-1^−^ TILs and Tpex both expressed GzmB (**Supp. Fig. C**), few perforin^+^ PD-1^−^ TILs or Tpex were detected in any genotype even after *in vivo* brefeldin treatment (**Fig. 6B**), confirming that perforin production is highly restricted to Ttex. Finally, KLRG1^+^ Ttex at baseline expressed less perforin and more CD107a than KLRG1^−^ Ttex (**Supp. Fig. 6D**). After *in vivo* brefeldin treatment, perforin levels between KLRG1^+^ and KLRG1^−^ A20^ZF7/ZF7^ Ttex became relatively similar (**Supp. Fig. 6E**), suggesting that the less exhausted appearing KLRG1^+^ Ttex subset was more actively degranulating perforin *in vivo*. Overall, these data highlight an unappreciated defect in Ttex degranulation that can be reversed upon A20^ZF7^ inactivation.

We next aimed to verify that Ttex-derived perforin – rather than perforin produced by another cell subset – drove the A20^ZF7/ZF7^ anti-tumor phenotype. We gated the FSFC data from our *in vivo* brefeldin-versus vehicle-treated mouse experiments on perforin^+^ cells and discovered that, as expected, almost all were CD45^+^ leukocytes (**Fig. 6E**). Among these CD45^+^perforin^+^ cells, nearly all were either CD8^+^ TILs (primarily Ttex) or tumor-infiltrating NK (TINK) cells, regardless of genotype (**Fig. 6F**). Three key pieces of evidence indicated that TINK cell-derived perforin was not the key driver of anti-tumor immunity in A20^ZF7/ZF7^ mice. First, as discussed, RAG1 deficiency, which ablates CD8^+^ TILs but not TINK cells, eliminated the anti-tumor phenotype (**Supp. Fig. 3A-3B**). Second, we observed significantly diminished TINK cell density in A20^ZF7/ZF7^ mice when compared to A20^+/+^ mice (**Fig. 6G**), implicating a key role for A20 in supporting TINK cell proliferation, survival or recruitment. Third, A20^ZF7/ZF7^ TINK cells in vehicle-treated mice expressed similarly high levels of perforin and GzmB as A20^+/+^ and A20^ZF7/+^ TINK cells, with no further increase following *in vivo* brefeldin treatment (**Fig. 6H**), suggesting that A20^ZF7/ZF7^ TINK cells are not robustly degranulating. Together these findings validate a critical anti-tumor role for A20^ZF7/ZF7^ Ttex-derived perforin, and highlight different roles for A20 in regulating CD8^+^ TIL versus TINK cell biology.

### Cas9-editing of A20^ZF7^ in human CAR-T cells enhances anti-tumor immunity *in vivo*

Given the potent ability of A20^ZF7^ to regulate anti-tumor immunity, we next tested whether specific ablation of the A20^ZF7^ motif in human CD19 *TRAC*-CAR-T cells would enhance leukemia control in preclinical models. We designed a Cas9 sgRNA targeting a double-stranded break (DSB) at Asp768 in the A20^ZF7^ motif to disrupt the domain via indel frameshift (Supp. Table 6), thereby inactivating the two (of four) most C-terminal zinc coordinating cysteines in the A20^ZF7^ domain that instruct M1 Ub binding^12^. This Cas9-directed edit at Asp768 is near the C-terminus of A20, and is therefore not expected to impact the functionality of the remainder of the A20 protein. We confirmed efficient A20^ZF7^ indel generation (91%), and immunoblotting revealed efficient deletion of A20 in A20^KO^ T cells, while A20^ZF7^ T cells retained A20 protein expression (**Supp. Fig. 7A**). We next injected A20^KO^, A20^ZF7^, or control AAVS1 CD19 CAR-T cells into NALM6-bearing NSG mice and monitored leukemia growth. Mice treated with either A20^KO^ or A20^ZF7^ CAR-T cells potently suppressed NALM6 leukemia growth and demonstrated increased complete response (CR) rates (**Fig. 7A; Supp. Fig. 7B**) and survival (**Fig. 7B**) when compared to mice treated with control AAVS1 CAR-T cells. Most recipients of A20^KO^ or A20^ZF7^ CAR-T cells that achieved CR remained leukemia-free for several months without relapse, and furthermore resisted iterative NALM6 rechallenge (**Supp. Fig. 7C**). Thus, as with A20^KO^ CAR-T cells, Cas9-directed abrogation of the A20^ZF7^ domain in CAR-T cells dramatically enhances the anti-tumor efficacy and functional persistence of these cells.

**Fig. 7.**
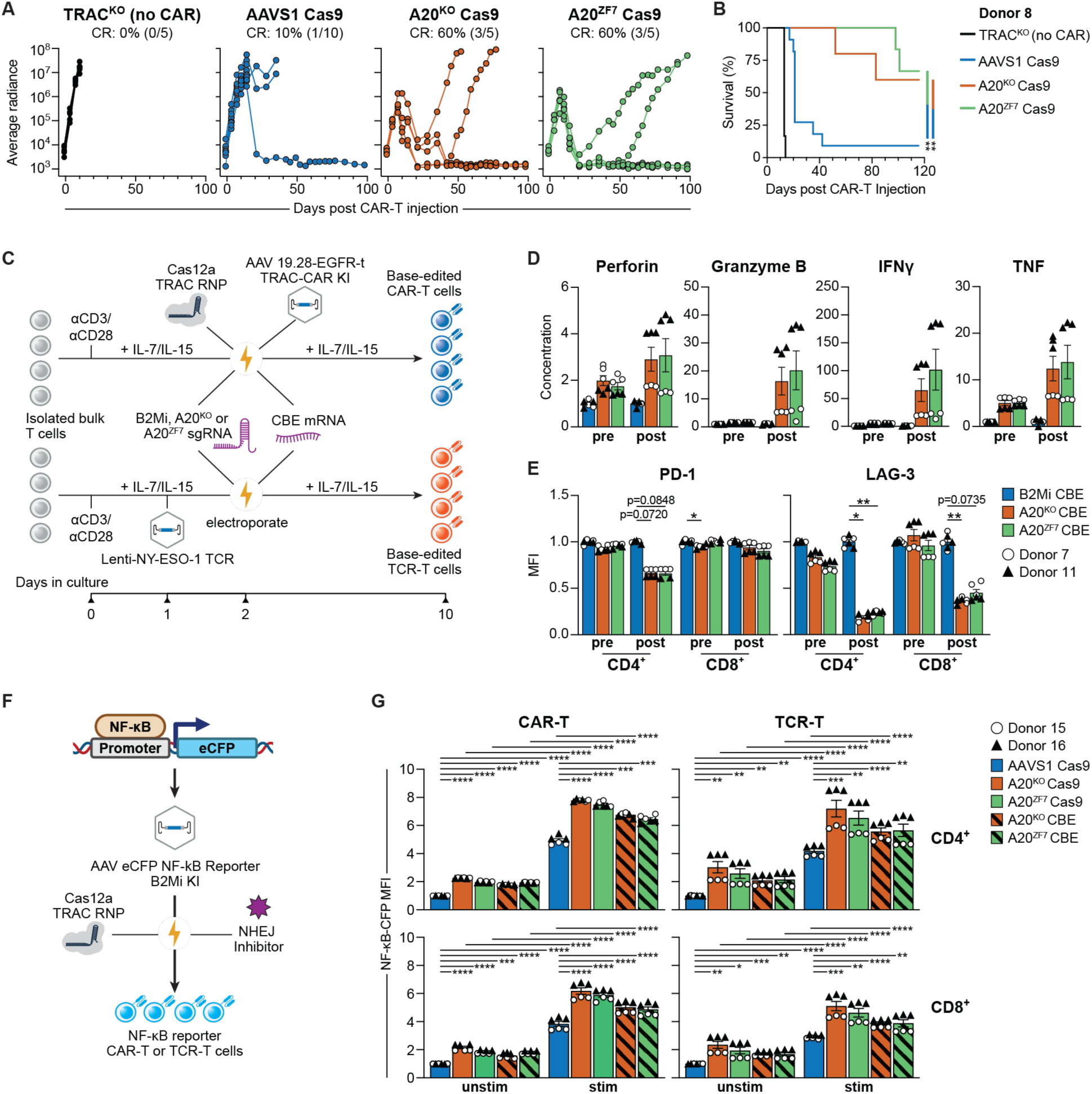
Base-editing to inactivate A20^ZF7^ enhances CAR-T cell function *in vitro*. **(A)** Individual NALM6 growth curves of mice treated with Cas9-edited CAR-T cells or with TRAC^KO^ (no CAR) T cells (n = 1 T cell donor, n = 5-10 mice per group). CR = complete response. Leukemia burden was quantified via BLI. Radiance units = photons/s/cm^2^/steradian. **(B)** Survival analysis corresponding to (A). Significance was assessed using log-rank test. **(C)** Schematic illustration for the generation of base-edited CD19 *TRAC*-CAR-T (top) or 1G4 TCR-T (bottom) cells. Cas12a RNP carrying *TRAC*-targeting sgRNAs (CAR-T only), cytosine base-editor (CBE)-encoding mRNA, and a CBE sgRNA were introduced into T cells via electroporation. **(D)** Normalized supernatant concentrations (pg/ml) of secreted effector molecules from base-edited CD19 CAR-T cells, as determined by LEGENDplex. CAR-T cells were analyzed before (pre) or after (post) 3 rounds of repetitive stimulation. **(E)** Normalized IR expression by CD4^+^ and CD8^+^ CD19 CAR-T cells either before (pre) or after (post) 5 rounds of repetitive stimulation. **(F)** Schematic illustration for the generation of NF-κB activity reporter CAR-T or TCR-T cells. Cells were electroporated with Cas12a RNP carrying B2Mi-targeting sgRNAs, then transduced with AAV carrying NF-κB reporter construct. For base-edited cells, steps in (C) were concurrently performed. For Cas9-edited cells, Cas9 RNPs with sgRNAs targeting AAVS1, A20^KO^ or A20^ZF7^ were electroporated in lieu of CBE mRNA and associated sgRNAs. **(G)** Normalized NF-kB-CFP MFI of Cas9 or base-edited CD19 CAR-T (left) or 1G4 TCR-T (right) cells before (unstim) or after 12 h stimulation (stim) with antigen-expressing tumor cells (E:T 1:2). **(D,E,G)** Bar graphs depict mean ± SEM of n = 2 donors with n = 3 technical replicates per donor. Each technical replicate was normalized to mean of control (AAVS1 or B2Mi) group for corresponding donor and condition. Significance was assessed using paired t test. *p<0.05; **p<0.01; ***p<0.001; ****p<0.0001.

### Base-editing of A20^ZF7^ enhances cytolytic function of human CAR-T cells *in vitro*

Cas9-based CRISPR editing relies upon DNA DSBs that lead to deletions or frameshifts. By contrast, CRISPR base editors (BEs) generate targeted base pair edits without causing DSB^77^. Hence, BE-directed mutagenesis usually mutates a single amino acid and avoids triggering DNA DSB repair pathways, which altogether confers a more defined and clinically safer approach to genomic engineering. To test the therapeutic potential of this strategy, we used BEs to introduce a targeted A20^ZF7^ domain missense mutation into CAR-T cells. From our ESM1b analysis and large-scale BE screen (**Fig. 2A**),^7^ we nominated a specific amino acid residue within the A20^ZF7^ motif (Cys779), and a corresponding sgRNA targeting a missense mutation (C779Y; Supp. Table 6) predicted to disrupt A20 function and enhance T cell activation (**Supp. Fig. 7D**). Notably, Cys779 represents the third (of four) Zn^++^-coordinating cysteines in the A20^ZF7^ motif, and is likely important for M1 Ub binding^55,56,78^. For comparison, we also generated base-edited A20^KO^ CAR-T cells using an sgRNA that introduces a premature stop codon in exon 2 of the A20 gene (R90*; Supp. Table 6), and confirmed elimination of the A20 protein by immunoblot (**Supp. Fig. 7A**). Control T cells from the same donors were edited with an sgRNA targeting an intronic sequence of the *B2M* gene encoding beta-2 microglobulin (β2m; Supp. Table 6). This intronic mutation (hereafter referred to as B2Mi) does not disrupt β2m surface protein expression (**Supp. Fig. 7E**). Next-generation sequencing analysis of T cells collected 7 days after editing confirmed that both the A20^KO^ and A20^ZF7^ sgRNAs edited A20 with 90% efficiency, generating the expected R90* or C779Y mutations, while the B2Mi sgRNA edited with 97% efficiency.

We next engineered CD19 CAR-T cells that were base-edited with either A20^KO^, A20^ZF7^ or control B2Mi sgRNAs. We introduced Cas12a RNP carrying *TRAC*-targeting sgRNAs, cytosine base-editor (CBE)-encoding mRNA, and a CBE sgRNA targeting either A20^KO^, A20^ZF7^ or B2Mi into T cells via electroporation (**Fig. 7C**). As CBE is coupled to a nicking Cas9 (nCas9) for targeting, we utilized Cas12a-mediated *TRAC*-CAR knock-in here to prevent guide swapping between the two Cas proteins^79^. We then tested our base-edited CD19 CAR-T cells using our *in vitro* repetitive stimulation assay. Base-edited CAR-T cells were repeatedly stimulated with fresh CD19^+^ tumor cells, then evaluated for expression of activation and exhaustion markers, production of cytokines and cytotoxic proteins, and tumor cell killing capacity. Repetitively stimulated A20^KO^ and A20^ZF7^ CAR-T cells exhibited enhanced proliferation when compared to control B2Mi CAR-T cells (**Supp. Fig. 7F**). Remarkably, A20^KO^ and A20^ZF7^ base-edited CAR-T cells also maintained robust tumoricidal capacity compared to control B2Mi CAR-T cells after three rounds of tumor stimulation (**Supp. Fig. 7G**). Furthermore, compared to control B2Mi CAR-T cells, base-edited A20^KO^ and A20^ZF7^ CAR-T cells produced more effector cytokines (IFNγ, TNF) and secreted substantially more cytotoxic granule proteins (perforin, GzmB) after repetitive stimulation (**Fig. 7D**), consistent with our findings of increased secretion by Cas9-edited A20^KO^ TCR-T cells (**Fig. 1D**) and increased degranulation by A20^ZF7/ZF7^ Ttex (**Fig. 6**). Lastly, repetitively stimulated A20^KO^ and A20^ZF7^ CAR-T cells expressed lower levels of exhaustion markers (PD-1 and LAG-3; **Fig. 7E**) and higher levels of activation markers (CD25, CD137, CD69 and CD154) (**Supp. Fig. 7H**). Importantly, by all metrics, A20^KO^ and A20^ZF7^ CAR-T cells performed similarly.

**Supp. Fig. 7.**
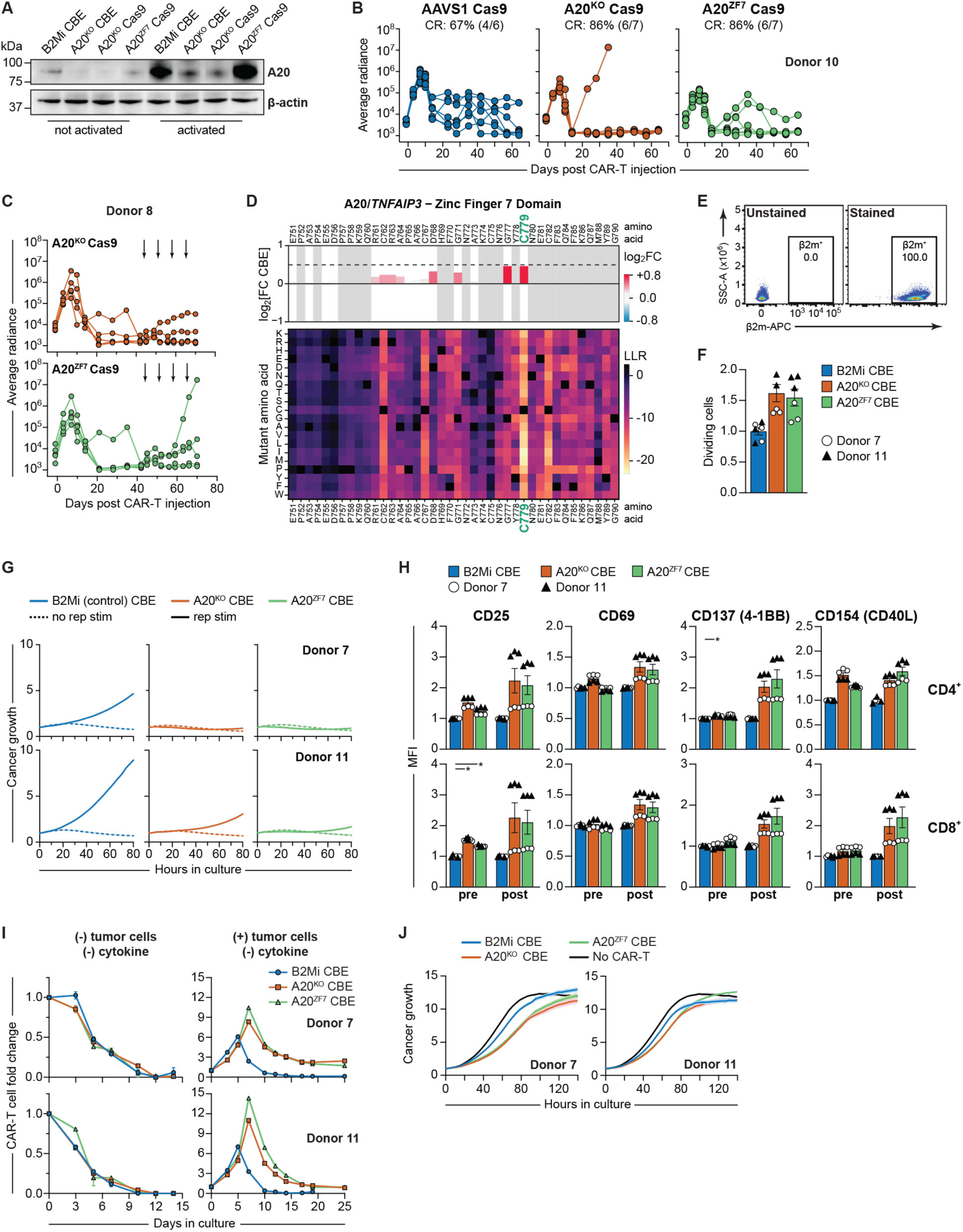
Cas9- or BE-directed A20^ZF7^ inactivation augments CAR-T cell anti-tumor activity without safety concerns. **(A)** Immunoblot of A20 expression by Cas9- or base-edited (CBE) human T cells that were either not activated, or activated with anti-CD3/CD28 for 24 hours. n = 1 donor. **(B)** Individual NALM6 growth curves of mice treated with Cas9-edited CAR-T cells or with TRAC^KO^ (no CAR) T cells (n = 1 T cell donor different from Fig. 7A, n = 6-7 mice per group). CR = complete response. Leukemia burden was quantified via BLI. Radiance units = photons/s/cm^2^/steradian. **(C)** Individual NALM6 growth curves of mice that controlled initial challenge (Fig. 7A) and underwent iterative rechallenge (black arrows). **(D)** CBE screen (top) and ESM1b heatmap (bottom) from Fig. 2A, zoomed in on A20^ZF7^ motif. sgRNA target (Cys779) highlighted in green. **(E)** Representative dot plots depicting β2m expression on B2Mi CAR-T cells 36 hours after base-editing. Inset = percentage of β2m^+^ cells. **(F)** Normalized dividing cells (CellTrace Violet^lo^), as determined via proliferation assay of CAR-T cells after 72 h of stimulation. **(G)** Normalized growth curves of A375-CD19 cells. CAR-T cells either without prior repetitive stimulation (no rep stim) or after 3 rounds of repetitive stimulation (rep stim) were co-cultured with A375-CD19 cells at an E:T ratio of 1:1 (mean ± SEM of n = 3 technical replicates; n = 2 donors). **(H)** Normalized expression of activation markers by CD4^+^ (top) and CD8^+^ (bottom) CAR-T cells before (pre) and after (post) 3 rounds of tumor stimulation. **(I)** Fold change of base-edited CD19 CAR-T cells cultured without cytokine either in the absence (left) or presence (right) of A375-CD19 target cells (E:T ratio 1:1). **(J)** Normalized growth curves of CD19-negative A375 melanoma cells co-cultured with CD19 CAR-T cells (E:T 1:1), or cultured without CAR-T cells. **(F,H)** Bar graphs depict mean ± SEM of n = 2 donors with n = 3 technical replicates per donor. Each technical replicate was normalized to mean of B2Mi group for corresponding donor and condition. Significance was assessed using paired t test. **(I,J)** All curves show mean ± SEM of n = 3 technical replicates from n = 2 T cell donors. *p<0.05.

We next determined whether these CRISPR editing approaches enhanced NF-κB activity, a common readout for A20 dysfunction^51,56^. To this end, we generated AAVS1 and Cas9- or CBE-directed A20^KO^ and A20^ZF7^ TCR-T or CAR-T cells, and concurrently transduced them with a construct encoding NF-κB-dependent cyan fluorescent protein (CFP) expression, knocked into the B2Mi locus to ensure consistent regulation of CFP expression across cells (**Fig. 7F**). Notably, both Cas9 and CBE A20^KO^ and A20^ZF7^ CAR-T and TCR-T cells exhibited higher basal NF-κB activity than control AAVS1 cells even prior to stimulation (**Fig. 7G**). Following stimulation with antigen-expressing tumor cells, all groups expectedly increased their NF-κB activity, but both Cas9- and base-edited A20^KO^ and A20^ZF7^ CAR-T and TCR-T cells demonstrated substantially higher activity than AAVS1 (**Fig. 7G**). Thus, functional ablation of A20 and augmentation of NF-κB activity within ACTs can be easily achieved using Cas9- or CBE-directed disruption of the A20^ZF7^ motif.

We next tested whether A20 ablation via base-editing enables human CAR-T cells to grow in a cytokine- or antigen-independent fashion, which could represent a safety challenge for ACT applications. First, we evaluated the growth of A20^KO^ and A20^ZF7^ CAR-T cells after the withdrawal of cytokines, where we observed consistent decreases in cell numbers and viability of both A20^KO^ and A20^ZF7^ CAR-T cells, similar to control B2Mi CAR-T cells (**Supp. Fig. 7I**). Further, the numbers of A20^KO^ and A20^ZF7^ CAR-T cells contracted as expected after initial exposure to tumor cells without cytokines (**Supp. Fig. 7I**). We also assessed the level of dependence on antigen of A20^KO^ and A20^ZF7^ CAR-T cells for tumor cell killing. Here, we exposed all CAR-T cell conditions to CD19-negative tumor cells and assessed their ability to eradicate tumor cells over several days. We observed a small degree of allogenic killing, which was similar across all conditions (**Supp. Fig. 7J**). Therefore, targeting A20 does not confer cytokine-independent growth or antigen-independent killing, suggesting A20-targeted CAR-T cells may be safe for clinical use.

### Base-editing of A20^ZF7^ enhances human CAR-T cell efficacy *in vivo*

To test the ability of base-edited A20-targeted CAR-T cells to suppress tumor growth *in vivo*, we treated NALM6-engrafted mice with either base-edited A20^KO^, A20^ZF7^, or control B2Mi CD19 CAR-T cells and serially measured growth of the luminescent NALM6 tumors via BLI. Mice treated with A20^KO^ or A20^ZF7^ base-edited CAR-T cells exhibited superior tumor control, CR rates and survival when compared to mice treated with B2Mi CAR-T cells (**Fig. 8A-8B; Supp. Fig. 8A-8B**). Importantly, mice that controlled their initial leukemia challenge appeared grossly normal based upon visual inspection and body weight (**Supp. Fig. 8C**). To understand why A20^KO^ and A20^ZF7^ engineered CAR-T cells suppress leukemia growth better than control B2Mi CAR-T cells, we evaluated bone marrow samples from tumor bearing mice 14 days after CAR-T cell adoptive transfer by flow cytometry. At this early time point, leukemia burden remained relatively even across treatment groups. Our findings showed a trend toward higher numbers of CAR-T cells in the bone marrow of mice treated with A20^KO^ or A20^ZF7^ CAR-T cells when compared to those treated with control B2Mi CAR-T cells (**Supp. Fig. 8D**). Notably, editing of A20 did not substantially change the distribution of memory phenotypes of the isolated CAR-T cells (**Supp. Fig. 8E**). Finally, the frequency of CAR-T cells co-expressing 2 or more IRs was markedly lower in A20^KO^ (24-30%) and A20^ZF7^ (34-46%) CAR-T cells when compared with B2Mi control (51-65%) (**Fig. 8C**), consistent with our findings in A20^ZF7/ZF7^ Ttex (**Fig. 4**). These data highlight that base-editing of A20^ZF7^ can lead to remarkable improvements *in vivo* using an already optimized best-in-class CAR-T cell therapy, accompanied by reduced exhaustion, without any clear safety risk.

**Fig. 8.**
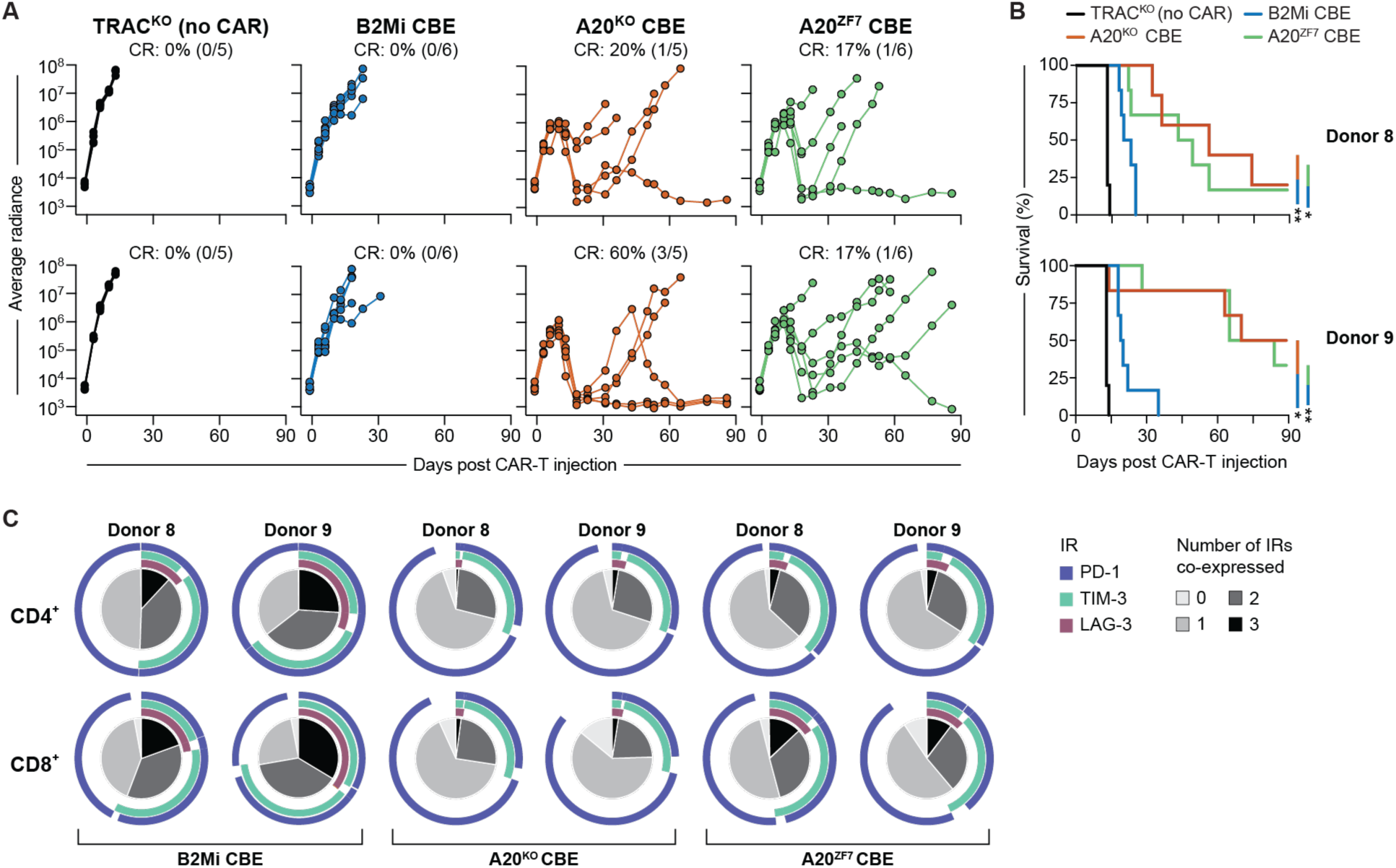
Base-editing to inactivate A20^ZF7^ enhances CAR-T cell function *in vitro* and *in vivo*. **(A)** Individual NALM6 growth curves of mice treated with base-edited (CBE) CAR-T cells, or with TRAC^KO^ (no CAR) T cells (n = 2 T cell donors, n = 5-6 mice per group per donor). CR = complete response. Leukemia burden was quantified via BLI. Radiance units = photons/s/cm^2^/steradian. **(B)** Survival analysis corresponding to (A). Survival significance was assessed using log-rank test. **(C)** SPICE analyses of IR co-expression on base-edited CD4^+^ and CD8^+^ CAR-T cells isolated from bone marrow of NALM6-bearing mice 14 days after CAR-T cell injection, as determined by flow cytometry. Plots depict mean of n = 3 mice per group per n = 2 donors. *p<0.05; **p<0.01.

**Supp. Fig. 8.**
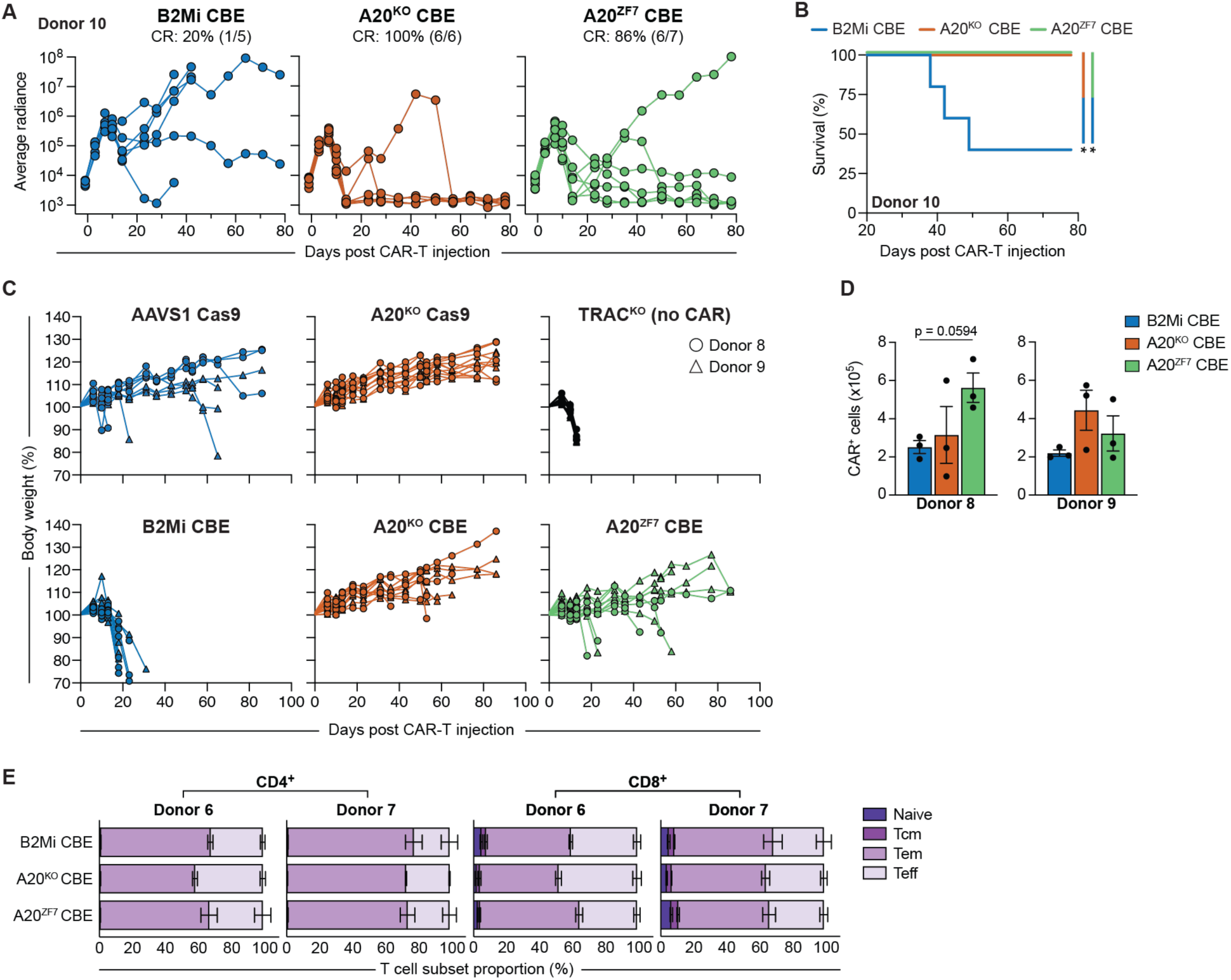
BE-directed A20^ZF7^ inactivation augments CAR-T cell anti-tumor activity. **(A)** Individual NALM6 growth curves of mice treated with Cas9-edited CAR-T cells or with TRAC^KO^ (no CAR) T cells (n = 1 T cell donor, n = 5-7 mice per group). Different donor than used in Fig. 8A. Leukemia burden was quantified via BLI. Radiance units = photons/s/cm^2^/steradian. **(B)** Survival analysis corresponding to (A). Survival significance was assessed using log-rank test. **(C)** Total body weight of NALM6-bearing mice after receiving Cas9- or base-edited CAR-T cell injection, expressed as percent change over baseline weight (measured on day −1 prior to CAR-T injection). Each graph depicts individual weight curves of n = 5-9 mice per donor per group (n = 2 T cell donors). **(D)** Absolute numbers of CAR-T cells harvested from the bone marrow of NALM6-bearing mice 14 days after CAR-T cell injection, as determined by flow cytometry (mean ± SEM, n = 2 T cell donors, n = 3 mice per group per donor). Significance was assessed using one-way ANOVA with Dunnett’s T3 multiple comparisons test. **(E)** Differentiation status of CD4^+^ or CD8^+^ base-edited CAR-T from (e). Naive = CD45RA^+^CD62L^+^; central memory (Tcm) = CD45RA^−^CD62L^+^; effector memory (Tem) = CD45RA^−^CD62L^−^; effector (Teff) = CD45RA^+^CD62L^−^ (mean ± SEM, n=2 T cell donors, n = 3 mice per group per donor). **(D,E)** Bar graphs depict mean ± SEM. *p<0.05.

## Discussion

In this study, we filtered the entire protein-coding genome through a CRISPR knockout screen, followed by a base-editing mutagenesis screen, a protein large language model, and transgenic knock-in mice to pinpoint a single protein domain — A20^ZF7^ — as a critical negative negative regulator of CD8^+^ TIL function. This precise molecular lever operated both in murine and human T cells, in liquid and solid tumors, and in immunodeficient and immunocompetent mice. We then base-edited a single A20^ZF7^ missense mutation at high editing efficiency that robustly enhanced CAR-T cell anti-tumor activity. Our study sets a new standard for engineering functionally enhanced CAR-T cell clinical products using base-editing. BE-directed gene ablation avoids Cas9-directed DNA double-stranded breaks and their potential associated chromosomal rearrangements^77,80^. We take this approach one step further by using base-editing to introduce a strategic missense mutation rather than total gene ablation. For multifunctional proteins like A20, this approach could prove superior to complete ablation, with fewer undesirable side effects. Recent large-scale base-editing mutagenesis screens have begun to pinpoint key amino acid residues that impact T cell functions *in vitro*^7,8^. We have now translated this approach to CAR-T cell enhancement *in vivo* by targeting A20^ZF7^. BE-guided gene ablation can also be multiplexed^81^, thus setting the stage for combining missense mutations to develop safer and more efficacious next-generation ACTs. Indeed, multiplexed editing within A20 itself might lead to further enhancements, as we and others have described synergy between the A20^ZF4^ and A20^ZF7^ motifs that prevents spontaneous inflammation^12,13^.

Interestingly, the aforementioned large-scale BE screens to identify amino acids relevant to T cell function converged on mutations found in hereditary autoimmune syndromes^7,8^. Likewise, inherited A20 mutations cause haploinsufficiency of A20 (HA20), a syndrome with autoimmune manifestations^82^. Also of note, T cell lymphoma oncogenic mutations intentionally expressed in CAR-T cells can boost their efficacy, particularly those that augment NF-κB signaling^83^, and somatic A20 mutations have been described in B and T cell lymphomas^84,85^. Altogether, these findings highlight that the optimal enhancement of ACTs may already be written in the genetics of existing pathophysiologic processes, including those tied to A20 dysfunction. Importantly, our studies using A20^KO^ or A20^ZF7^ CAR-T cells showed no evidence of dysregulated proliferation *in vitro* or *in vivo*. While continued safety monitoring will be paramount as A20-deficient ACTs move forward toward clinical use, preliminarily we find no safety concerns.

Our prior genome-scale CRISPR screens identified a role for A20 in restraining proliferation of acutely stimulated human T cells^31,33^. We herein leveraged this platform to reveal that A20 potently orchestrates T cell dysfunction in the setting of chronic antigen exposure. Our data extend the findings of previous genome-scale human T cell exhaustion screens^5,6^ by incorporating substantially longer repetitive tumor stimulation. In our screen, A20 became progressively enriched with increasing rounds of stimulation, suggesting that A20’s role is potentiated during later stages of exhaustion. These results align with multiomics analyses wherein Ttex differentiation correlated with increased A20/*Tnfaip3* chromatin accessibility and decreased NF-κB transcriptional signatures^86,87^. The mechanism by which A20 promotes terminal exhaustion has yet to be elucidated, though our findings suggest that A20 supports TOX expression. Indeed, the predominance of KLRG1^+^ A20^ZF7/ZF7^ Ttex mirrors the skewing of chronically stimulated TOX-deficient CD8^+^ T cells toward KLRG1^+^ cells^65,67^. A20 might also promote exhaustion via restraint of NF-κB signaling. In acutely activated T cells, A20 disrupts NF-κB activity following recruitment to the CARD11-BCL10-MALT1 (CBM) signalosome^14,15^, possibly via the A20^ZF7^ motif^17^. Acutely activated CAR-T cells also recruit A20 to the CAR endodomain^88^. Repeated TCR or CAR stimulation might similarly engage A20 to repress NF-κB and induce TOX. However, unlike A20^ZF7^ CAR-T cells and A20^ZF7/ZF7^ Ttex, TOX-deficient CD8^+^ TILs remain dysfunctional in the TME^66^. Thus, A20 likely also regulates TOX-independent pathways to implement exhaustion.

We reveal two layers of anti-tumor effector mechanisms unlocked by targeting A20^ZF7^ in T cells and CAR-T cells. The first layer involves enhanced production of IFNγ and TNF, consistent with prior studies of A20^KO^ CD8^+^ TILs^20–22^. We extend these studies by directly interrogating the role of IFNγ and TNF in anti-tumor immunity, and by showing that inactivation of a single A20^ZF7^ allele promotes cytokine-dependent tumor suppression. However, A20^ZF7/ZF7^ mice suppressed tumor growth even in the absence of these cytokines, highlighting a more potent second layer — the perforin-granzyme axis — accessible only below a certain threshold of A20^ZF7^ activity. While our understanding of Tex biology to date has concentrated upon epigenetic and transcriptional regulation of effector programs, we discover an unappreciated posttranscriptional regulatory step in Ttex at the level of perforin degranulation. Indeed, Ttex consistently express perforin and granzymes^2–4^, yet their native ability to degranulate has not been studied. We used *in vivo* brefeldin treatment to demonstrate that A20^ZF7/ZF7^ Ttex rapidly degranulated perforin, while A20^+/+^ and A20^ZF7/+^ Ttex did not. Thus, assays that solely measure Ttex perforin or granzyme content may not accurately reflect their true cytotoxic potential. Moving forward, the use of *in vivo* brefeldin treatment could help discriminate the functional state of Ttex. While the pathways that connect TCR engagement to degranulation remain incompletely understood^89^, genome-scale CRISPR screens have begun to discover mediators in acutely activated T cells^90^. Similar screens in chronically stimulated T cells might find additional mediators of Ttex degranulation beyond A20. Importantly, the dependence on cytokines versus perforin for anti-tumor immunity may vary based on the tumor model or immunotherapy used^23–30^. Our study emphasizes the importance of finding targets such as A20^ZF7^ that can enhance TIL and CAR-T cell activity in multiple ways.

## Resource Availability

Further information and requests for resources and reagents should be directed to and will be fulfilled by the lead contact, Averil Ma.

Unique and stable reagents generated in this project are available upon request. Material transfer agreements for these reagents may be required.

Flow cytometry and other standard immunological data will be deposited at ImmPort Shared Data (https://import.org).

This paper also analyzes existing, publicly available data, accessible at https://www.ncbi.nlm.nih.gov/geo/.

Other data reported in this paper will be shared by the lead contact upon request. This paper does not report original code.

## Acknowledgements

We thank Stacie Dodgson (Gladstone Institutes) for advice regarding manuscript preparation, Tami Tolpa (Gladstone Institutes) for helping with figure illustrations, Vinh Nguyen (UCSF Flow Core) for assistance with flow cytometry, Joseph J. Muldoon for guidance on design of *in vivo* studies, Marcus Alvarez for aid with visualization of the genome wide CRISPR screen, Carl Ward for assistance with BE screen analysis and interpretation, and Judy Lieberman for scientific discussion. We also thank the NCI BRB Preclinical Biologics Repository for provision of human cytokines.

This work was supported by NIH/NCI R01 CA288237 and the Parker Institute for Cancer Immunotherapy (A. Ma); NIH/NCI 1K08 CA252605-01, a Burroughs Wellcome Fund Career Award for Medical Scientists, the Lydia Preisler Shorenstein Donor Advised Fund, the Pascarella Scholars Fund, CRISPR Cures for Cancer, and the Parker Institute for Cancer Immunotherapy (J.C.); Grand Multiple Myeloma Translational Initiative, CRISPR Cures for Cancer and the Parker Institute for Cancer Immunotherapy (J.E.); NIH/NIDDK T32 DK007007 and by the Cancer Research Institute Irvington Fellowship / Samuel and Ruth Engelberg Postdoctoral Fellowship (CRI4667) (A.B.); South-Eastern Norway Regional Health Authority (2018046) (I.Ø); a Deutsche Forschungsgemeinschaft (DFG) fellowship (D.S.); the Susan G. Komen Breast Cancer Foundation (SAB190003) (A.A); NIAID R01AI171184, NIDCR U01DE028891, UC2DE032254, NIAID P01AI172523, NHGRI UM1HG012076, NIAMS P30AR070155, the Department of Defense, the Arc Research Institute, Parker Institute for Cancer Immunotherapy, and the Chan Zuckerberg Initiative (C.J.Y.); NCI R01CA276368, the Parker Institute for Cancer Immunotherapy, the Lloyd J. Old STAR award from the Cancer Research Institute (CRI), the Simons Foundation, and the CRISPR Cures for Cancer Initiative (A. Marson).

## Author contributions

A.B., S.B., L.R.S., B.A.M., J.E., J.C. and A. Ma conceived and designed the studies. A.B., S.B., L.R.S., C.C., and M.P. performed and analyzed *in vivo* experiments, while Z.L., Q.L., J.S. and C.H.W. provided assistance. S.B., N.K. and J.C. performed genome-wide CRISPR screen, and M.D. analyzed the next-generation sequencing screen data. A.B., M.P., R.A., D.S., and I.Ø performed and analyzed mouse tumor and T cell experiments, while N.L., E.Y., P.A. and C.J.B. provided assistance and guidance. S.B., L.R.S., C.C. and C.H. performed and analyzed human T cell experiments. C.J.Y. and A. Marson provided guidance for genomic ESM1b and base-editing analyses. The manuscript was written by A.B., S.B., L.R.S., B.A.M., J.E., J.C. and A. Ma and edited by C.J.B., A.A., C.J.Y. and A. Marson with input from all authors.

## Declaration of generative AI and AI-assisted technologies

The protein language model ESM1b (Meta AI) was utilized to predict the phenotypic impact of missense variants throughout the open reading frame of T*NFAIP3*/A20.

## Methods

### Mice

Male and female mice between 6–12 weeks of age were used for all experiments, following a protocol approved by the UCSF Institutional Animal Care and Use Committee. All mice were housed with a 12 h/12 h light/dark cycle, with food and water available *ad libitum*. A20^fl/fl^, A20^OTU^, A20^ZF4^ and A20^ZF7^ mice have been described^11,12,41^. The following mouse strains were obtained from The Jackson Laboratory: C57BL/6J (Strain #000664), BALB/cJ (#000651), NOD-*scid* IL2Rγ^null^ (NSG) (#005557), CD45.1 (#002014), *Cd8a*^−/−^ (#002665), E8I^cre^ (#008766), *Ifng*^−/−^ (#002287), *Ighm*^−/−^ (#002288), OT-I (#003831), *Prf1*^−/−^ (#002407), *Tcrd*^−/−^ (#002120), and *Tnf*^−/−^ (#005540). For CT26 experiments, A20^ZF7^ KI mice on a C57BL/6J background were backcrossed to BALB/cJ mice for seven generations.

### Human cell lines

A375 (ATCC), A375 mKate^+^,^33^ A375-CD19 and A375-CD19-mKate (generated by Alexander Marson’s lab at Gladstone Institutes/UCSF)^31^ cells were cultured in complete RPMI consisting of RPMI 1640 (Thermo Fisher) supplemented with 10% FBS (Corning), 1% glutamine (Life Technologies), 100 U/ml penicillin (Thermo Fisher) and 0.1 mg/ml streptomycin (Thermo Fisher). NALM6 (ATCC) expressing luciferase and eGFP (generated by Michel Sadelain’s lab at MSKCC)^91^ were cultured in complete RPMI supplemented with 1x MEM NEAA (Thermo Fisher), 1mM sodium pyruvate (Thermo Fisher), 55 µM 2-Mercaptoethanol (Thermo Fisher), and 1 mM HEPES (Thermo Fisher). HEK293T (ATCC) and Lenti X-HEK293T cells (TakaraLenti-XTM 293T) were cultured in DMEM (Thermo Fisher) supplemented with 10% fetal bovine serum (FBS; Corning), 1% glutamine (Life Technologies), 100 U/ml penicillin (Thermo Fisher) and 0.1 mg/ml streptomycin (Thermo Fisher). Flow cytometry was routinely performed on NALM6 and A375-CD19 cells to confirm CD19 antigen expression. NALM6, A375, A375-CD19 and 293T were routinely tested with the MycoAlert Detection kit (Lonza) and were found to be mycoplasma-free.

### Murine cell lines

MC38 (Kerafast), MC38-OVA (see below), B16F10 (ATCC) and B16-OVA (Sigma-Aldrich) tumor cells were cultured in complete DMEM (cDMEM; Corning), defined as DMEM supplemented with 10% fetal bovine serum (FBS; R&D Systems), 100 U/ml penicillin (MilliporeSigma), 0.1 mg/ml streptomycin (MilliporeSigma), 2 mM L-glutamine (Thermo Fisher), 10 mM HEPES buffer (CellGro), 1x MEM nonessential amino acids (Corning), and 1 mM sodium pyruvate (CellGro). CT26 (ATCC) tumor cells were cultured in complete RPMI 1640 (cRPMI; Gibco) with the aforementioned supplementation. MC38-OVA cells were generated by transfecting MC38 with pCI-neo-mOVA (Addgene #25099)^92^ by using Lipofectamine LTX Reagent with PLUS (Thermo Fisher Scientific), then selecting cells in 400 μg/ml G-418 (Research Products International). Murine tumor cell lines were routinely tested with the Lookout One-step Mycoplasma PCR Detection Kit (Sigma-Aldrich) and found to be mycoplasma-free.

### Primary human T cell isolation and culture

Primary human T cells were isolated from human PBMCs via negative selection (EasySep; STEMCELL). Leukopaks from deidentified healthy donors (STEMCELL) were used and came with consent from donors as collected by STEMCELL. Following isolation, T cells were activated using CTS Dynabeads Human T-Cell Activator CD3/CD28 (Thermo Fisher) at a cell:bead ratio of 1:1. Cells were cultured at 1 × 10^6^ cells/ml in complete human T cell growth medium defined as X-Vivo-15 media (Lonza) supplemented with 5% human serum (Gemini), 50 µM 2-mercaptoethanol (Thermo Fisher), and 10 mM N-acetyl-L-cysteine (MilliporeSigma). Cells were split every 48-72 hours. Human IL-7 and IL-15 (5 ng/ml each; Miltenyi) was added, unless stated otherwise.

### Primary murine T cell isolation and culture

Naive mouse polyclonal CD8^+^ T cells or OT-I cells were isolated by harvesting spleens from A20^ZF7^ KI or OT-I mice and mechanically disaggregating through a 70 μm cell strainer (Corning). Splenocytes were lysed in ACK buffer (Quality Biological) for 30 seconds, quenched in PBS (Gibco), filtered again, and counted. Naive CD8^+^ T cells were then isolated from splenocytes via negative selection (EasySep; STEMCELL).

### Preclinical tumor models in immunodeficient mice

Male NSG mice aged 8-12 weeks were used for human TCR-T and CAR-T cell experiments and randomized by tumor burden before T cell injections. For the A375 melanoma study, mice received 1 × 10^6^ A375 cells s.c. in the flank. Seven days later, mice were injected i.v. with 0.75 × 10^6^ 1G4 TCR-T cells. Tumor size was measured every 3-4 days and tumor volume was calculated using the formula v = 1/6 * π * length * width * (length + width) / 2.. For the NALM6 leukemia study, mice were injected i.v. with 0.5 × 10^6^ effLuc-eGFP NALM6 cells. Four days later, mice were injected i.v. with 0.1 × 10^6^ CD19 CAR-T cells. Tumor growth was monitored via BLI using a Xenogen IVIS Spectrum Imaging System (PerkinElmer). Mice were injected i.p. with 3 mg of D-luciferin (Gold Biotechnology) and were imaged 10 min later. For NALM6 iterative rechallenge, mice were injected i.v. with 3.0 × 10^6^ NALM6 cells on days 44, 51, 58, and 65 following CAR-T cell injection. Weight and health of mice were routinely monitored.

### Heterotopic tumor models in immunocompetent mice

MC38 (5 × 10^5^), MC38-OVA (5 × 10^5^), B16F10 (1 × 10^5^), B16-OVA (1 × 10^5^) or CT26 (5 × 10^5^) tumor cells were injected s.c. in 100 μL PBS. For B16F10 and B16-OVA, tumor cells were mixed 1:1 in PBS:Matrigel (Corning). Tumor size was measured every 3 days as length x width^2^ / 2. Experimental endpoint was achieved when length or width reached greater than 2 cm, or when tumors demonstrated significant ulceration. For A20^ZF7/ZF7^ regressor rechallenge, 60-80 days after initial MC38 inoculation, MC38 (2 × 10^6^) cells were injected s.c. in the opposite flank from the primary challenge.

### CD4^+^ and CD8^+^ T cell antibody-mediated depletion

For depletion of CD4^+^ T cells, 500 μg anti-CD4 (clone GK1.5; BioXCell) or isotype control (rat IgG2b, clone LTF-2; BioXCell) was injected intraperitoneally (i.p.) two days prior to tumor inoculation. On day 1 after tumor inoculation, and every 3 days thereafter until experimental endpoint, mice received 250 μg anti-CD4 or isotype control i.p. CD4^+^ T cell depletion in blood was confirmed by flow cytometry 14 days after tumor inoculation.

For depletion of CD8^+^ T cells, 500 μg anti-CD8α (clone 2.43; BioXCell) or isotype control (rat IgG2b, clone LTF-2; BioXCell) was injected intraperitoneally (i.p.) two days prior to tumor inoculation. On day 5 after tumor inoculation, and every 7 days thereafter until experimental endpoint, mice received 250 μg anti-CD8α or isotype control i.p. CD4^+^ T cell depletion in blood was confirmed by flow cytometry 14 days after tumor inoculation.

### CD8^+^ T cell reconstitution of Cd8a^−/−^ mice

*Cd8a^−/−^* (CD45.1/1) recipient mice underwent sublethal X-ray irradiation (450 cGy; X-Rad320; Precision X-Ray Irradiation). Naive polyclonal CD8^+^ T cells were isolated from donor mice as above. Three hours after irradiation, 3-8 × 10^6^ naive CD8^+^ T cells were injected i.v. into recipients. After 12 days, and two days prior to tumor inoculation, engraftment was confirmed by flow cytometry on blood cells. In one experiment, recipient mice were euthanized 14 days after tumor inoculation, and tumors were analyzed by flow cytometry to confirm CD8^+^ TIL reconstitution.

### OT-I cell adoptive transfer

For adoptive transfer of naive OT-I cells, isolation of OT-I cells was performed as above, and 1 × 10^6^ cells were injected i.v. into recipient mice one day prior to MC38-OVA inoculation. For adoptive transfer of pre-activated OT-I cells, total splenocytes harvested from OT-I mice were cultured at 2 × 10^6^ cells/ml in cRPMI containing 1 μg/ml OVA^257–264^ peptide (Invivogen) for 48 h. Cells were then counted, washed and replated at 2 × 10^5^ cells/ml in fresh cRPMI containing 50 μM β-mercaptoethanol (MilliporeSigma) and 50 IU/ml human IL-2 (NCI BRB Preclinical Biologics Repository), and expanded for an additional 48 h. Finally, 2.5-10 × 10^6^ pre-activated OT-I cells were injected i.v. into MC38-OVA-bearing mice on day 7 after tumor inoculation. For adoptive co-transfer experiments, recipient MC38-OVA-bearing WT mice (CD45.1/2) received equal numbers (2.5 × 10^6^ each) of A20^+/+^ (CD45.1/1) and A20^ZF7/ZF7^ (CD45.2/2) pre-activated OT-I cells on day 7 after tumor inoculation, and tumors were analyzed by FSFC on day 14 after tumor inoculation.

### Lentiviral production and transduction

Lenti-X 293T cells (Takara Bio) were seeded at 23 × 10^6^ cells per 225 cm² dish coated with poly-L-lysine (MilliporeSigma) in Opti-MEM (Thermo Fisher) supplemented with 5% FBS, 1X MEM Non-Essential Amino Acids (Thermo Fisher), and 1mM Sodium Pyruvate. The next day, 293T cells were transfected with transfer plasmids along with second-generation lentiviral packaging plasmids, pMD2.G (Addgene#12259) and psPAX2 (Addgene#12260), using the Lipofectamine 3000 reagent (Thermo Fisher), following the manufacturer’s protocol. Six hours after transfection, the medium was replaced with fresh media plus ViralBoost Reagent (500X) (Alstem). Viral supernatants collected after 24 and 48 hours were centrifuged at 500x*g* at 4°C for 10 min to remove debris and concentrated using Lenti-X concentrator (Takara Bio), then stored at 4°C overnight. The virus was further concentrated by centrifugation at 1500x*g* for 45 min at 4°C and resuspended in Opti-MEM at 100X the original volume, and then stored at −80°C. For T cell transduction with 1G4 TCR^93^ (Supp. Table 8), concentrated lentivirus was added 24 hours post-TCR activation to T cells at a 1:25 volume-to-volume ratio cultured in T cell media and gently mixed by tilting.

### Adeno-associated virus (AAV) production and transduction

The AAV cargo plasmid containing an HDR template for a TRAC CAR was co-transfected with packaging plasmids pHelper and pAAV Rep-Cap into HEK293T cells using polyethylenimine (PEI) to produce AAV6 particles,^40^ which were purified using iodixanol (STEMCELL) gradient ultracentrifugation. Viral titers were quantified through qPCR analysis performed on AAV samples treated with DNase I (NEB) and digested with proteinase K (Qiagen). HDR templates targeting TRAC were quantified using primers specific to the left homology arm of the HDR template specified in Supp. Table 7. qPCR was conducted using the SsoFast EvaGreen Supermix (Bio-Rad) on a StepOnePlus Real-Time PCR System (Applied Biosystems).

### CBE mRNA production

An *in vitro* transcription (IVT) plasmid was constructed that contained evoCDA1-BE4max along with a mutated T7 promoter. IVT templates were generated through PCR amplification of evoCDA1-BE4max, utilizing a forward primer to correct the T7 mutation and a reverse primer to add a poly-A tail (primers specified in Supp. Table 7). The resulting PCR product included the WT T7 promoter, a 5′ untranslated region with a Kozak sequence, the codon-optimized evoCDA1-BE4max coding sequence, a 3′ untranslated region, and a 145-bp poly-A tail. After purification, the PCR product was stored at −20°C until use. IVT reactions were conducted using the HiScribe T7 High Yield RNA Synthesis Kit (New England Biolabs), substituting UTP with N1-Methylpseudouridine-5’-Triphosphate (Trilink Biotechnologies) and the addition of 4 mM CleanCap (TriLink Biotechnologies). The transcribed mRNA was purified using lithium chloride and eluted in RNA storage solution (Thermo Fisher Scientific). The mRNA product was analyzed using an Agilent 4200 Tapestation system and subsequently stored at −80°C.

### CRISPR editing of primary human T Cells

Lyophilized sgRNA (Synthego) was rehydrated in TE buffer (Synthego) to a concentration of 80 μM for Cas9 editors and 120 μM for Cas12a editors. 48 hours after activation, T cells were de-beaded using EasySep magnets (STEMCELL) and electroporation of Cas9–sgRNA–RNP was conducted using the Amaxa P3 Primary Cell 96-well 4D-Nucleofector Kit (Lonza). For Cas9-edited T cells, sgRNAs were used to target AAVS1 and *TNFAIP3* (where guides were designed to either disrupt expression of the entire gene (A20^KO^), or to cause a truncation that eliminates the A20 ZF7 domain (the last exon) (A20^ZF7^) (sgRNA sequences specified in Supp. Table 6). For targeted insertion of the CAR into the *TRAC* locus, sgRNA for Cas9 or crRNA sequences for Cas12a were used as specified in Supp. Table 6. These sgRNAs were mixed with Cas9 or Cas12a protein (stock concentration 40 μM, Q3 MacroLab/UC Berkeley) at a molar ratio of 2:1 (sgRNA:Cas) and incubated at a temperature of 37°C for 15 min. 3 μl of the resulting RNP was mixed with 2 × 10^6^ T cells that had been resuspended in 20 μL of P3 and transferred into a 96-well electroporation plate using the pulse code EH115. T cells were rapidly recovered by addition of 100 μL of T cell media and incubated at 37°C for 15 min. Cells were then resuspended in T cell media at a concentration of 2 × 10^6^ live T cells per ml with the addition of IL-7 and IL-15 and transferred to appropriate culture vessels.

### TRAC targeting of CD19 CAR construct

For AAV-mediated CRISPR knock-in of CD19 CAR (Supp. Table 8) into *TRAC* locus, cells were edited as described above. The *TRAC* locus was targeted utilizing sgRNAs specified in Supp. Table 6. The recombinant AAV6 donor vector was added 30 min after electroporation at a multiplicity of infection of 5 × 10^4^, with an overnight incubation in serum-free T cell growth medium. The following day, edited cells were resuspended in complete T cell growth medium, expanded first at a density of 2 × 10^6^ cells/ml, and maintained at 1.5 × 10^6^ cells/ml. Flow cytometry was used to evaluate the knock-in efficiency through CAR staining with an anti-G4S antibody (Cell Signaling Technologies).

### Base-editing

48 hours after activation of human T cells with anti-CD3/CD28 Dynabeads (ThermoFisher), cells were de-beaded and resuspended in P3 buffer plus supplements (Lonza) at 2 × 10^6^ cells per 20 μL. Next, 2 µg of CBE mRNA along with 60 nmols of synthetic modified sgRNA (Synthego) targeting either B2Mi, A20^KO^ or A20^ZF7^ (sgRNAs specified in Supp. Table 6) were added to the resuspended cells. Cells were then electroporated and cultured as described in the Cas9-RNP electroporation section.

### Genome-wide CRISPR/Cas9 screening

T cells were isolated and stimulated as previously outlined, and 24 hours later were transduced with a lentiviral pool containing the genome-wide Brunello sgRNA library (Addgene #73178)^34^ at a volume to volume (v/v) ratio of 1:500 to achieve a targeted transduction efficiency of 50%. The human Brunello CRISPR knockout pooled library was a gift from David Root and John Doench. Following this transduction, the T cells were washed with PBS and electroporated with Cas9-RNP containing a non-targeting sgRNA (sequence specified in Supp. Table 6), and subsequently cultured as described. On day 12, T cells were stained with Dextramer-HLAA*0201/SLLMWITQV-APC (Immudex) and the percentage of TCR positive cells was determined by flow cytometry. These modified T cells were then co-cultured with A375 melanoma cells at an effector to target (E:T) ratio of 1:1 in complete T cell growth medium with the addition of 100 IU/ml IL-2 (R&D Systems). After 9 rounds of exposure to fresh tumor cells every 48 to 72 hours (PMID: 36002574), cells were collected and genomic DNA was isolated using the NucleoSpin Blood XL kit (Macherey-Nagel). Genomic DNA concentration was quantified using Invitrogen 1X dsDNA High Sensitivity assay kit (Thermo Fisher) on a Qubit Fluorometer (Invitrogen). PCR of the amplicon containing the guide sequences was carried out using Ex Taq DNA Polymerase (Takara Bio) and P5/P7 primers (IDT) (sequences specified in Supp. Table 7). The resulting amplicons were purified using SPRIselect Beads (Beckman-Coulter) and QC was conducted using a D1000 ScreenTape assay on a TapeStation (Agilent) prior to next-generation sequencing. Samples were pooled and sequenced on a NovaSeq X system at The Center for Advanced Technology (CAT) at UCSF.

MAGeCK52 v0.5.9.5 was used to quantify guide counts in each sample and tested for guide enrichment (PMID 25476604). Paired robust rank aggregation (RRA) analysis of guide count data from two donors was performed using the default parameters. Samples of T cells collected before co-culture with tumor cells (Time-Zero [T0]) were compared with samples collected after 4 (Time-Intermediate [TI], 1 donor), and 9 rounds of co-culture (Time-Final [TF]) as well as an age-matched arm not exposed to tumor cells [TF_ctrl_]. Guides with a read count of ≤ 40 in the control samples were filtered out. Gene-level RRA results and guide counts are included in Supp. Tables 1-5.

### Human T cell repetitive stimulation assay

For repetitive stimulation of human TCR-T or CAR-T cells, the day before setting up the assay, A375 or A375-CD19 cells were seeded in cRPMI. The following day, the medium was replaced with complete T cell growth medium, and antigen-specific T cells were added on top of the tumor cells at a 1:1 effector-to-target (E:T) ratio, with IL-2 added at a concentration of 100 IU/ml. Subsequent co-cultures were conducted every 48 to 72 hours. For each co-culture, T cells were collected, counted with the Cellaca MX High-throughput Cell Counter (Revvity), and then replated onto fresh tumor cells at the same 1:1 E:T ratio.

### Generation of NF-κB reporter AAV plasmid

NF-κB-eCFP reporter sequence (Addgene 118094)^94^ containing NF-κB response element, minimal promoter, and eCFP coding sequence was cloned into AAV backbone containing homology arms for B2M intron flanking polypyrimidine tracts, branchpoint sequence, splice acceptors and P2A sequences as previously defined for homology-directed repair^95^. The cassette was inserted in a reverse orientation with a terminating polyA tail SV40 sequence to allow for transcription of eCFP reporter without interference of *B2M* mRNA transcription.

### CRISPR editing of NF-κB-eCFP reporter CAR-T and TCR-T cells

Lyophilized sgRNA (Synthego) was rehydrated in TE buffer (Synthego) to a concentration of 120 μM for Cas9 base editors and 120 μM for Cas12a editors. 48 h after activation, T cells were de-beaded using EasySep magnets (STEMCELL), and electroporation of Cas12a–crRNA–RNP was conducted using the Amaxa P3 Primary Cell 96-well 4D-Nucleofector Kit (Lonza). For targeted insertion of the CAR into the *TRAC* locus and NF-κB reporter cassette into the first B2M intron (B2Mi), crRNA sequences for Cas12a were used as specified in Supp. Table 6. This crRNA was mixed with AsCas12a Ultra protein (stock concentration 40 μM, Q3 MacroLab/UC Berkeley) at a molar ratio of 2:1 (sgRNA:Cas) individually and incubated at a temperature of 37°C for 15 min. Cas9 RNPs with sgRNAs targeting AAVS1, A20^KO^ or A20^ZF7^, or CBE mRNA with sgRNAs targeting A20^KO^ or A20^ZF7^, were included as detailed above (Supp. Table 6). 2.5μl of each RNP mix, 0.5μL of sgRNA, and 0.5 μl of mRNA was mixed with 2 × 10^6^ T cells that had been resuspended in 18μL of P3 and transferred into a 96-well electroporation plate using the pulse code EH115. T cells were rapidly recovered by addition of 100 μl of serum-free T cell media containing 1 μM M3814 (ChemieTek), a non-homologous end joining inhibitor. Recovered cells were transferred into a GREX (Wilson Wolf, 80192M), combining a total of 4 × 10^6^ electroporated cells in a single well, an additional 400 μl of serum-free media and 1 μM M3814 was added, and cells were incubated at 37°C for 30 min. Recombinant AAV6 donor vectors were then added at a multiplicity of infection of 5 × 10^4^ with an overnight incubation. The following day, edited cells were diluted by adding 7.5 ml of complete T cell growth medium and allowed to expand for 5 days. Additional cytokines (human IL-7 and IL-15, 40 ng each) were added to culture every other day to supplement growth.

### Flow cytometric NF-κB reporter assay

A375 (for TCR-T experiments) or A375-CD19 (for CAR-T experiments) cells were seeded in cRPMI. The following day, the medium was replaced with complete T cell growth medium, and antigen-specific TCR-T or CAR-T cells were added on top of the tumor cells at a 1:2 effector-to-target (E:T) ratio, with IL-2 added at a concentration of 100 IU/ml. Cells were then collected 12 hours after stimulation by forcefully pipetting surface of flask. TCR-T or CAR-T cells were then stained with the appropriate fluorophore-conjugated surface antibody mix and incubated for 20 min at room temperature (RT). After staining, the cells were washed, resuspended in 120 μl of FACS buffer, and analyzed on a Cytek Aurora full spectrum flow cytometer.

### Murine T cell repetitive stimulation assay

Performed as previously described^70^. Briefly, splenocytes were harvested from WT mice, and dendritic cells (DCs) were isolated via EasySep Mouse Pan-DC Enrichment Kit (STEMCELL). DCs were resuspended in cRPMI at 1 × 10^6^ cells/ml and pulsed with 1 μg/ml OVA^257–264^ peptide for 30 min at 37°C. Next, naive OT-I cells isolated as above were added at a 1:1 ratio. After 2, 4 and 6 days, remaining OT-I cells were replated at 1 × 10^5^ cells/ml in cRPMI containing 50 μM β-mercaptoethanol, 50 IU/ml human IL-2 and fresh 1 μg/ml OVA^257–264^ peptide. On day 8, remaining OT-I T cells were collected for FSFC.

### Cytokine withdrawal and antigen-dependence

CAR-T cells were cultured in complete T cell growth medium without the addition of cytokines and either stimulated once on day 0 with CD19^+^ A375 melanoma cells at an effector-to-target (E:T) ratio of 1:1 or left unstimulated. For the stimulated arm, proliferation of CAR^+^ cells was monitored over a 25-day period by automated cell counting with a Cellaca MX High-throughput Cell Counter (Revvity) using AO/PI staining solution (Revvity). CAR^+^ cell percentages were evaluated alongside every count by flow cytometry using an anti-G4S antibody (Cell Signaling Technologies). The growth rates of CAR^+^ cells were calculated, and the fold change relative to the initial count was determined. In the unstimulated arm, total T cell counts were used to assess proliferation and viability using AO/PI over a 15-day period and the fold change was calculated based on the initial count.

### Sanger sequencing

Frozen cell pellets were resuspended in QuickExtract DNA Extraction Solution (Biosearch Technologies) at a concentration of 100 × 10^5^ cells per 50 µL of solution. gDNA was extracted following instructions according to manufacturer’s protocol. 5 µL of resulting solution was used as input template for a PCR reaction using Phusion High-Fidelity DNA Polymerase (NEB) to survey Exon 2, 5, 9 at the A20/*TNFAIP3* locus, and AAVS1 using the primers specified in Supp. Table 7. Resulting amplicons were purified using SPRI selection beads (Beckman Coulter) at 2X concentration following kit instructions. Purified amplicons were submitted to Quintara Biosciences for sanger sequencing using the forward and reverse primer used for the PCR amplification. Indel % and knock-out (KO) score (proportion of indels that are at least 21 bp long or indicate a frameshift) were analyzed with the Synthego ICE analysis tool.

### Next-generation sequencing of base-edited T cells

Cell pellets and gDNA extraction were prepared as above. 5uL of extraction solution was used as input for the PCR reaction to survey *TNFAIP3* Exon 2, *TNFAIP3* Exon 9 (A20^ZF7^ domain), and B2Mi loci using the primer pairs specified in Supp. Table 7. The PCR reaction was comprised of Q5 High-Fidelity 2X Master Mix (New England Biolabs), loading 5 µL of the gDNA extraction in a reaction volume of 50 µL, and following recommended cycling conditions from manufacturer for 32 cycles. The resulting amplicons were purified using SPRI selection beads (Beckman Coulter) at 2X concentration following the kit instructions. Amplicons were analyzed with Agilent High Sensitivity D5000 ScreenTape (Agilent) on a TapeStation (Agilent) to quantify expected product sizes. Purified amplicons were submitted to Quintara Biosciences for next-generation sequencing and analyzed using custom python pipeline to calculate total mutant basepair abundance at targeted loci.

### Flow cytometry of engineered human T cells in vitro

For *in vitro* analysis of cell surface activation markers, TCR-T and CAR-T cells (2 × 10^5^ to 5 × 10^5^) were seeded into round-bottom 96-well plates (Corning). TCR-T cells were then stimulated with anti-CD3/CD28 Dynabeads, and CD19 CAR-T cells were stimulated with CD19^+^ A375 melanoma cells at a bead/cell to cell ratio of 1 to 1 at 37°C for 6 hours. Cells were then centrifuged at 300g for 5 min, washed once with 200 μL of FACS buffer, and stained with the appropriate antibodies (1 μL per 100 μL staining buffer) for 20 min at 4°C in the dark. Cells were then washed twice with FACS buffer. For exhaustion and differentiation markers, cells were stained without prior stimulation following the same approach. Samples were analyzed using the Attune NXT Cytometer (Invitrogen) or LSRFortessa X-50 (BD) and data were processed and analyzed using FlowJo 10.9.1 (FlowJo, Inc.). For compensation controls, UltraComp eBeads compensation beads (ThermoFisher) were used. Antibodies used for staining are specified in Supp. Table 9.

### Proliferation assay

TCR-T or CAR-T cells were resuspended in PBS (Gibco) and stained with the fluorescent dye CellTrace Violet (Thermo Fisher Scientific) at a working concentration of 5 µM, following the manufacturer’s protocol. After staining, TCR-T cells were resuspended in 200 µL complete T cell growth medium and stimulated with 3.125 µL/mL of ImmunoCult Human CD3/CD28 T Cell Activator (STEMCELL) for 72 hours. For stimulation of CAR-T cells, CD19^+^ A375 melanoma cells were used. Flow cytometry was used to measure cell division percentages across conditions and analyzed with the proliferation modeling tool in FlowJo 10.9.1.

### Flow cytometry of bone marrow in NALM6-bearing mice

For analysis of bone marrow, NALM6-bearing mice were euthanized 14 days after CAR-T cell injection. For bone marrow extraction, the femur and tibia from each leg were dissected and crushed in MACS buffer consisting of PBS (Gibco), 2% FBS and 1mM EDTA using a mortar and pestle. The resulting cell suspension was filtered through a 70 μm cell strainer (Corning), centrifuged at 300x*g* for 5 min, and red blood cells were lysed using ACK lysing buffer (Quality Biological) for 2 min, with the reaction subsequently quenched using MACS buffer. Each sample was first resuspended in 500 μL of MACS buffer. 250 μL of each sample were used for analysis, centrifuged, and resuspended in 100 μL of FACS buffer. Cells were first incubated with 10 μL of mouse Fc block (Miltenyi Biotec) per sample and incubated for 20 min at room temperature. The cells were then stained with the appropriate antibody mix and incubated for 45 min at room temperature (RT). After staining, the cells were washed, resuspended in 150 μL of FACS buffer with 50 μL of counting beads per sample (Thermo Fisher), and analyzed on an Aurora spectral flow cytometer (Cytek). Data were processed and analyzed in FlowJo. SPICE plots were generated using SPICE 6.1. Antibodies used for staining are specified in Supp. Table 9.

### Multiplex immunoassay (LEGENDplex)

Before and after several rounds of co-culture with tumor cells, 2 × 10^5^ TCR T or CAR-T cells were resuspended in 200 μL fresh complete T cell growth medium without cytokines. TCR-T cells were seeded on top of A375 melanoma cells at an effector to target ratio of 1 to 1 or stimulated with CTS Dynabeads Human T-Cell Activator CD3/CD28 at a cell to bead ratio of 1 to 1. CAR-T were seeded on top of CD19^+^ A375 melanoma cells at an effector to target ratio of 1 to 1. 24 hours later, the supernatant from the co-cultures was harvested and spun for 5 min at 300g at 4 °C to remove cell debris. The collected supernatant was then stored at −80 °C and subsequently analyzed using the LEGENDplex Human CD8/NK Panel (13-plex) (Biolegend) as per the manufacturer’s guidelines. Results were analyzed using the LEGENDplex analysis tool (Biolegend). When necessary, the supernatant was diluted, and the resulting concentration was multiplied by the dilution factor to obtain the absolute concentration in pg/ml.

### FSFC of murine TILs

Harvested tumors were digested for 30 min at 37°C in RPMI containing 0.13 Wünsch units/ml Liberase TM (Roche) and 72 μg/ml DNase I (Roche), with intermittent agitation. Tumors were then resuspended in FSFC buffer (PBS + 2% FBS + 5 mM EDTA) and incubated for 15 min at 37°C, after which samples were filtered through 70 μm nylon cell strainer to achieve a single cell suspension. Samples were then stained for 15 min at room temperature (RT) in PBS containing Zombie NIR live/dead stain (BioLegend), then washed in FSFC buffer. Samples were then treated with 10 μg/ml anti-CD16/CD32 (BioXCell) in FSFC buffer containing a 1:10 dilution of Brilliant Stain Buffer (Thermo Fisher) for 10 min at RT, after which surface antibody staining cocktail was added and samples were incubated for an additional 20 min at 37°C. Samples were washed in FSFC buffer, then fixed and permeabilized for 40 min at RT using eBioscience Foxp3 Transcription Factor Staining Kit (Thermo Fisher). Samples were then blocked for 10 min at RT in Foxp3 kit buffer containing 1:10 dilution of Brilliant Buffer and 2% rat serum (Jackson ImmunoResearch), after which intracellular antibody staining cocktail was added and samples were incubated an additional 30 min at RT. Samples were washed, resuspended in FSFC buffer, acquired on a Cytek Aurora, and analyzed with FlowJo. We set a minimum cutoff of 100 total cells within a given CD8^+^ TIL subset for the sample to be included in downstream analyses. Antibodies used for staining are specified in Supp. Table 9.

### Flow cytometry of murine blood leukocytes

Anesthetized mice underwent retro-orbital phlebotomy with heparanized capillary tubes (Thermo Fisher). Blood cells were lysed for 15 min in ACK buffer, quenched in excess PBS, filtered via 70 μm cell strainer, and pelleted before performing live/dead and surface stain.

### Intracellular cytokine staining of murine TILs

Tumor single cell suspensions were plated at 1 × 10^6^ cells/ml in cRPMI containing 50 ng/ml PMA (MilliporeSigma) and 500 ng/ml ionomycin (MilliporeSigma) and incubated at 37°C. After 1 hour, 5 μg/ml brefeldin A (BioLegend) was added, and cells were incubated an additional 4 h at 37°C. Cells were subsequently pelleted and submitted to live/dead and surface stain as above. Cells were then fixed, permeabilized, and stained for cytokines and perforin using the Cytofix/Cytoperm kit (BD Biosciences) per manufacturer’s protocol.

### CD107a staining

Tumor single cell suspensions (1 × 10^6^ cells/ml) were incubated for 1 h at 37°C in cRPMI containing 50 ng/ml PMA, 500 ng/ml ionomycin, 2 μM monensin (BioLegend), 5 μg/ml brefeldin A, 10 μg/ml anti-CD16/CD32, and 2 μg/ml PE anti-mouse CD107a. Cells were subsequently pelleted and submitted to live/dead and surface stain as above. CD8^+^ TIL subsets were identified as in Supp. Fig. 4A, with the surface marker Ly108 acting as a surrogate for TCF-1 to delineate Tpex from Ttex.

### In vivo brefeldin treatment

As described^76^, six hours prior to euthanization, day 12-15 MC38-bearing mice received i.p. injection of 250 μg brefeldin (MilliporeSigma) or 5% DMSO (vehicle control). Mice were closely monitored for six hours and showed no visible signs of distress.

### In vitro killing assay

Killing assays were performed using TCR-T and CAR-T cells, where antigen-specific T cells were co-cultured with pre-plated mKate^+^ A375 melanoma cells in a 96-well flat-bottom plate (Corning). T cells were seeded in various effector-to-target (E:T) ratios. For the TCR-T assays, mKate^+^ A375 melanoma cells (ATCC) were utilized, whereas for the CAR-T assays, CD19-expressing mKate^+^ A375 cells were used. Plates were imaged every 4-6 hours for at least 72 hours using the IncuCyte S3 live-cell imaging system (Essen Bioscience). The red object counts of mKate+ objects per well were recorded over time. Cancer cell growth was calculated at each time point for each replicate by normalization to the red object count at time zero.

### Immunoblotting

For activated conditions only, A20 modified and control T cells were stimulated for 24 hours with CTS Dynabeads Human T-Cell Activator CD3/CD28 (Thermo Fisher). Frozen cell pellets were resuspended in Pierce RIPA buffer (Thermo Fisher) plus complete protease inhibitor (Roche) and incubated on a rotator at 4 °C for 30 min and spun down at >16,000 g for 20 min at 4 °C. The cell lysates were stored at −80 °C if not used immediately. Protein concentrations were measured using the Pierce BCA Protein Assay (Thermo Fisher). 20 µg of protein from each sample were loaded onto 4–20% tris-glycine SDS gels (Bio-Rad) and transferred to a PVDF membrane (Bio-Rad) using the Bio-Rad Trans-Blot Transfer System. The membrane was blocked with AdvanBlock-Chemi solution (Advansta) for 1 hour at RT and incubated with primary antibodies overnight at 4 °C. The following primary antibodies were utilized: A20/*TNFAIP3* antibody (1:1000 dilution, Santa Cruz), β-actin rabbit monoclonal antibody (1:5000 dilution, Abclonal). For secondary antibodies, anti-rabbit IgG HRP-linked antibody (1:2000 dilution, Cell Signaling) and anti-mouse IgG HRP-linked antibody (1:2000 dilution, Cell Signaling) were used. Membranes were developed with SuperSignal West Femto Maximum Sensitivity Substrate (Thermo Scientific) and imaged on a ChemiDoc Touch Imaging System (Bio-Rad). Images were cropped using Adobe Photoshop but were not further altered. Antibodies used for staining are specified in Supp. Table 9.

### Quantifications and statistical analyses

Details for the statistical analyses for all experiments are provided in the figure legends. Plots were generated in GraphPad Prism. Paired t tests of *in vitro* data from 2-3 human donors do not account for correlation among the technical replicates and do not include corrections for multiple comparison. The survival outcomes for mice are depicted through Kaplan-Meier curves.

## References

1. Baessler, A., and Vignali, D.A.A. (2024). T Cell Exhaustion. Annu Rev Immunol 42, 179–206. 10.1146/annurev-immunol-090222-110914.

2. Im, S.J., Hashimoto, M., Gerner, M.Y., Lee, J., Kissick, H.T., Burger, M.C., Shan, Q., Hale, J.S., Nasti, T.H., Sharpe, A.H., et al. (2016). Defining CD8^+^ T cells that provide the proliferative burst after PD-1 therapy. Nature 537, 417–421. 10.1038/nature19330.

3. Siddiqui, I., Schaeuble, K., Chennupati, V., Fuertes Marraco, S.A., Calderon-Copete, S., Pais Ferreira, D., Carmona, S.J., Scarpellino, L., Gfeller, D., Pradervand, S., et al. (2019). Intratumoral Tcf1^+^PD-1^+^CD8^+^ T cells with stem-like properties promote tumor control in response to vaccination and checkpoint blockade immunotherapy. Immunity 50, 195–211.e110. 10.1016/j.immuni.2018.12.021.

4. Miller, B.C., Sen, D.R., Al Abosy, R., Bi, K., Virkud, Y.V., LaFleur, M.W., Yates, K.B., Lako, A., Felt, K., Naik, G.S., et al. (2019). Subsets of exhausted CD8^+^ T cells differentially mediate tumor control and respond to checkpoint blockade. Nat Immunol 20, 326–336. 10.1038/s41590-019-0312-6.

5. Wang, D., Prager, B.C., Gimple, R.C., Aguilar, B., Alizadeh, D., Tang, H., Lv, D., Starr, R., Brito, A., Wu, Q., et al. (2021). CRISPR Screening of CAR T Cells and Cancer Stem Cells Reveals Critical Dependencies for Cell-Based Therapies. Cancer Discov 11, 1192–1211. 10.1158/2159-8290.CD-20-1243.

6. Freitas, K.A., Belk, J.A., Sotillo, E., Quinn, P.J., Ramello, M.C., Malipatlolla, M., Daniel, B., Sandor, K., Klysz, D., Bjelajac, J., et al. (2022). Enhanced T cell effector activity by targeting the Mediator kinase module. Science 378, eabn5647. 10.1126/science.abn5647.

7. Schmidt, R., Ward, C.C., Dajani, R., Armour-Garb, Z., Ota, M., Allain, V., Hernandez, R., Layeghi, M., Xing, G., Goudy, L., et al. (2024). Base-editing mutagenesis maps alleles to tune human T cell functions. Nature 625, 805–812. 10.1038/s41586-023-06835-6.

8. Walsh, Z.H., Shah, P., Kothapalli, N., Shah, S.B., Nikolenyi, G., Brodtman, D.Z., Leuzzi, G., Rogava, M., Mu, M., Ho, P., et al. (2024). Mapping variant effects on anti-tumor hallmarks of primary human T cells with base-editing screens. Nat Biotechnol. 10.1038/s41587-024-02235-x.

9. Malynn, B.A., and Ma, A. (2019). A20: A multifunctional tool for regulating immunity and preventing disease. Cell Immunol 340, 103914. 10.1016/j.cellimm.2019.04.002.

10. Lee, E.G., Boone, D.L., Chai, S., Libby, S.L., Chien, M., Lodolce, J.P., and Ma, A. (2000). Failure to regulate TNF-induced NF-kappaB and cell death responses in A20-deficient mice. Science 289, 2350–2354. 10.1126/science.289.5488.2350.

11. Lu, T.T., Onizawa, M., Hammer, G.E., Turer, E.E., Yin, Q., Damko, E., Agelidis, A., Shifrin, N., Advincula, R., Barrera, J., et al. (2013). Dimerization and ubiquitin mediated recruitment of A20, a complex deubiquitinating enzyme. Immunity 38, 896–905. 10.1016/j.immuni.2013.03.008.

12. Razani, B., Whang, M.I., Kim, F.S., Nakamura, M.C., Sun, X., Advincula, R., Turnbaugh, J.A., Pendse, M., Tanbun, P., Achacoso, P., et al. (2020). Non-catalytic ubiquitin binding by A20 prevents psoriatic arthritis-like disease and inflammation. Nat Immunol 21, 422–433. 10.1038/s41590-020-0634-4.

13. Martens, A., Priem, D., Hoste, E., Vetters, J., Rennen, S., Catrysse, L., Voet, S., Deelen, L., Sze, M., Vikkula, H., et al. (2020). Two distinct ubiquitin-binding motifs in A20 mediate its anti-inflammatory and cell-protective activities. Nat Immunol 21, 381–387. 10.1038/s41590-020-0621-9.

14. Coornaert, B., Baens, M., Heyninck, K., Bekaert, T., Haegman, M., Staal, J., Sun, L., Chen, Z.J., Marynen, P., and Beyaert, R. (2008). T cell antigen receptor stimulation induces MALT1 paracaspase-mediated cleavage of the NF-kappaB inhibitor A20. Nat Immunol 9, 263–271. 10.1038/ni1561.

15. Düwel, M., Welteke, V., Oeckinghaus, A., Baens, M., Kloo, B., Ferch, U., Darnay, B.G., Ruland, J., Marynen, P., and Krappmann, D. (2009). A20 negatively regulates T cell receptor signaling to NF-kappaB by cleaving Malt1 ubiquitin chains. J Immunol 182, 7718–7728. 10.4049/jimmunol.0803313.

16. Onizawa, M., Oshima, S., Schulze-Topphoff, U., Oses-Prieto, J.A., Lu, T., Tavares, R., Prodhomme, T., Duong, B., Whang, M.I., Advincula, R., et al. (2015). The ubiquitin-modifying enzyme A20 restricts ubiquitination of the kinase RIPK3 and protects cells from necroptosis. Nat Immunol 16, 618–627. 10.1038/ni.3172.

17. Dabbah-Krancher, G., Ruchinskas, A., Kallarakal, M.A., Lee, K.P., Bauman, B.M., Epstein, B., Yin, H., Krappmann, D., Schaefer, B.C., and Snow, A.L. (2024). A20 intrinsically influences human effector T-cell survival and function by regulating both NF-κB and JNK signaling. Eur J Immunol 54, e2451245. 10.1002/eji.202451245.

18. Just, S., Nishanth, G., Buchbinder, J.H., Wang, X., Naumann, M., Lavrik, I., and Schlüter, D. (2016). A20 Curtails Primary but Augments Secondary CD8+ T Cell Responses in Intracellular Bacterial Infection. Sci Rep 6, 39796. 10.1038/srep39796.

19. LaFleur, M.W., Milling, L.E., Prathima, P., Li, V., Lemmen, A.M., Streeter, I.S.L., Heisig, P.K.S., Derosia, N.M., Riffo, E., Xu, H., et al. (2025). A STUB1-CHIC2 complex inhibits CD8+ T cells to restrain tumor immunity. Nat Immunol 26, 1476–1487. 10.1038/s41590-025-02231-6.

20. Giordano, M., Roncagalli, R., Bourdely, P., Chasson, L., Buferne, M., Yamasaki, S., Beyaert, R., van Loo, G., Auphan-Anezin, N., Schmitt-Verhulst, A.M., and Verdeil, G. (2014). The tumor necrosis factor alpha-induced protein 3 (TNFAIP3, A20) imposes a brake on antitumor activity of CD8 T cells. Proc Natl Acad Sci U S A 111, 11115–11120. 10.1073/pnas.1406259111.

21. Mukohara, F., Iwata, K., Ishino, T., Inozume, T., Nagasaki, J., Ueda, Y., Suzawa, K., Ueno, T., Ikeda, H., Kawase, K., et al. (2024). Somatic mutations in tumor-infiltrating lymphocytes impact on antitumor immunity. Proc Natl Acad Sci U S A 121, e2320189121. 10.1073/pnas.2320189121.

22. Luo, C., Zhang, R., Guo, R., Wu, L., Xue, T., He, Y., Jin, Y., Zhao, Y., Zhang, Z., Zhang, P., et al. (2025). Integrated computational analysis identifies therapeutic targets with dual action in cancer cells and T cells. Immunity. 10.1016/j.immuni.2025.02.007.

23. van den Broek, M.E., Kägi, D., Ossendorp, F., Toes, R., Vamvakas, S., Lutz, W.K., Melief, C.J., Zinkernagel, R.M., and Hengartner, H. (1996). Decreased tumor surveillance in perforin-deficient mice. J Exp Med 184, 1781–1790. 10.1084/jem.184.5.1781.

24. Prévost-Blondel, A., Neuenhahn, M., Rawiel, M., and Pircher, H. (2000). Differential requirement of perforin and IFN-gamma in CD8 T cell-mediated immune responses against B16.F10 melanoma cells expressing a viral antigen. Eur J Immunol 30, 2507–2515. 10.1002/1521-4141(200009)30:9<2507::AID-IMMU2507>3.0.CO;2-V.

25. Street, S.E., Cretney, E., and Smyth, M.J. (2001). Perforin and interferon-gamma activities independently control tumor initiation, growth, and metastasis. Blood 97, 192–197. 10.1182/blood.v97.1.192.

26. Poehlein, C.H., Hu, H.M., Yamada, J., Assmann, I., Alvord, W.G., Urba, W.J., and Fox, B.A. (2003). TNF plays an essential role in tumor regression after adoptive transfer of perforin/IFN-gamma double knockout effector T cells. J Immunol 170, 2004–2013. 10.4049/jimmunol.170.4.2004.

27. Cao, X., Cai, S.F., Fehniger, T.A., Song, J., Collins, L.I., Piwnica-Worms, D.R., and Ley, T.J. (2007). Granzyme B and perforin are important for regulatory T cell-mediated suppression of tumor clearance. Immunity 27, 635–646. 10.1016/j.immuni.2007.08.014.

28. Waldner, M.J., Wirtz, S., Becker, C., Seidel, D., Tubbe, I., Cappel, K., Hähnel, P.S., Galle, P.R., Schuler, M., and Neurath, M.F. (2010). Perforin deficiency attenuates inflammation and tumor growth in colitis-associated cancer. Inflamm Bowel Dis 16, 559–567. 10.1002/ibd.21107.

29. Juneja, V.R., McGuire, K.A., Manguso, R.T., LaFleur, M.W., Collins, N., Haining, W.N., Freeman, G.J., and Sharpe, A.H. (2017). PD-L1 on tumor cells is sufficient for immune evasion in immunogenic tumors and inhibits CD8 T cell cytotoxicity. J Exp Med 214, 895–904. 10.1084/jem.20160801.

30. Kearney, C.J., Vervoort, S.J., Hogg, S.J., Ramsbottom, K.M., Freeman, A.J., Lalaoui, N., Pijpers, L., Michie, J., Brown, K.K., Knight, D.A., et al. (2018). Tumor immune evasion arises through loss of TNF sensitivity. Sci Immunol 3. 10.1126/sciimmunol.aar3451.

31. Carnevale, J., Shifrut, E., Kale, N., Nyberg, W.A., Blaeschke, F., Chen, Y.Y., Li, Z., Bapat, S.P., Diolaiti, M.E., O’Leary, P., et al. (2022). RASA2 ablation in T cells boosts antigen sensitivity and long-term function. Nature. 10.1038/s41586-022-05126-w.

32. Blaeschke, F., Chen, Y.Y., Apathy, R., Daniel, B., Chen, A.Y., Chen, P.A., Sandor, K., Zhang, W., Li, Z., Mowery, C.T., et al. (2023). Modular pooled discovery of synthetic knockin sequences to program durable cell therapies. Cell 186, 4216–4234.e4233. 10.1016/j.cell.2023.08.013.

33. Shifrut, E., Carnevale, J., Tobin, V., Roth, T.L., Woo, J.M., Bui, C.T., Li, P.J., Diolaiti, M.E., Ashworth, A., and Marson, A. (2018). Genome-wide CRISPR Screens in Primary Human T Cells Reveal Key Regulators of Immune Function. Cell 175, 1958–1971.e1915. 10.1016/j.cell.2018.10.024.

34. Doench, J.G., Fusi, N., Sullender, M., Hegde, M., Vaimberg, E.W., Donovan, K.F., Smith, I., Tothova, Z., Wilen, C., Orchard, R., et al. (2016). Optimized sgRNA design to maximize activity and minimize off-target effects of CRISPR-Cas9. Nat Biotechnol 34, 184–191. 10.1038/nbt.3437.

35. Lv, J., Qin, L., Zhao, R., Wu, D., Wu, Z., Zheng, D., Li, S., Luo, M., Wu, Q., Long, Y., et al. (2023). Disruption of CISH promotes the antitumor activity of human T cells and decreases PD-1 expression levels. Mol Ther Oncolytics 28, 46–58. 10.1016/j.omto.2022.12.003.

36. Prinzing, B., Zebley, C.C., Petersen, C.T., Fan, Y., Anido, A.A., Yi, Z., Nguyen, P., Houke, H., Bell, M., Haydar, D., et al. (2021). Deleting DNMT3A in CAR T cells prevents exhaustion and enhances antitumor activity. Sci Transl Med 13, eabh0272. 10.1126/scitranslmed.abh0272.

37. Gurusamy, D., Henning, A.N., Yamamoto, T.N., Yu, Z., Zacharakis, N., Krishna, S., Kishton, R.J., Vodnala, S.K., Eidizadeh, A., Jia, L., et al. (2020). Multi-phenotype CRISPR-Cas9 Screen Identifies p38 Kinase as a Target for Adoptive Immunotherapies. Cancer Cell 37, 818–833.e819. 10.1016/j.ccell.2020.05.004.

38. Liao, X., Li, W., Zhou, H., Rajendran, B.K., Li, A., Ren, J., Luan, Y., Calderwood, D.A., Turk, B., Tang, W., et al. (2024). The CUL5 E3 ligase complex negatively regulates central signaling pathways in CD8^+^ T cells. Nat Commun 15, 603. 10.1038/s41467-024-44885-0.

39. Belk, J.A., Yao, W., Ly, N., Freitas, K.A., Chen, Y.T., Shi, Q., Valencia, A.M., Shifrut, E., Kale, N., Yost, K.E., et al. (2022). Genome-wide CRISPR screens of T cell exhaustion identify chromatin remodeling factors that limit T cell persistence. Cancer Cell 40, 768–786.e767. 10.1016/j.ccell.2022.06.001.

40. Eyquem, J., Mansilla-Soto, J., Giavridis, T., van der Stegen, S.J., Hamieh, M., Cunanan, K.M., Odak, A., Gönen, M., and Sadelain, M. (2017). Targeting a CAR to the TRAC locus with CRISPR/Cas9 enhances tumour rejection. Nature 543, 113–117. 10.1038/nature21405.

41. Tavares, R.M., Turer, E.E., Liu, C.L., Advincula, R., Scapini, P., Rhee, L., Barrera, J., Lowell, C.A., Utz, P.J., Malynn, B.A., and Ma, A. (2010). The ubiquitin modifying enzyme A20 restricts B cell survival and prevents autoimmunity. Immunity 33, 181–191. 10.1016/j.immuni.2010.07.017.

42. Aki, A., Nagasaki, M., Malynn, B.A., Ma, A., and Kagari, T. (2017). Hypomorphic A20 expression confers susceptibility to psoriasis. PLoS One 12, e0180481. 10.1371/journal.pone.0180481.

43. Hammer, G.E., Turer, E.E., Taylor, K.E., Fang, C.J., Advincula, R., Oshima, S., Barrera, J., Huang, E.J., Hou, B., Malynn, B.A., et al. (2011). Expression of A20 by dendritic cells preserves immune homeostasis and prevents colitis and spondyloarthritis. Nat Immunol 12, 1184–1193. 10.1038/ni.2135.

44. Vande Walle, L., Van Opdenbosch, N., Jacques, P., Fossoul, A., Verheugen, E., Vogel, P., Beyaert, R., Elewaut, D., Kanneganti, T.D., van Loo, G., and Lamkanfi, M. (2014). Negative regulation of the NLRP3 inflammasome by A20 protects against arthritis. Nature 512, 69–73. 10.1038/nature13322.

45. Duong, B.H., Onizawa, M., Oses-Prieto, J.A., Advincula, R., Burlingame, A., Malynn, B.A., and Ma, A. (2015). A20 restricts ubiquitination of pro-interleukin-1β protein complexes and suppresses NLRP3 inflammasome activity. Immunity 42, 55–67. 10.1016/j.immuni.2014.12.031.

46. Komander, D., and Barford, D. (2008). Structure of the A20 OTU domain and mechanistic insights into deubiquitination. Biochem J 409, 77–85. 10.1042/BJ20071399.

47. Boone, D.L., Turer, E.E., Lee, E.G., Ahmad, R.C., Wheeler, M.T., Tsui, C., Hurley, P., Chien, M., Chai, S., Hitotsumatsu, O., et al. (2004). The ubiquitin-modifying enzyme A20 is required for termination of Toll-like receptor responses. Nat Immunol 5, 1052–1060. 10.1038/ni1110.

48. Wertz, I.E., O’Rourke, K.M., Zhou, H., Eby, M., Aravind, L., Seshagiri, S., Wu, P., Wiesmann, C., Baker, R., Boone, D.L., et al. (2004). De-ubiquitination and ubiquitin ligase domains of A20 downregulate NF-kappaB signalling. Nature 430, 694–699. 10.1038/nature02794.

49. Bosanac, I., Wertz, I.E., Pan, B., Yu, C., Kusam, S., Lam, C., Phu, L., Phung, Q., Maurer, B., Arnott, D., et al. (2010). Ubiquitin binding to A20 ZnF4 is required for modulation of NF-κB signaling. Mol Cell 40, 548–557. 10.1016/j.molcel.2010.10.009.

50. Shembade, N., Ma, A., and Harhaj, E.W. (2010). Inhibition of NF-kappaB signaling by A20 through disruption of ubiquitin enzyme complexes. Science 327, 1135–1139. 10.1126/science.1182364.

51. Skaug, B., Chen, J., Du, F., He, J., Ma, A., and Chen, Z.J. (2011). Direct, noncatalytic mechanism of IKK inhibition by A20. Mol Cell 44, 559–571. 10.1016/j.molcel.2011.09.015.

52. Wertz, I.E., Newton, K., Seshasayee, D., Kusam, S., Lam, C., Zhang, J., Popovych, N., Helgason, E., Schoeffler, A., Jeet, S., et al. (2015). Phosphorylation and linear ubiquitin direct A20 inhibition of inflammation. Nature 528, 370–375. 10.1038/nature16165.

53. Rives, A., Meier, J., Sercu, T., Goyal, S., Lin, Z., Liu, J., Guo, D., Ott, M., Zitnick, C.L., Ma, J., and Fergus, R. (2021). Biological structure and function emerge from scaling unsupervised learning to 250 million protein sequences. Proc Natl Acad Sci U S A 118. 10.1073/pnas.2016239118.

54. Brandes, N., Goldman, G., Wang, C.H., Ye, C.J., and Ntranos, V. (2023). Genome-wide prediction of disease variant effects with a deep protein language model. Nat Genet 55, 1512–1522. 10.1038/s41588-023-01465-0.

55. Tokunaga, F., Nishimasu, H., Ishitani, R., Goto, E., Noguchi, T., Mio, K., Kamei, K., Ma, A., Iwai, K., and Nureki, O. (2012). Specific recognition of linear polyubiquitin by A20 zinc finger 7 is involved in NF-κB regulation. EMBO J 31, 3856–3870. 10.1038/emboj.2012.241.

56. Verhelst, K., Carpentier, I., Kreike, M., Meloni, L., Verstrepen, L., Kensche, T., Dikic, I., and Beyaert, R. (2012). A20 inhibits LUBAC-mediated NF-κB activation by binding linear polyubiquitin chains via its zinc finger 7. EMBO J 31, 3845–3855. 10.1038/emboj.2012.240.

57. Hoft, D.F., Lynch, R.G., and Kirchhoff, L.V. (1993). Kinetic analysis of antigen-specific immune responses in resistant and susceptible mice during infection with Trypanosoma cruzi. J Immunol 151, 7038–7047.

58. Labanieh, L., and Mackall, C.L. (2023). CAR immune cells: design principles, resistance and the next generation. Nature 614, 635–648. 10.1038/s41586-023-05707-3.

59. Jin, Y., An, X., Mao, B., Sun, R., Kumari, R., Chen, X., Shan, Y., Zang, M., Xu, L., Muntel, J., et al. (2022). Different syngeneic tumors show distinctive intrinsic tumor-immunity and mechanisms of actions (MOA) of anti-PD-1 treatment. Sci Rep 12, 3278. 10.1038/s41598-022-07153-z.

60. Lewis, K.E., Selby, M.J., Masters, G., Valle, J., Dito, G., Curtis, W.R., Garcia, R., Mink, K.A., Waggie, K.S., Holdren, M.S., et al. (2017). Interleukin-21 combined with PD-1 or CTLA-4 blockade enhances antitumor immunity in mouse tumor models. Oncoimmunology 7, e1377873. 10.1080/2162402X.2017.1377873.

61. Vignali, P.D.A., DePeaux, K., Watson, M.J., Ye, C., Ford, B.R., Lontos, K., McGaa, N.K., Scharping, N.E., Menk, A.V., Robson, S.C., et al. (2023). Hypoxia drives CD39-dependent suppressor function in exhausted T cells to limit antitumor immunity. Nat Immunol 24, 267–279. 10.1038/s41590-022-01379-9.

62. LaFleur, M.W., Nguyen, T.H., Coxe, M.A., Miller, B.C., Yates, K.B., Gillis, J.E., Sen, D.R., Gaudiano, E.F., Al Abosy, R., Freeman, G.J., et al. (2019). PTPN2 regulates the generation of exhausted CD8 T cell subpopulations and restrains tumor immunity. Nat Immunol 20, 1335–1347. 10.1038/s41590-019-0480-4.

63. Jansen, C.S., Prokhnevska, N., Master, V.A., Sanda, M.G., Carlisle, J.W., Bilen, M.A., Cardenas, M., Wilkinson, S., Lake, R., Sowalsky, A.G., et al. (2019). An intra-tumoral niche maintains and differentiates stem-like CD8 T cells. Nature 576, 465–470. 10.1038/s41586-019-1836-5.

64. Tillé, L., Cropp, D., Charmoy, M., Reichenbach, P., Andreatta, M., Wyss, T., Bodley, G., Crespo, I., Nassiri, S., Lourenco, J., et al. (2023). Activation of the transcription factor NFAT5 in the tumor microenvironment enforces CD8^+^ T cell exhaustion. Nat Immunol 24, 1645–1653. 10.1038/s41590-023-01614-x.

65. Khan, O., Giles, J.R., McDonald, S., Manne, S., Ngiow, S.F., Patel, K.P., Werner, M.T., Huang, A.C., Alexander, K.A., Wu, J.E., et al. (2019). TOX transcriptionally and epigenetically programs CD8^+^ T cell exhaustion. Nature 571, 211–218. 10.1038/s41586-019-1325-x.

66. Scott, A.C., Dündar, F., Zumbo, P., Chandran, S.S., Klebanoff, C.A., Shakiba, M., Trivedi, P., Menocal, L., Appleby, H., Camara, S., et al. (2019). TOX is a critical regulator of tumour-specific T cell differentiation. Nature 571, 270–274. 10.1038/s41586-019-1324-y.

67. Alfei, F., Kanev, K., Hofmann, M., Wu, M., Ghoneim, H.E., Roelli, P., Utzschneider, D.T., von Hoesslin, M., Cullen, J.G., Fan, Y., et al. (2019). TOX reinforces the phenotype and longevity of exhausted T cells in chronic viral infection. Nature 571, 265–269. 10.1038/s41586-019-1326-9.

68. Chen, Z., Ji, Z., Ngiow, S.F., Manne, S., Cai, Z., Huang, A.C., Johnson, J., Staupe, R.P., Bengsch, B., Xu, C., et al. (2019). TCF-1-Centered Transcriptional Network Drives an Effector versus Exhausted CD8 T Cell-Fate Decision. Immunity 51, 840–855.e845. 10.1016/j.immuni.2019.09.013.

69. Li, H., van der Leun, A.M., Yofe, I., Lubling, Y., Gelbard-Solodkin, D., van Akkooi, A.C.J., van den Braber, M., Rozeman, E.A., Haanen, J.B.A.G., Blank, C.U., et al. (2020). Dysfunctional CD8 T Cells Form a Proliferative, Dynamically Regulated Compartment within Human Melanoma. Cell 181, 747. 10.1016/j.cell.2020.04.017.

70. Wu, J.E., Manne, S., Ngiow, S.F., Baxter, A.E., Huang, H., Freilich, E., Clark, M.L., Lee, J.H., Chen, Z., Khan, O., et al. (2023). In vitro modeling of CD8^+^ T cell exhaustion enables CRISPR screening to reveal a role for BHLHE40. Sci Immunol 8, eade3369. 10.1126/sciimmunol.ade3369.

71. Schietinger, A., Philip, M., Krisnawan, V.E., Chiu, E.Y., Delrow, J.J., Basom, R.S., Lauer, P., Brockstedt, D.G., Knoblaugh, S.E., Hämmerling, G.J., et al. (2016). Tumor-Specific T Cell Dysfunction Is a Dynamic Antigen-Driven Differentiation Program Initiated Early during Tumorigenesis. Immunity 45, 389–401. 10.1016/j.immuni.2016.07.011.

72. Scharping, N.E., Rivadeneira, D.B., Menk, A.V., Vignali, P.D.A., Ford, B.R., Rittenhouse, N.L., Peralta, R., Wang, Y., DePeaux, K., Poholek, A.C., and Delgoffe, G.M. (2021). Mitochondrial stress induced by continuous stimulation under hypoxia rapidly drives T cell exhaustion. Nat Immunol 22, 205–215. 10.1038/s41590-020-00834-9.

73. Voskoboinik, I., Whisstock, J.C., and Trapani, J.A. (2015). Perforin and granzymes: function, dysfunction and human pathology. Nat Rev Immunol 15, 388–400. 10.1038/nri3839.

74. Paley, M.A., Kroy, D.C., Odorizzi, P.M., Johnnidis, J.B., Dolfi, D.V., Barnett, B.E., Bikoff, E.K., Robertson, E.J., Lauer, G.M., Reiner, S.L., and Wherry, E.J. (2012). Progenitor and terminal subsets of CD8^+^ T cells cooperate to contain chronic viral infection. Science 338, 1220–1225. 10.1126/science.1229620.

75. Sade-Feldman, M., Yizhak, K., Bjorgaard, S.L., Ray, J.P., de Boer, C.G., Jenkins, R.W., Lieb, D.J., Chen, J.H., Frederick, D.T., Barzily-Rokni, M., et al. (2018). Defining T Cell States Associated with Response to Checkpoint Immunotherapy in Melanoma. Cell 175, 998–1013.e1020. 10.1016/j.cell.2018.10.038.

76. Kovacs, S.B., Oh, C., Aachoui, Y., and Miao, E.A. (2021). Evaluating cytokine production by flow cytometry using brefeldin A in mice. STAR Protoc 2, 100244. 10.1016/j.xpro.2020.100244.

77. Nahmad, A.D., Reuveni, E., Goldschmidt, E., Tenne, T., Liberman, M., Horovitz-Fried, M., Khosravi, R., Kobo, H., Reinstein, E., Madi, A., et al. (2022). Frequent aneuploidy in primary human T cells after CRISPR-Cas9 cleavage. Nat Biotechnol 40, 1807–1813. 10.1038/s41587-022-01377-0.

78. Ma, A., and Malynn, B.A. (2012). A20: linking a complex regulator of ubiquitylation to immunity and human disease. Nat Rev Immunol 12, 774–785. 10.1038/nri3313.

79. Glaser, V., Flugel, C., Kath, J., Du, W., Drosdek, V., Franke, C., Stein, M., Pruß, A., Schmueck-Henneresse, M., Volk, H.D., et al. (2023). Combining different CRISPR nucleases for simultaneous knock-in and base editing prevents translocations in multiplex-edited CAR T cells. Genome Biol 24, 89. 10.1186/s13059-023-02928-7.

80. Tsuchida, C.A., Brandes, N., Bueno, R., Trinidad, M., Mazumder, T., Yu, B., Hwang, B., Chang, C., Liu, J., Sun, Y., et al. (2023). Mitigation of chromosome loss in clinical CRISPR-Cas9-engineered T cells. Cell 186, 4567–4582.e4520. 10.1016/j.cell.2023.08.041.

81. Chiesa, R., Georgiadis, C., Syed, F., Zhan, H., Etuk, A., Gkazi, S.A., Preece, R., Ottaviano, G., Braybrook, T., Chu, J., et al. (2023). Base-Edited CAR7 T Cells for Relapsed T-Cell Acute Lymphoblastic Leukemia. N Engl J Med 389, 899–910. 10.1056/NEJMoa2300709.

82. Chen, Y., Ye, Z., Chen, L., Qin, T., Seidler, U., Tian, D., and Xiao, F. (2020). Association of Clinical Phenotypes in Haploinsufficiency A20 (HA20) With Disrupted Domains of A20. Front Immunol 11, 574992. 10.3389/fimmu.2020.574992.

83. Garcia, J., Daniels, J., Lee, Y., Zhu, I., Cheng, K., Liu, Q., Goodman, D., Burnett, C., Law, C., Thienpont, C., et al. (2024). Naturally occurring T cell mutations enhance engineered T cell therapies. Nature 626, 626–634. 10.1038/s41586-024-07018-7.

84. Kato, M., Sanada, M., Kato, I., Sato, Y., Takita, J., Takeuchi, K., Niwa, A., Chen, Y., Nakazaki, K., Nomoto, J., et al. (2009). Frequent inactivation of A20 in B-cell lymphomas. Nature 459, 712–716. 10.1038/nature07969.

85. Braun, F.C., Grabarczyk, P., Möbs, M., Braun, F.K., Eberle, J., Beyer, M., Sterry, W., Busse, F., Schröder, J., Delin, M., et al. (2011). Tumor suppressor TNFAIP3 (A20) is frequently deleted in Sézary syndrome. Leukemia 25, 1494–1501. 10.1038/leu.2011.101.

86. Giles, J.R., Ngiow, S.F., Manne, S., Baxter, A.E., Khan, O., Wang, P., Staupe, R., Abdel-Hakeem, M.S., Huang, H., Mathew, D., et al. (2022). Shared and distinct biological circuits in effector, memory and exhausted CD8+ T cells revealed by temporal single-cell transcriptomics and epigenetics. Nat Immunol 23, 1600–1613. 10.1038/s41590-022-01338-4.

87. Daniel, B., Yost, K.E., Hsiung, S., Sandor, K., Xia, Y., Qi, Y., Hiam-Galvez, K.J., Black, M., J Raposo, C., Shi, Q., et al. (2022). Divergent clonal differentiation trajectories of T cell exhaustion. Nat Immunol 23, 1614–1627. 10.1038/s41590-022-01337-5.

88. Dou, Z., Bonacci, T.R., Shou, P., Landoni, E., Woodcock, M.G., Sun, C., Savoldo, B., Herring, L.E., Emanuele, M.J., Song, F., et al. (2024). 4-1BB-encoding CAR causes cell death via sequestration of the ubiquitin-modifying enzyme A20. Cell Mol Immunol 21, 905–917. 10.1038/s41423-024-01198-y.

89. Pores-Fernando, A.T., and Zweifach, A. (2009). Calcium influx and signaling in cytotoxic T-lymphocyte lytic granule exocytosis. Immunol Rev 231, 160–173. 10.1111/j.1600-065X.2009.00809.x.

90. Dong, M.B., Wang, G., Chow, R.D., Ye, L., Zhu, L., Dai, X., Park, J.J., Kim, H.R., Errami, Y., Guzman, C.D., et al. (2019). Systematic Immunotherapy Target Discovery Using Genome-Scale In Vivo CRISPR Screens in CD8 T Cells. Cell 178, 1189–1204.e1123. 10.1016/j.cell.2019.07.044.

91. Brentjens, R.J., Santos, E., Nikhamin, Y., Yeh, R., Matsushita, M., La Perle, K., Quintás-Cardama, A., Larson, S.M., and Sadelain, M. (2007). Genetically targeted T cells eradicate systemic acute lymphoblastic leukemia xenografts. Clin Cancer Res 13, 5426–5435. 10.1158/1078-0432.CCR-07-0674.

92. Yang, J., Sanderson, N.S., Wawrowsky, K., Puntel, M., Castro, M.G., and Lowenstein, P.R. (2010). Kupfer-type immunological synapse characteristics do not predict anti-brain tumor cytolytic T-cell function in vivo. Proc Natl Acad Sci U S A 107, 4716–4721. 10.1073/pnas.0911587107.

93. Robbins, P.F., Li, Y.F., El-Gamil, M., Zhao, Y., Wargo, J.A., Zheng, Z., Xu, H., Morgan, R.A., Feldman, S.A., Johnson, L.A., et al. (2008). Single and dual amino acid substitutions in TCR CDRs can enhance antigen-specific T cell functions. J Immunol 180, 6116–6131. 10.4049/jimmunol.180.9.6116.

94. Jutz, S., Leitner, J., Schmetterer, K., Doel-Perez, I., Majdic, O., Grabmeier-Pfistershammer, K., Paster, W., Huppa, J.B., and Steinberger, P. (2016). Assessment of costimulation and coinhibition in a triple parameter T cell reporter line: Simultaneous measurement of NF-κB, NFAT and AP-1. J Immunol Methods 430, 10–20. 10.1016/j.jim.2016.01.007.

95. Chang, C.R., Vykunta, V.S., Lee, J.H.J., Li, K., Kochendoerfer, C., Muldoon, J.J., Wang, C.H., Mazumder, T., Sun, Y., Goodman, D.B., et al. (2025). SEED-Selection enables high-efficiency enrichment of primary T cells edited at multiple loci. Nat Biotechnol. 10.1038/s41587-024-02531-6.

